# Long-read direct RNA sequencing reveals epigenetic regulation of chimeric gene-transposon transcripts in *Arabidopsis thaliana*

**DOI:** 10.1101/2022.09.21.507229

**Authors:** Jérémy Berthelier, Leonardo Furci, Shuta Asai, Munissa Sadykova, Tomoe Shimazaki, Ken Shirasu, Hidetoshi Saze

## Abstract

Transposable elements (TEs) are accumulated in both intergenic and intragenic regions in plant genomes. Intragenic TEs often act as regulatory elements of associated genes and are also co-transcribed with genes, generating chimeric TE-gene transcripts. Despite the potential impact on mRNA regulation and gene function, the prevalence and transcriptional regulation of TE-gene transcripts are poorly understood. By long-read direct RNA sequencing and a dedicated bioinformatics pipeline, “ParasiTE”, we investigated the transcription and RNA processing of TE-gene transcripts in *Arabidopsis thaliana*. We identified a global production of TE-gene transcripts in thousands of *A. thaliana* gene loci, with TE sequences often being associated with alternative transcription start sites or transcription termination sites. The epigenetic state of intragenic TEs affects RNAPII elongation and usage of alternative poly(A) signals within TE sequences, regulating alternative TE-gene isoform production. Co-transcription and inclusion of TE-derived sequences into gene transcripts impact regulation of RNA stability and environmental responses of some loci. Our study provides novel insights into TE-gene interactions that contributes to mRNA regulation, transcriptome diversity, and environmental responses in plants.

## Introduction

Transposable elements (TEs) are major components of eukaryotic genomes, influencing the organization of chromosome structure and expansion of genome size^1,2^. TEs can insert into intragenic regions of gene units, including promoter, 5’/3’-untranslated regions (UTRs), exons, and introns, impacting both genetic and epigenetic regulation of gene transcription and transcriptome diversity^3^. For example, TEs can act as regulatory elements of nearby genes in response to environmental stresses^4–6^. Co-transcription and exonization of TE sequences generate chimeric TE-gene transcripts (TE-Gts), often resulting in the co-option of TE sequences for innovation of novel gene products^7^. In the human genome, it has been reported that 4% of proteincoding genes possess TE-derived exon sequences and that 5% of alternatively spliced internal exons are derived from short nuclear interspersed element *Alu*^8–10^. In contrast, it has been estimated that in *Arabidopsis*, TE sequences are associated with 7.8 % of expressed genes, and 1.2 % may contribute to the protein coding regions^11^.

Intragenic TE sequences can profoundly impact pre- and post-transcriptional regulation of mRNA^12^. TE sequences affect various alternative splicing (AS) processes of precursor mRNAs, resulting in intron retention (IR), exon skipping (ES), or the creation of alternative splice donor (or alternative 5’ splicing site; A5SS) and alternative splice acceptor sites (or alternative 3’ splicing site; A3SS)^13^. In addition, the internal promoter of a TE sequence can act as an alternative transcription start site (ATSS), while TE sequences in introns or 3’-UTRs can create alternative transcription termination sites (ATTS) owing to the presence of alternative polyadenylation (APA) signals within TE sequences^14^. In *Arabidopsis*, ~40% of intron-containing genes are alternatively spliced^15^, and tissue-specific and stress-responsive IR and ES events have been detected in a subset of loci^16^.

To suppress the harmful effects of active TEs, plants and other organisms have evolved epigenetic mechanisms, such as DNA methylation and histone modifications^17^. In plants, DNA is methylated at cytosines in CG and non-CG contexts (CHG and CHH, where H can be A, T, C or G). CG methylation is maintained by METHYLTRANSFERASE 1 (MET1), while non-CG methylation is in part regulated by the histone H3K9 methylase KRYPTONITE/SUVH4 together with SUVH5 and SUVH6^17^. The chromatin remodeler DECREASED DNA METHYLATION 1 (DDM1) is required for the maintenance of both DNA methylation and H3K9 methylation^18^. On the other hand, ectopic accumulation of H3K9 methylation and non-CG methylation in genic regions are prevented by the H3K9 demethylase INCREASE IN BONSAI METHYLATION 1 (IBM1)^19^.

In mammals, changes in DNA methylation and histone modification induce spurious transcription initiation from ATSSs in TE sequences^20,21^. In addition, a range of cancer cell types overexpress TE-derived alternative splice variants of oncogenes^22^. In plants, several transcription start sites (TSSs) of maize genes have been identified within TEs^6^, and in *Arabidopsis thaliana*, epigenetically regulated cryptic TSSs are mostly embedded TE sequences^23^. DNA methylation of intronic TEs has been shown to affect the mRNA splicing of genes that regulate the seed coat of soybean and the alternative poly(A) sites required for fruit development of oil palm^24,25^. Previous studies have shown that intronic TEs tend to be accumulated in plant disease resistance (*R*) genes, associated with repressive chromatin marks, including DNA methylation and H3K9 methylation, in the *Arabidopsis* and rice genomes^26,27^. These heterochromatic intragenic TEs are regulated by a protein complex that comprises ENHANCED DOWNY MILDEW 2 (EDM2), INCREASE IN BONSAI METHYLATION 2 (IBM2/ASI/SI), and ASI1-IMMUNOPRECIPITATED PROTEIN 1 (AIPP1)^28,29,27,30–33^. On the other hand, TEs inserted in the 3’-UTR regions of genes often regulate mRNA stability, translation, and subcellular localization^34–37^.

Despite the potential importance of TE-Gts in plant developmental traits and environmental responses, limitations of short-read RNA sequencing and a lack of bioinformatic pipelines have hindered the detection and comprehensive analyses of these transcripts in plants. The recent development of long-read sequencing technologies now permits in-depth analyses of the complexity of mRNA transcription and processing dynamics^38–43^, and transposon regulation^44^. In this study, we employed Oxford Nanopore Direct RNA Sequencing (ONT-DRS) technology to investigate the prevalence of TE-Gts in the *A. thaliana* transcriptome. By developing a new bioinformatic tool, ParasiTE, we identified TE-Gts associated with AS, ATSS, and ATTS. Additionally, we investigated the epigenetic and environmental regulation of TE-Gts using a mutant panel and public transcriptome dataset. Our study revealed a global production of TE-Gts generated from about 3,000 gene loci in the *Arabidopsis* transcriptome, with many TEs associated with ATSS and ATTS being found in 5’/3’-UTRs. We also demonstrate that TE sequences in 3’- UTRs contribute to the response of genes response to environmental signals and regulation of RNA stability.

## Results

### DRS of the *Arabidopsis* transcriptome

To understand the impact of TEs on RNA processing and the prevalence of chimeric TE-Gt formation in the *Arabidopsis* transcriptome, we exploited ONT-DRS which allows for native long-transcript sequencing^45^. We performed DRS with two biological replicates of wild-type Columbia (Col-0) seedlings and obtained over 1.2 million raw reads for each replicate (Supplementary Data 1). DRS could capture long TE-gene chimeric transcript isoforms, such as *Resistance to Peronospora parasitica 4* (*RPP4*)-*ATCOPIA4* in wild-type^46,32,42^, which have not been represented in the precedent annotations of *Arabidopsis* transcript isoforms, such as Araport11^47^ or the latest comprehensive *Arabidopsis* transcriptome dataset, AtRTD3^43^ (Supplementary Fig. 1a). However, the low coverage of the DRS dataset may hinder the comprehensive detection of potential TE-Gts. To circumvent the issue, we combined publicly available wild-type Col-0 DRS data^39^ with our wild-type Col-0 DRS data to increase read coverage for the detection of TE-Gts (Supplementary Table 1). After base corrections using paired-end Illumina short reads, the DRS transcriptome was assembled by Stringtie2^48^ (Supplementary Fig. 1b) (see Methods for details). The DRS dataset was further merged with either the Araport11 or AtRTD3 transcript dataset, yielding the unique dataset of transcriptome annotation datasets DRS-Araport11 and DRS-AtRTD3 (Fig. 1a; Supplementary Fig. 1b, Supplementary Table 2; data available at https://plantepigenetics.oist.jp/). DRS-Araport11 and DRS-AtRTD3 covered about 20% more genes with TE-Gts than the original Araport11 and AtRTD3 datasets (Supplementary Table 3). We mainly employed the DRS-AtRTD3 transcriptome dataset for downstream analyses since it showed a higher number of TE-Gts than DRS-Araport11 (Supplementary Fig. 1c, Supplementary Table 3).

**Figure 1.**
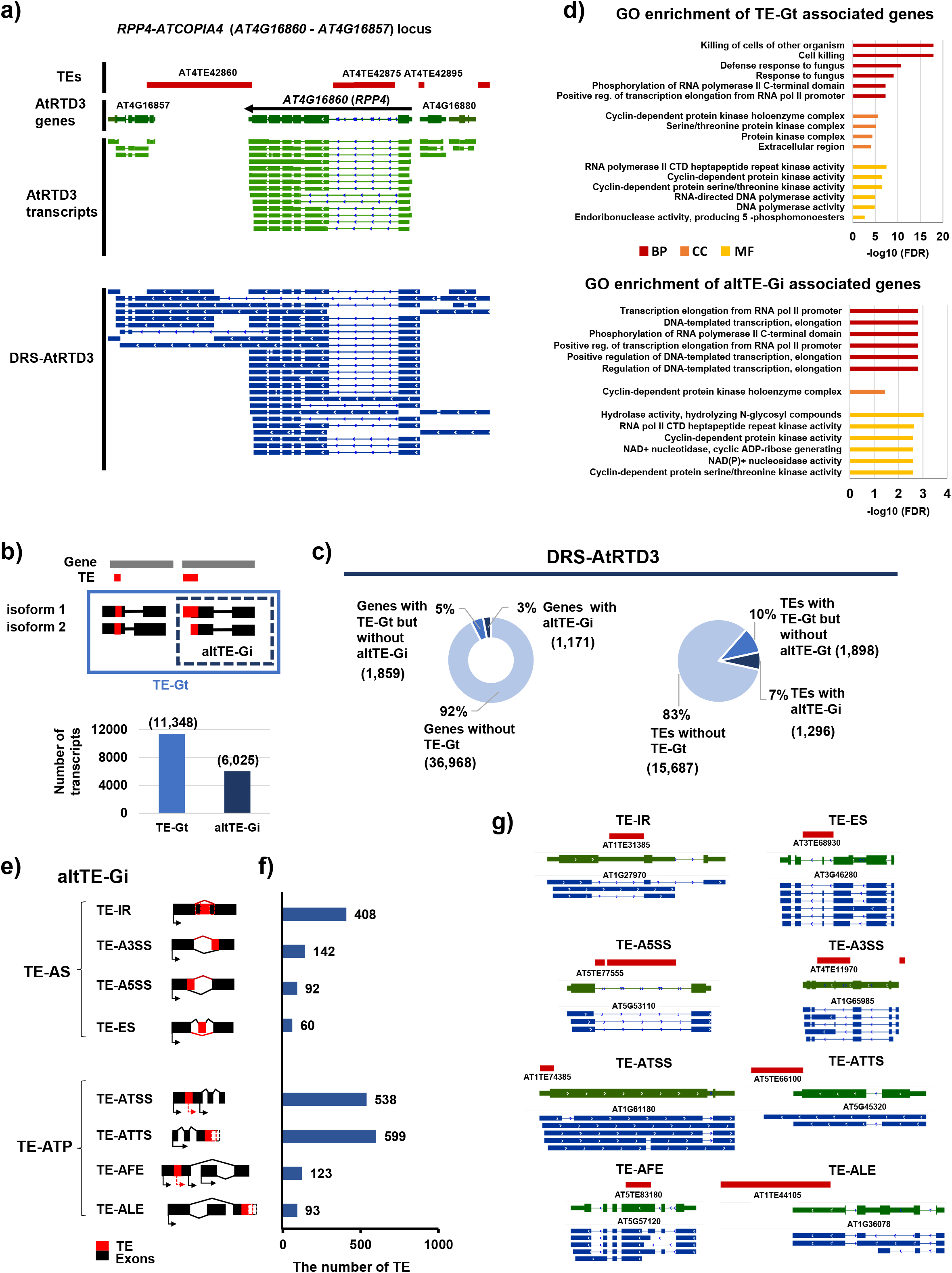
Detection of TE-gene transcripts (TE-Gts) and alternative TE-gene isoforms (altTE-Gis) in DRS-AtRTD3 of the *Arabidopsis* transcriptome. **a)** A representative TE-Gt associated with the *RPP4-ATCOPIA4* locus detected in DRS-AtRTD3. **b)** The number of TE-Gts and altTE-Gis detected by ParasiTE. **c)** The number of genes (left) and TEs (right) associated with TE-Gts and altTE-Gis identified in DRS-AtRTD3. **d)** Enriched Gene Ontology terms of genes associated with TE-Gts (top) and altTE-Gis in DRS-AtRTD3 (bottom). BP, Biological Process; CC, Cellular Component; MF, Molecular Function. **e)** RNA processing events associated with altTE-Gis (TE-AS and TE-ATP), and **f)** the number of TEs associated with altTE-Gi events in DRS-AtRTD3. Some TEs are included in several altTE-Gi categories. TE-AFE and TE-ALE are included in TE-ATSS and TE-ATTS, respectively. **g)** Representative loci associated with TE-AS and TE-ATP events detected in DRS-AtRTD3. Red, TE annotation; Green, AtRTD3 annotation (collapsed); Blue, DRS-AtRTD3 annotation (extended).

### ParasiTE: a tool for the detection of TE-Gts and their alternative isoforms

Although there are various bioinformatic tools for identifying alternative promoters of genes provided by TEs^49,50^, to our knowledge, no tools existed that had been specifically designed to detect TE-Gts and their alternative isoforms from transcriptome datasets according to the associated RNA processing events. Therefore, we developed a new tool named ParasiTE to detect TE-Gts in transcriptome datasets. The ParasiTE pipeline is composed of five main steps: 1) removal of gene-like TE annotation and associated transcripts, 2) classification of intragenic and intergenic TEs, 3) classification of intronic and exonic TEs, 4) annotation of TE-Gts, and 5) annotation of TE-Gts with alternative transcript isoforms (altTE-Gis; Fig. 1b). Details of each step and additional filtering steps are described in Supplementary Note 1. ParasiTE further classifies altTE-Gis into TE-associated alternative splicing (TE-AS) and TE-associated alternative transcript production (TE-ATP) according to the associated RNA processing events detected by the tool CATANA^51^. TE-AS events include TE-A5SS, TE-A3SS, TE-ES, and TE-IR. TE-ATP events include TE-ATSS and TE-ATTS, and ParasiTE further distinguishes TE-associated alternative first exon (TE-AFE) from TE-ATSS and TE-ALE TE-associated alternative last exon (TE-ALE) from TE-ATTS^51^ (Supplementary Note 1). Validation of ParasiTE with Araport11 transcript annotation and manual inspection of the output data demonstrated that ParasiTE could detect TE-ATP events with higher accuracy than TE-AS events (Supplementary Note 1).

### Identification and classification of TE-Gts in *A. thaliana*

Analysis of TE-Gts detected by ParasiTE with the DRS-AtRTD3 and DRS-Araport11 datasets revealed a total of 11,348 TE-Gt and 6,025 altTE-Gi events in DRS-AtRTD3 (Fig. 1b). We found that about 17% of *A. thaliana* TEs (n = 3,194 out of a curated 18,881 TE annotations) were involved in TE-Gt formation, and 7% of which were transcribed as altTE-Gis (Fig. 1c; Supplementary Table 2, 3). A similar proportion of TE-Gts and altTE-Gis was detected in the DRS-Araport11 dataset (Supplementary Fig. 1c, d). About 8% (n = 3,030) of genes (39,998 gene models based on DRS-AtRTD3 annotation (Supplementary Table 2); 27,628 genes were associated with Arabidopsis Genome Initiative [AGI] codes) were associated with TE-Gts, which was similar to a previously reported estimation (7.8%)^11^. Among the TE-Gts, we found that 3% (n = 1,171) of genes were involved in altTE-Gi production (Fig. 1c; Supplementary Table 2, 3). The Gene Ontology (GO) analysis of biological functions of the TE-Gts suggested an enrichment in terms related to plant defense responses against pathogens, such as “Cell killing” and “Defense response to fungus”, while altTE-Gis were implicated in Pol II-dependent RNA transcription (Fig. 1d).

ParasiTE can classify altTE-Gis according to the associated RNA processing events (Fig. 1e–g). Among the TE-AS, TE-IR (408 TEs) was more frequent than TE-A3SS (142 TEs), TE-A5SS (92 TEs), or TE-ES (60 TEs; Fig. 1f, g). Interestingly, we found more TEs involved in TE- TP events with TE-ATSS (538 TEs) and TE-ATTS (599 TEs) than TE-AS (Fig. 1f, g). A similar distribution of altTE-Gis was detected in the DRS-Araport11 and DRS-AtRTD3 datasets (Fig. 1f, g; Supplementary Fig. 1d). The high frequency of TE-ATP events may have resulted from the higher number of TEs located in 5’ or 3’ regions of genes compared with TEs in the gene bodies (Supplementary Fig. 2a). Indeed, using annotations of AtRTD3 (coding sequences [CDS] and 5’/3’-UTR predictions)^43^, we found that TE sequences associated with TE-Gts or altTE-Gis were mainly located in 3’-UTRs of genes (Supplementary Fig. 2b), while we estimated that TEs in 1% of TE-Gt and 3% of altTE-Gi events were associated with CDS of *Arabidopsis* genes (Supplementary Fig. 2b). These results demonstrate that ParasiTE can efficiently detect chimeric TE-Gts and that TE sequences nearby or within genes contribute as a source for transcriptome diversity in *Arabidopsis*.

### Contribution of TE superfamilies to TE-Gt production

ParasiTE detected 3,194 exonic and 441 intronic TEs in the DRS-AtRTD3 dataset (Supplementary Fig. 2c). Among them, Helitron (or DHH; Wicker’s code^52^) and Mutator (DTM) were abundant families, which were also enriched in TE-Gt and altTE-Gi events (Fig. 2a; Supplementary Data 3). Most of the TE sequences associated with altTE-Gis were short and truncated, while retrotransposons (Copia [RLC], Gypsy [RLG], and LINE [RIX]) superfamily sequences were relatively long when compared with other TE superfamilies (Fig. 2b). We then examined DNA methylation levels of intronic TEs, and TEs associated with TE-Gt and altTE-Gi events by employing a DNA methylation calling method with the Nanopore long-read DNA sequencing data^53^, which allows for the detection of inner DNA methylation of long, repetitive TE; this is generally difficult to analyze by short-read bisulfite sequencing (Supplementary Fig. 3a, b and Supplementary Data 4). We found significantly lower DNA methylation in TEs associated with TE-Gts and altTE-Gis compared with intronic TEs and those involved in TE-AS events (Fig. 2c). These results indicate that degenerated and hypomethylated TE sequences are preferentially transcribed and incorporated into TE-Gts.

**Figure 2.**
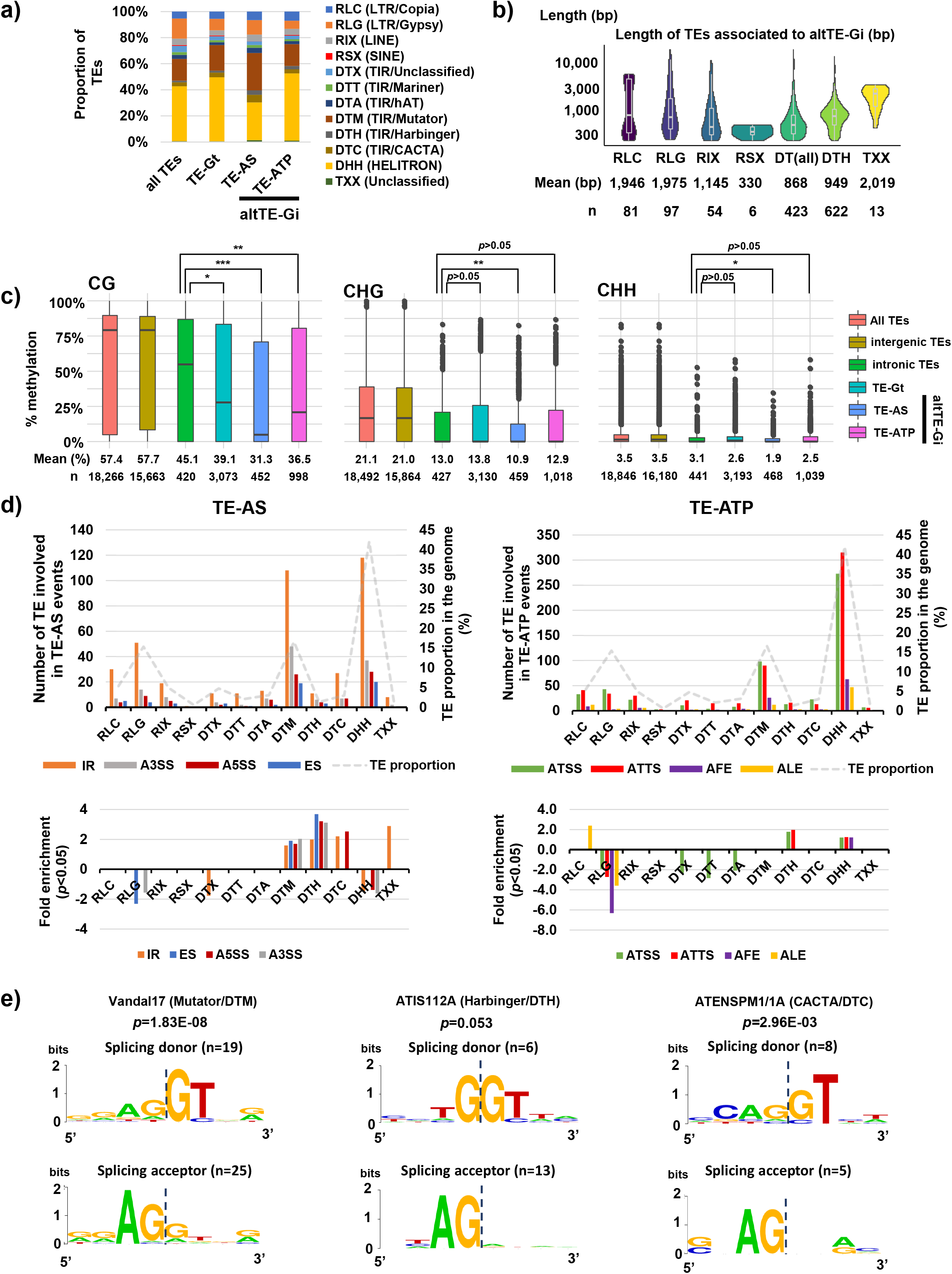
TE superfamilies associated with altTE-Gis. **a)** Proportion of TE superfamilies in the *Arabidopsis thaliana* genome, TE-Gt, and altTE-Gi. **b)** The length of TEs associated with altTE-Gis. **c)** DNA methylation (CG, CHG, and CHH contexts) of all TEs, intragenic TEs, intronic TEs, and TEs associated with TE-Gts, and with altTE-Gis. *P*-values were obtained by the Mann-Whitney U test; *, *p* < 0.05; **, *p* < 0.01; ***, *p* < 0.001. **d)** Top: the number of TEs associated with altTE-Gis (left, TE-AS; right, TE-ATP). The proportion of the TE superfamilies in the genome is also displayed as a reference. Bottom: Fold-enrichment of TE superfamilies significantly enriched in TE-AS or TE-ATP. *P*-values were obtained by the hypergeometric test. Only TE superfamilies with fold-enrichment of *p* < 0.05 are shown. **e)** Enrichment of nucleotides at splicing donor and splicing acceptor sites associated with TE superfamilies VANDAL17 (DTM), ATIS112A (DTH), and ATENSPM1/1A (DTC). *P*-values obtained by the hypergeometric test are indicated.

Intriguingly, we found significant over-enrichment of DNA transposons Mutator (DTM) and CACTA (DTC) superfamilies in TE-AS, including in TE-IR and TE-A5SS events (Fig. 2d; Supplementary Data 3). Further analysis of TE-IR events revealed enrichments of Mutator *VANDAL17* (DTM) and CACTA *ATENSPM1/1A* (DTC) at exon-intron junctions, providing canonical “GT-AG” sequences as splice donor/acceptor sites (Fig. 2e; Supplementary Data 3). The TE sequences involved in TE-IR were generally short (< 1,000 bp; Supplementary Fig. 3c), indicating that they are degenerated TE sequences retained in gene bodies. On the other hand, long terminal repeat LTR/Gypsy (RLG) was under-represented in most of the splicing events, while LTR/Copia (RLC) was significantly over-represented, especially in TE-ALE events (Fig. 2d). Overall, these results suggest that enrichment of particular TE superfamilies in intragenic regions contributes to splicing events of a subset of genes.

### Alteration of epigenetic modifications impacts TE-Gt production

TEs are often activated by changes in epigenetic modifications, which affects the expression of associated genes^23^. To understand the epigenetic regulation of TE-gene transcription, we analyzed altTE-Gi expression and isoform usage in mutants defective in the regulation of heterochromatic marks, including DNA methylation (*met1* and *ddm1*) and histone H3K9 methylation (H3K9m; *suvh456* and *ibm1*), and also mutants show defects in the transcription of intragenic TEs (*ibm2* and *edm2*). By analyzing changes in altTE-Gi usage in mutants using short-read RNA-seq datasets, we found that approximately 10–20 % of altTE-Gis showed significant differential isoform usage (or isoform switching) in the epigenetic mutants (Fig. 3a). This number was higher in *met1, ddm1*, and *suvh456*, which suggests that DNA methylation and H3K9m regulate transcription of altTE-Gis (Fig. 3a; Supplementary Data 5). We then performed de novo assembly of the DRS data based only on the mutant transcriptomes to identify mutant-specific altTE-Gis. DRS with three biological replicates yielded a total of 3.2–5.2 million reads for each mutant (Supplementary Data 1, 6). altTE-Gis in the mutant-DRS transcriptome assemblies detected by ParasiTE were further analyzed for differential isoform usage with short-read RNA-seq data, changes in DNA and H3K9 methylation, and association with the IBM2/EDM2 binding peaks obtained by chromatin immunoprecipitation (ChIP)-seq (Fig. 3a; Supplementary Fig. 4 and Supplementary Data 6). Due to the low coverage of the DRS data, a smaller number of altTE-Gis was identified in mutant-DRS transcriptome assemblies compared with DRS-AtRTD3 (Fig. 3a, b; Supplementary Fig. 4a). However, the *ddm1*-DRS transcriptome showed a higher number of altTE-Gis than the other mutants, suggesting that activation of TEs in this mutant may contribute to mutant-specific TE-Gt production (Epi-altTE-Gi). Indeed, in *ddm1* and in *met1*, we detected a previously reported TE-Gt in the *SQN (AT2G15790*) locus with TE-ATSS events, with activation of *AT2TE28020/25* (TIR/Mutator) in the upstream region inducing various readthrough transcripts with *SQN* (Fig. 3b)^54,23^. For *AT2G16050*, similar examples were found, with *AT2TE28420/25/30* (Helitron; Supplementary Fig. 5)^23^. In addition, we identified novel mutant-specific TE-ATSS at several gene loci, including *AT1G75990* with *AT1TE93320* (TIR/Mutator) and *AT3G23080* with *AT3TE34420/30* (TIR/hAT; Supplementary Fig. 5). We also identified a gene encoding singlestranded nucleotide-binding protein RH3 (*AT2G40960*) with *AT2TE77005/15* (LTR/Gypsy; while only 5’ and 3’ LTRs were annotated, we found that the region encodes a complete TE sequence belonging to the Gypsy-39_AT family; Fig. 3b, c). These TEs were hypomethylated in *ddm1* and/or *met1*, and based on the published cap analysis of gene expression (CAGE)-seq^23^, the altTE-Gis were transcribed from cryptic TSSs. These results demonstrate that epigenetic repression of ATSS associated with intergenic TEs, especially those located in the 5’ region of genes, is essential for the suppression of TE-Gt production with downstream genes.

**Figure 3.**
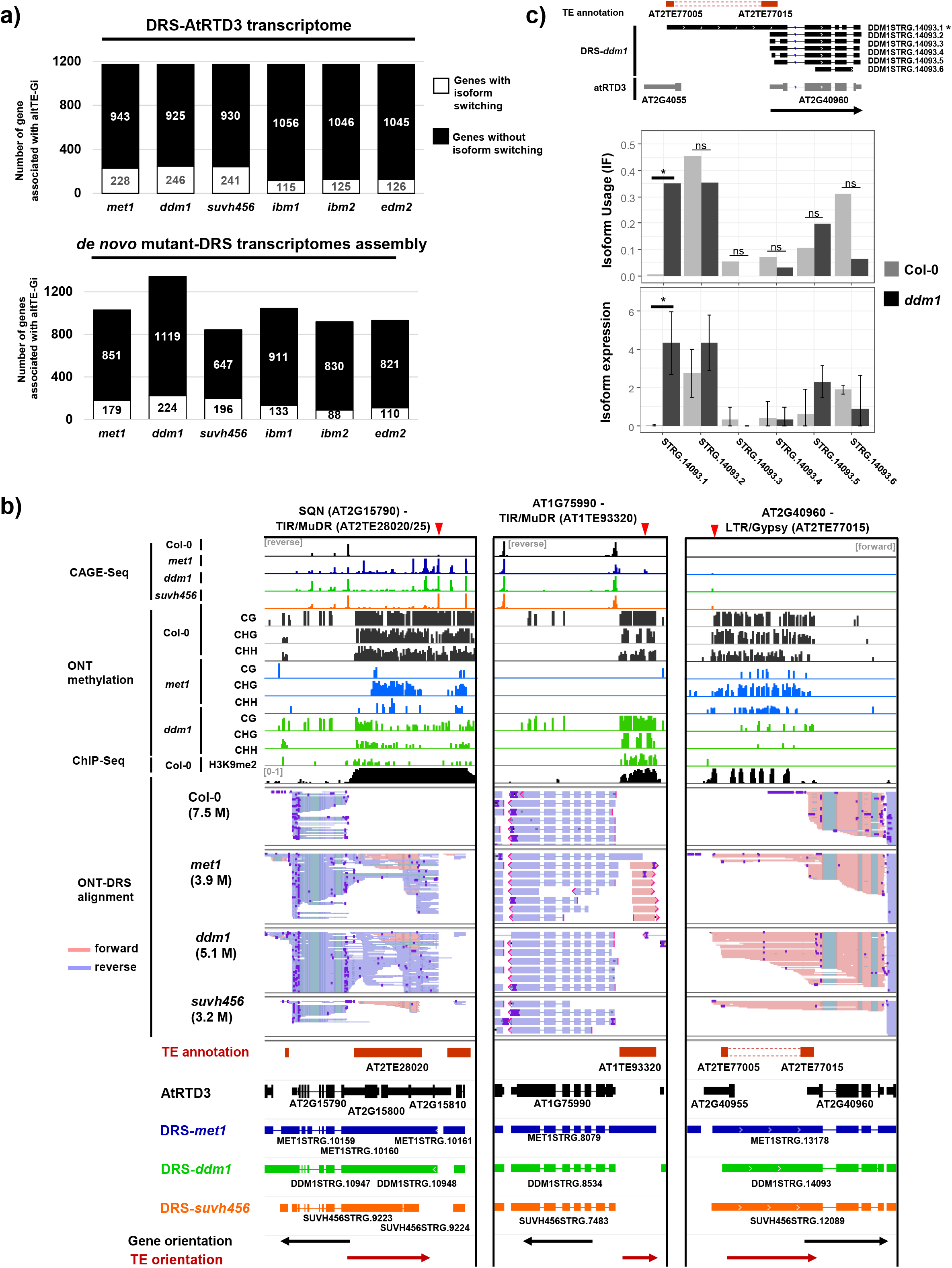
Epigenetic regulation of altTE-Gis. **a)** Top: Number of genes in each mutant showing isoform switching of altTE-Gis detected in DRS-AtRTD3. Bottom: Number of genes in each mutant showing change in isoform usage of altTE-Gis detected in mutant-DRS dataset. **b)** Representative genome loci showing epigenetically regulated altTE-Gi (Epi-altTE-Gi) production with TE-ATSS events. Tracks (from top to bottom): CAGE-seq (reads per million. Only forward or reverse strands are shown); Col-0 ChIP-seq of H3K9me2 (bin per million); methylation level of each mutant in CG, CHG, and CHH contexts (0–100%); DRS read alignments of Col-0 and indicated mutants; TE and AtRTD3 transcript annotations, *de novo* assembly of transcripts in mutants, and the orientation of genes and TEs. Red arrows on the top indicate cryptic TSSs detected in epigenetic mutants. **c)** Epi-altTE-Gi production and isoform switching detected at the *AT2G40960* locus in *ddm1*. Top: *ddm1*-DRS transcripts aligned to the *AT2G40960* locus. Middle: Isoform usage (IF) of altTE-Gis. Bottom: Expression levels of isoforms. Benjamini–Hochberg false discovery (FDR) corrected *p*-values (*q*-values; *, *q* < 0.05) for isoform switching and expression.

### Epigenetic regulation of TE-ATTS formation in *A. thaliana*

Together with TE-ATSS, TE-ATTS events were frequently detected in the transcriptome of DRS-AtRTD3 and of mutant-DRS (Fig. 1e, f; Supplementary Fig. 5, 6). Although TE sequences in promoter regions are known to act as regulatory elements of downstream genes in both animals and plants^2^, the impact of TEs inserted in the 3’ region of genes on transcription in *Arabidopsis* is less well understood. Based on the Col-0 DRS data, we detected poly(A) sites within 420/599 TEs associated with TE-ATTS events (Supplementary Fig. 6a), which may correspond to APA sites of associated genes. Indeed, 437/599 TEs associated with TE-ATTS events overlapped with poly(A) sites in the public APA data for the *Arabidopsis* genome^55^ (Supplementary Fig. 6a), suggesting frequent transcription termination at APA in TE sequences.

In the wild-type Col-0, TE-ATTS were formed as a result of readthrough transcription and termination at APA(s) in TEs inserted in the 3’ region of genes, including *ATCOPIA4* in *RPP4*, Gypsy/ATGP2 in *RPP5-like* (*AT4G16900*), *ONSEN/ATCOPIA78* in both *GEM-RELATED 5* (*GER5; AT5G13200*) and *BESTROPHIN-LIKE PROTEIN1* (*BEST1; AT3G61320*) loci (Fig. 4a; Supplementary Fig. 6–9). Despite active transcription of the TE regions, these TEs were often associated with repressive chromatin marks, such as DNA and H3K9 methylation, in the wildtype (Fig. 4a; Supplementary Fig. 6c, 8–11 and Supplementary Data 5, 6). TE-ATTS formation was also detected in intronic TEs in the epigenetic mutants, with the loss of heterochromatic marks in intronic TEs resulting in premature polyadenylation of mRNAs at promoter-proximal APAs within TE sequences^56,29,26^ of *R* gene *Resistance to Peronospora Parasitica 7* (*RPP7; AT1G58602*) and chloroplast protein gene *PPD7* (*AT3G05410*) (Supplementary Fig. 7, 12). In a few loci, however, we detected Epi-altTE-Gis showing rather enhanced readthrough transcription of 3′ TE sequences in the mutant backgrounds, such as *ibm1*, *edm2*, and *ibm2* (Supplementary Fig. 10, 11).

**Figure 4.**
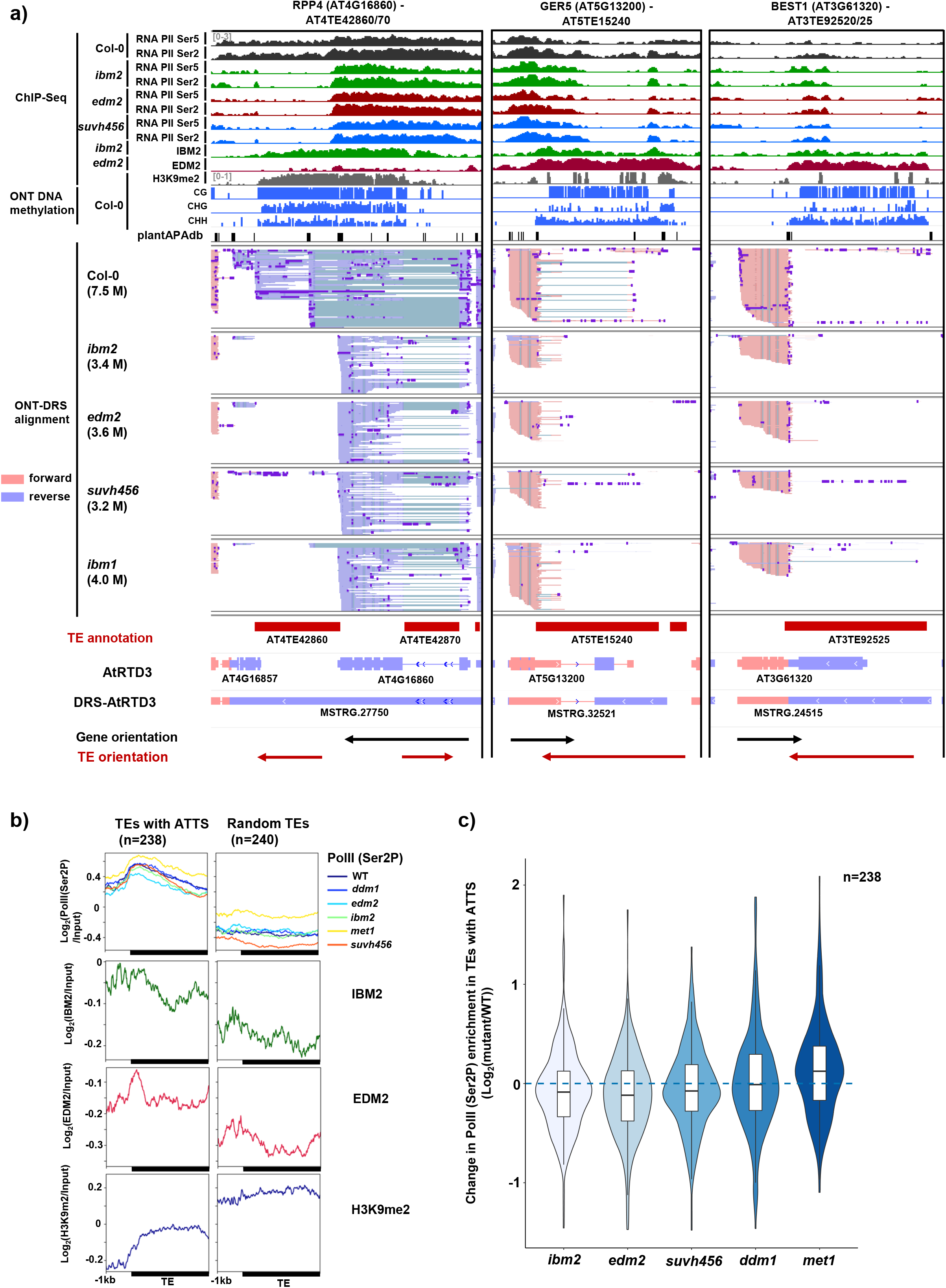
Epigenetic regulation of TE-ATTS. **a)** Representative genome loci showing Epi-altTE-Gi production with TE-ATTS events. Tracks (from top to bottom): ChIP-seq data for RNA Pol II phosphorylated at Ser5/Ser2 in CTD repeats (bins per million); ChIP-seq data for IBM2 and EDM2 localization (bins per million); Col-0 ChIP-seq of H3K9me2 (reads per million); methylation levels of Col-0 in CG, CHG, and CHH contexts (0–100%); poly(A) sites obtained from the PlantAPA database; DRS read alignments of Col-0 and indicated mutants; TE and transcript annotations of AtRTD3 and DRS-AtRTD3 in this study and the orientation of genes and TEs. **b)** Metaplots for ChIP-seq signals of Pol II (Ser2P), IBM2, EDM2, and H3K9me2 over TEs with ATTS (isoform switching with *q* < 0.05 detected at least once among mutants; Supplementary Data 5; n = 238) or randomly selected TEs (n = 240). **c)** Change in Pol II (Ser2P) ChIP-seq signals in TEs with ATTS (n = 238) between the wild-type and mutants.

Our ChIP-seq analysis of RNA Pol II phosphorylated at Ser5 and Ser2 of the carboxyterminal domain (CTD) repeats showed that in the wild-type Col-0, Pol II can elongate through intronic TEs or TEs in 3′ UTRs, despite the presence of repressive chromatin marks (Fig. 4a, b; Supplementary Fig. 6c, 12). *suvh456* caused a loss of Pol II peaks in the TE sequences (Fig. 4a, c; Supplementary Fig. 12), suggesting elongation defects and the release of Pol II from these TE regions. This result suggests that maintenance of heterochromatic modifications such as H3K9 and DNA methylation are required for Pol II elongation through the intragenic TE sequences in these loci. Mutants of IBM2 and EDM2 also showed similar premature polyadenylation and PolII elongation defects in TE regions (Fig. 4b; Supplementary Fig. 6b, S12), suggesting that rather than enhancing splicing of intragenic TE sequences, these factors likely suppress usage of proximal APAs and Pol II release during the transcription of heterochromatic TEs in intragenic regions. These results demonstrate that epigenetic mechanisms impact TE-ATTS formation by regulating Pol II elongation and usage of APA in intronic or 3’ TE sequences.

### The presence of TEs in 3’-UTRs contributes to environmental responses and RNA stability

To understand the regulatory role of 3’-UTR TEs, we further investigated representative gene loci containing LTR/Copia superfamily retrotransposons in their 3′-UTRs; this TE superfamily showed overrepresentation in the TE-ALE events in the *Arabidopsis* genome (Fig. 2d). *GER5* (*AT5G13200*) responds to abiotic stress and phytohormones including abscisic acid (ABA), and is involved in reproductive development and regulation of seed dormancy^57^. The *GER5* locus has an insertion of *ATCOPIA78/ONSEN* in the 3’-UTR in an antisense orientation (Fig. 4a, 5a; Supplementary Fig. 7 and 13a). In addition to altTE-Gi with TE-ATTS events, ONT-DRS also detected antisense transcripts of *GER5* promoted by ONSEN LTRs in Col-0 (Supplementary Fig. 13a). Quantitative PCR (qPCR) showed that *ibm2, edm2*, and *suvh456* caused transcription defects in the *ATCOPIA78* region and reduced transcription of the long altTE-Gi spanning the *ATCOPIA78* (*MSTRG.32521.2;* Fig. 5b, c; Supplementary Fig. 13b). Instead, the mutants showed increased expression of short altTE-Gis terminated at proximal APA sites in the TE sequences (such as *MSTRG. 32521.4*, and *.6;* Fig. 5c), showing an isoform switching of the altTE-Gis. ABA treatment induced higher expression of *GER5* as well as TE-Gts in wild-type Col-0 (Fig. 5b, c), while *ibm2, edm2*, and *suvh456* showed higher expression of short altTE-Gis (*MSTRG.32521 .4 and .6*) compared with Col-0 (Fig. 5c). Interestingly, *Arabidopsis* accessions with no *ATCOPIA78* insertion in the *GER5* locus (Ler-0, Kyoto, An-1, and Sha) showed enhanced induction of *GER5* by ABA treatment compared with accessions with TE insertions (Col-0, Lan-0, Pna-17; Fig. 5d), demonstrating that the presence of *ATCOPIA78* represses induction of *GER5* transcripts, especially upon ABA treatment. Variable expression of the altTE-Gis was observed in the TE-containing Col-0, Lan-0, and Pna-17 (Supplementary Fig. 13c), suggesting additional ecotype-specific regulation of the transcripts. Since the presence of the TE in the 3’-UTR may also affect the stability of *GER5* mRNA^37^, we examined its stability in Col-0 (TE+), Ler-0 (TE-), Sha (TE-), *ibm2*, *edm2*, and *suvh456* by treatment with the RNA synthesis inhibitor cordycepin (Fig. 5e; Supplementary Fig. 14a). We found that *GER5* mRNA stability in wild-type Col-0 was comparable to TE-less accessions Ler-0 and Sha (Fig. 5e). However, *ibm2, edm2*, and *suvh456* showed a rapid reduction of *GER5* transcript levels after cordycepin treatment, suggesting that defects in the transcription of the altTE-Gi spanning the *ATCOPIA78* resulted in *GER5* mRNA instability. Indeed, AU-rich elements (AREs)^58^ were prevalent in *GER5* transcripts terminated at proximal APAs in *ATCOPIA78* (Supplementary Fig. 14b), which may have led to enhanced degradation of *GER5* transcripts^37^. Thus, the presence of the TE in 3’-UTR of *GER5* contributes to responsiveness to ABA signaling as well as regulation of *GER5* mRNA stability.

**Figure 5.**
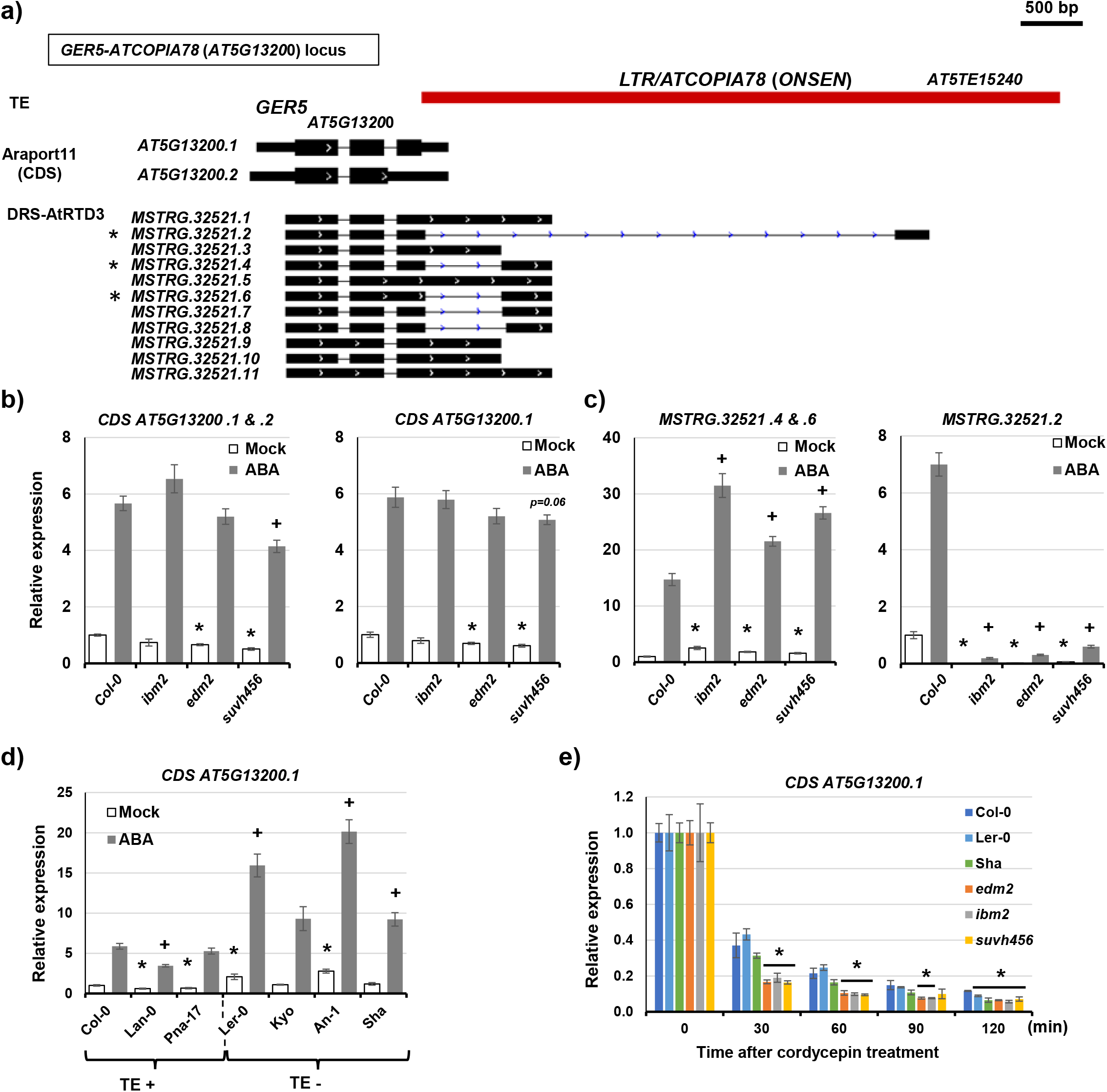
Epigenetic regulation of altTE-Gis in the *GER5* locus and the impact on environmental responses and RNA stability. **a)** *GER5-ATCOPIA78/ONSEN* locus. TEs, Araport11 gene annotation, and DRS-AtRTD3 transcript isoforms are shown. * indicates isoforms examined by RT-qPCR in **c**. **b)**, **c)** Relative expression of transcripts corresponding to *GER5* protein coding sequence (CDS; *AT5G13200.1* and *AT5G13200.2*) and altTE-Gi (*MSTRG.3521.2, 4, 6*) under mock and ABA stress conditions in indicated genotypes. Bars represent the means of four biological replicates ± standard error of the mean (SEM). *, *p* < 0.05 by *t*-test for comparison between Col-0 and mutants under mock conditions, and +, *p* < 0.05 by *t*-test under ABA stress conditions. **d)** Relative expression of transcripts corresponding to *GER5* CDS (*AT5G13200.1*) in the *A. thaliana* ecotypes with or without *ATCOPIA78/ONSEN* insertion in the 3’-UTR. Bars represent the means of four biological replicates ± SEM. *, *p* < 0.05 by *t*-test for comparison between Col-0 and mutants. **e)** Relative transcript levels of *GER5 (AT5G13200.2*) at 0, 30, 60, 90, and 120 min after cordycepin treatment in Col-0, *ibm2, edm2, suvh456*, and ecotypes without *ATCOPIA78/ONSEN* insertion (Ler-0 and Sha). Expression levels at 0 min are set as 1. Bars represent the means of four biological replicates ± SEM. *, *p* < 0.05 by *t*-test.

Similarly, we examined the impacts of TE insertion in the *RPP4-ATCOPIA4* locus (Fig. 4a, 6a). *RPP4* is an *R* gene that encodes nucleotide-binding and leucine-rich repeat domains (NLRs), which mediates the effector triggered immunity (ETI) response to isolates of the downy mildew oomycete *Hyaloperonospora arabidopsidis* (*Hpa*) Emwa1 and Emoy2 ^46,59^. In the *RPP4* locus of Col-0, *ATREP15* (Helitron) is inserted in the first intron, and *ATCOPIA4* (LTR/Copia) insertion in the sixth exon of the ancestral locus caused truncation of the 3’ part of the protein (Supplementary Fig. 7)^46^. In the Col-0 DRS data of this and a previous study^42^, *RPP4* transcript isoforms coding for at least seven open reading frames (ORFs) have been detected (Supplementary Fig. 15–17). Interestingly, despite the ~ 4.7 kb of the *ATCOPIA4* intragenic TE insertion, the *RPP4* locus still generates transcript isoforms that preserve the ORFs of the original exons in the 3’ regions (annotated as *AT4G16857, CDS E* in Supplementary Fig. 15), encoding a putative C-terminal jelly roll/Ig-like domain (C-JID; Supplementary Fig. 15,16). The C-JID domain might have a role in pathogen recognition of RPP4^60^, although RPP4 isoform without the C-JID region is sufficient for triggering ETI response^59^. Consistent with the previous reports^32,61^, epigenetic mutants *ibm2, edm2*, and *suvh456* caused a loss of altTE-Gis spanning *ATCOPIA4* (such as *MSTRG.27750.7;* Fig. 4a, 6a, b; Supplementary Fig. 17). Moreover, we found that altTE-Gi associated with the Helitron *ATREP15* (*MSTRG.27750.20*) was also downregulated in epigenetic mutants (Fig. 6a, b; R-P5). Similar to *GER5* transcripts, *RPP4* transcripts in *ibm2*, *edm2*, and *suvh456* showed a rapid reduction after cordycepin treatment (Fig. 6c), suggesting that the production of altTE-Gis spanning the *ATCOPIA4* may affect the stability of *RPP4* mRNA (Fig. 6c; Supplementary Fig. 14a, 18). We further investigated the impacts of the altTE-Gi production on the *RPP4* function by assessing the ETI response to *Hpa*^46,62^. *Arabidopsis* natural accessions and epigenetic mutants were inoculated with *Hpa* isolate *Emoy2*, which is recognized by *RPP4* and elicits downstream responses^46^. *Arabidopsis* accessions NFA-10 and Kas-2, which are known to be susceptible to the isolate^63^, showed strong infection symptoms as expected (Fig. 6d). Interestingly, *ibm2*, *edm2*, and *suvh456* showed increased ETI effectiveness against *Hpa* Emoy2 (Fig. 6d), which is in contrast to previous reports showing either a mild reduction in resistance or no differences in the mutants, respectively^32,61^. These differences in response may have originated from the tissues or isolates used in the study (true leaves in this study vs. cotyledon, for which has been reported age-related differences in resistance^64^; Emoy2 vs. Emwa1). To directly dissect the effects of TE insertion on the *RPP4-*mediated *Hpa* response, we also examined T-DNA insertion lines in the *ATCOPIA4*. The SALK_005767 line showed susceptibility to *Hpa* (Fig. 6a, d), consistent with a previous report^65^. Interestingly, however, the second T-DNA line (SALKseq_035375) inserted downstream of *ATCOPIA4* showed a resistant response (Fig. 6d). qPCR showed that the SALK T-DNA insertions caused differential altTE-Gi usage for *RPP4* under normal condition, such as different altTE-Gi expression for R-P1 and R-P4 (Fig. 6b), which may have contributed to the varying susceptibilities to *Hpa*. Although it is not clear how the T-DNA insertions in the TE sequence affect the overall *RPP4*-mediated pathogen response, the results suggest that the presence of the TE in the 3′ region of *RPP4* modulates the response to *Hpa* and the stability of *RPP4* transcripts.

**Figure 6.**
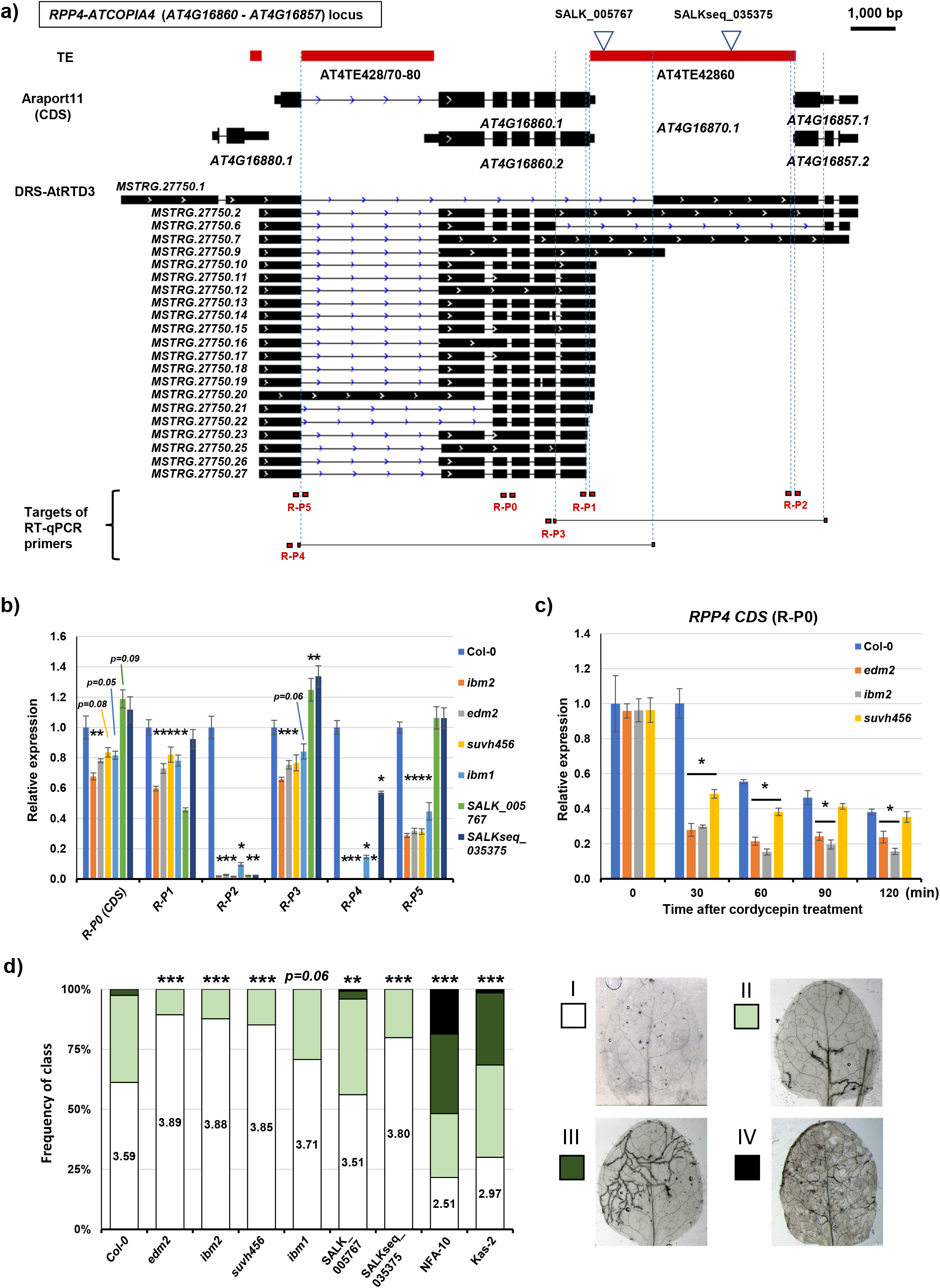
Epigenetic regulation of altTE-Gis in the *RPP4* locus and the impact on environmental responses and RNA stability. **a)** *RPP4-ATCOPIA4* locus. TEs, Araport11 gene annotation, DRS-AtRTD3 transcript isoforms, and primers for RT-qPCR are shown. **b)** Relative expression of *RPP4* transcripts detected by RT-qPCR with primers indicated in **a**. Bars represent the means of four biological replicates ± SEM. *, *p* < 0.05 by *t*-test. **c)** Relative transcript levels of *RPP4* at 0, 30, 60, 90, and 120 min after cordycepin treatment in Col-0, *ibm2, edm2*, and *suvh456*. Expression levels at 0 min are set as 1. Bars represent the means of four biological replicates ± SEM. *, *p* < 0.05 by *t*-test. **d)** Incompatibility of *A. thaliana* ecotypes and mutants against *Hyaloperonospora arabidopsidis* infection. NFA-10 and Kas-2 are ecotypes without the *RPP4* locus and were used as controls. Class I (white), hypersensitive response surrounding conidia penetration sites; class II (light green), presence of trailing necrosis in ≤50% leaf area; class III (dark green), presence of trailing necrosis in ≤75% leaf area; class IV (black), compromised ETI immunity, presence of pathogen hyphae not targeted by HR and conidiophores. Statistically significant differences in frequency distribution of the classes between lines and Col-0 were determined by Pearson’s chi-squared test; *, *p* < 0.05; **, *p* < 0.01; ***, *p* < 0.001. 70–130 leaves were analyzed per line across three separate experimental replicates.

### Environmental stresses affect alternative transcription termination by TE sequences

In addition to the epigenome modulation by mutants, we examined whether environmental stresses influence isoform switching and differential expression of altTE-Gis. Using public transcriptome data from biotic/abiotic stress treatment studies (ABA, methyl jasmonate, flagellin 22, salicylic acid, cold, heat, warm, salt, drought, and ultraviolet), we investigated changes in isoform usage and differential expression of representative altTE-Gis as well as mutant-specific Epi-altTE-Gis with TE-ATSS or TE-ATTS events (Fig. 3, 4; Supplementary Fig. 7–12). We found that Epi-altTE-Gis with TE-ATSS generated in a mutant background (*met1, ddm1*, and *suvh456*) were overall stably silenced under the stresses (Supplementary Fig. 19). In contrast, altTE-Gis and Epi-altTE-Gis with TE-ATTS detected in the epigenetic mutants (*met1, ddm1, suvh456, ibm1, ibm2*, and *edm2;* we also added the new RNA-seq dataset of *ibm12, ibm2-i.7*^29^, and *ibm2edm2*) showed changes in isoform usage and differential expression under various stresses, which often mirrored the responses of the epigenetic mutants (Fig. 6). Isoform switching and changes in expression manifested upon heat stress in gene loci with the heat-responsive *ATCOPIA78/ONSEN* insertion (Fig. 7; Supplementary Fig. 20a-f). These data suggest that environmental signals can regulate the transcription and processing of altTE-Gi via epigenetic regulation, which may contribute to an adaptive response to environmental changes.

**Figure 7.**
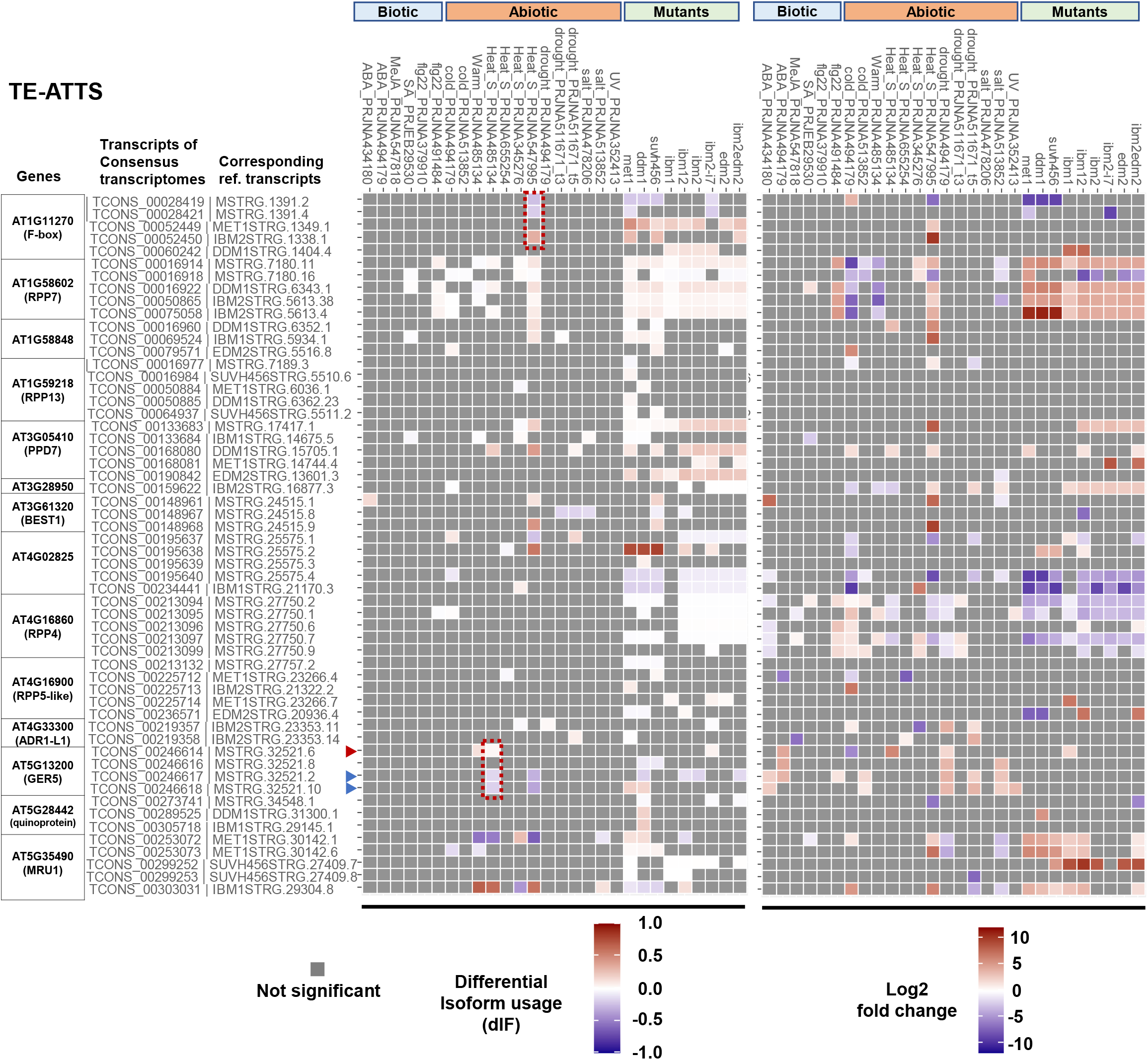
Regulation of Epi-altTE-Gi candidates under various stress conditions. Heatmaps showing statistically significant differential isoform usage (dIF; left) and differential expression fold-change (right) with TE-ATTS under stress conditions or in epigenetic mutants. Transcriptome data with the stress treatment studies are from the public RNA-seq data. The Epi-altTE-Gi candidates *AT1G58848* and *AT1G59218 (RPP13*) found by others were also added^32^. Red dotted lines highlight predicted isoform switching at *GER5 (AT5G13200*) and F-box gene (*AT1G11270*) loci in Col-0 under heat stress conditions. The names of the transcripts from the consensus DRS transcriptome as well as one corresponding to the reference transcript are indicated (more than one reference transcript from DRS-AtRTD3 or mutant-DRS are applicable to some transcripts).

## Discussion

In this study, we employed ONT-DRS technology to dissect the complexity of TE-Gts and their isoform production^38,39,41,42^. In addition, we developed the new bioinformatics tool ParasiTE to comprehensively detect TE-Gt isoforms present in the *Arabidopsis* transcriptome. To our knowledge, ParasiTE is the first tool capable of identifying chimeric TE-Gt isoforms and annotating associated RNA processing events (Fig. 1; Supplementary Note 1). The ParasiTE pipeline can be applied to a wide range of species using transcript, gene, and TE annotations as inputs and can be combined with short/long-read RNA-seq datasets to systemically identify TE-Gt isoforms under various experimental conditions.

We found that about 3,000 *Arabidopsis* genes are associated with TE-Gt production, corresponding to about 8% of protein-coding genes annotated in AtRTD3 (Fig. 1), which is close to an estimation made by a previous study^11^. In general, TEs tend to be short and degenerated in genic regions because of their deleterious effects on gene function^1,26^. Indeed, we found that TEs involved in altTE-Gis are shorter and less methylated than intergenic and intronic TEs (Fig. 2). Nevertheless, we still detected enrichment of DNA transposon superfamilies that provide canonical splice donor-acceptor sequences (Fig. 2d, e). In addition, we also identified a modest but significant enrichment of Helitron superfamily TEs associated with TE-ATSS/ATTS/AFE, and LTR/Copia superfamily TEs with TE-ALE events (Fig. 2d). In animals and plants, TEs can provide cis-regulatory sequences for expression of associated genes^66,67,2,34,6,23,68^. Thus, partial TE sequences could be co-opted as regulatory elements for transcriptional and post-transcriptional regulation of mRNA. On the other hand, young intact TE insertions are often polymorphic among natural accessions (Fig. 5)^69–72^, suggesting that intragenic TEs contribute to gene control for adaptation to the local environment and phenotypic diversity^73^.

Our previous study revealed that thousands of cryptic TSSs in TEs are suppressed by repressive epigenetic modifications^23^. Similarly, we found that usage of cryptic APA sites within TEs in introns or 3’-UTRs was suppressed by repressive epigenetic marks (Fig. 3–6). We detected Pol II elongation and transcription of long read-through RNAs over several kb of heterochromatic TEs (Fig.4; Supplementary Fig. 7–12), indicating that heterochromatin per se is more permissive for Pol II transcription than previously considered. This process can be mechanistically separated from the IBM2/EDM2/AIPP1 pathway that suppresses the usage of cryptic APA in heterochromatic TEs since in *ibm2/*edm2, Pol II enrichment at the gene body of the *RPP4* locus associated with heterochromatic marks was not greatly affected (Fig. 4b), while Pol II signals sharply decreased downstream of cryptic APA in the 3’-UTR of *AT4G16860* in the mutants (Fig. 4b; Supplementary Fig. 14). In *Drosophila*, the Rhino-Deadlock-Cutoff (RDC) complex licenses Pol II transcription of piRNA clusters marked by H3K9me3^74,75^. The Cutoff protein in the complex suppresses the usage of APA sites in TEs, suggesting conservation of a mechanism involving Pol II transcription of heterochromatic repeats^74,75^.

In this study, we also investigated how TE-Gts are regulated by environmental signals. We focused on the *GER5* locus harboring the insertion of *LTR/ATCOPIA78*(*ONSEN*) in the 3’-UTR (Fig. 5). *ONSEN* has heat-responsive elements in the LTR regions and is activated in response to heat stress via binding of heat-shock factors^76^. *ONSEN* insertions found in *GER5* and also *BEST1* (Supplementary Fig. S7) are considered mobile and are highly expressed copies under heat stress^70^. Indeed, heat-shock experiments induced isoform switching of altTE-Gis in *GER5* (Supplementary Fig. 20), showing that the *ONSEN* insertion confers heat responsiveness to the locus. *ONSEN* is known to introduce various transcriptional modulations to associated genes, including activation, AS, acquisition of heat responsiveness, and production of antisense RNAs^70,77^. Interestingly, we also found that the insertion of *ONSEN* causes a variable response of *GER5* transcription to ABA treatment in the natural accessions (Fig.5; Supplementary Fig. 13), which may be independent from the heat response and instead result from the instability of *GER5* mRNA caused by the TE sequence in the 3’-UTR (Fig. 4). TE sequences provide ARE motifs that may induce rapid degradation and turnover of the transcripts, as has been observed in *FLOWERING LOCUS C* mRNA in *Capsella*, which results in variations in flowering timing^37^. Whether the insertion of *ONSEN* in the *GER5* locus and production of altTE-Gis are beneficial for environmental response and local adaptation mechanisms remains unclear. We identified enhanced expression of the shorter form of *GER5* mRNA in epigenetic mutants, including *suvh456* (Fig. 5). Interestingly, a recent report showed that *GER5* is overexpressed with CHH hypomethylation in response to aphid feeding and is also constitutively activated in *suvh4*/*kyp* background, suggesting that *GER5* altTE-Gi may contribute to the defense response and resistance to aphids^78^.

We found functional enrichment of pathogen responses including “Cell killing” and “Defense response to fungus”, in genes associated with TE-Gt production (Fig.1d). In plants, the ETI response to microbial pathogens is mediated by *NLR* genes, which often form gene clusters containing repetitive sequences and TEs^79^. *NLR* gene clusters are the most genetically and epigenetically divergent loci in *Arabidopsis* populations^80^, and TEs contributed to the rapid evolution of *NLR* genes by enhancing recombination and exon shuffling^5^. We consistently found various altTE-Gi transcriptions in the *R* genes *RPP4*, *RPP5-like*, and *RPP7* (Fig. 4, 6, 7, Supplementary Fig. 6, 9). In these loci, TE sequences were often spliced out from mature mRNAs or introduced premature poly(A) sites, suggesting a minor contribution of TE sequences to the acquisition of novel protein-coding functions. In *Arabidopsis*, mRNA isoforms with premature termination codon are targeted for degradation by the nonsense-mediated mRNA decay (NMD) mechanism, which is often activated by stress-induced AS events^15,16^. Previous studies revealed that the NMD pathway targets NLR gene transcripts and is essential for tight control of immune receptor thresholds and suppression of auto-immune responses^81–83^. Thus, in addition to the epigenetic control of TEs surrounding NLR genes^81,56,84^, transcripts with TE sequences might provide a signal for post-transcriptional regulation of NLR gene transcripts. In this study, we further identified the production of TE-Gts at gene loci with various functions in response to environmental signals (Fig. 7), suggesting mechanistic roles of TE-Gt production in the regulation of mRNA processing and environmental responses.

## Methods

### Plant materials

*A. thaliana* mutants *met1-3, ddm1-1, ibm1-4, ibm2-2*, and *edm2-9* have been described previously^85,86,19,29,87^. *suvh456* seeds were kindly provided by Dr. Kakutani. All mutants were in the Col-0 background. The mutants *ibm1ibm2* and *ibm2edm2* were obtained by crossing *ibm1-4* with *ibm2-2* and *ibm2-2* with *edm2-9*, respectively. *ibm2-i7* is a transgenic line from the transformation of *ibm2-2* with the genomic DNA of *IBM1* without part of the sequence of the intron 7, as previously described^29^. Second generation homozygous *met1, ddm1, ibm1, ibm2, edm2, ibm1ibm2, and ibm2edm2* and T4 generation *ibm2-i7* were used for the RNA experiments described below. The *ibm2-2* transgenic line complemented with the genomic DNA of *IBM2* fused with the FLAG-HA tag was previously described^29^. A transgenic line expressing EDM2-Myc-HA was obtained by transforming *edm2-9* with the genomic DNA of *EDM2* fused with MYC-HA tag sequence. The seeds were germinated and grown on 1/2 Murashige and Skoog (MS) plates under long-day conditions (16 h light; 8 h dark) at 22°C.

### RNA extraction, Nanopore DRS and Illumina sequencing

For ONT-DRS (Oxford Nanopore Technologies, Oxford, UK) and Illumina short-read sequencing (Illumina Inc., San Diego, CA), 10-day-old whole seedlings of wild-type Col-0 and mutant plants were pooled for RNA extraction. Total RNA was extracted using RNAiso (TAKARA, Japan), and genome DNA was digested with TURBO DNase (Thermo Fisher Scientific, USA). For ONT-DRS, the Oligotex-dt30 <Super> mRNA purification kit (TAKARA) was used for poly(A) mRNA purification. Purified mRNA from wild-type Col-0, *met1, ddm1, suvh456, ibm1, ibm2*, and *edm2* were used to generate libraries using the Direct RNA Sequencing Kit (SQK-RNA002, Oxford Nanopore Technologies) according to the manufacturer’s instructions. Sequencing was performed using MinION device (device: MIN-101B, flow cells: FLO-MIN-106 R9 version; Oxford Nanopore Technologies) at the Okinawa Institute of Science and Technology Graduate University (OIST) Sequencing Center (SQC) or at our laboratory. We performed at least two sequencing runs for Col-0 and each mutant as biological replicates (Supplementary Data 1). To obtain a higher read depth for Col-0 ONT-DRS, previously generated public data for Col-0 ONT-DRS obtained by a standard ONT-DRS method (five runs) and by a 5’-cap capturing method (two runs) were combined with our Col-0 ONT-DRS data for detection of TE-Gts. Bases calling was performed with Guppy (v4.4.2.; https://nanoporetech.com/). Previously published short-read Illumina sequencing data (paired-end, 100 bp) for Col-0 and *ibm2*^26^, and new RNA-seq datasets for *ibm1*, *edm2, ibm2-i7* (described previously^29^)*, ibm1ibm2*, and *ibm2edm2* were obtained as described previously^26^. In addition, Illumina paired-end read data of *met1, ddm1, suvh456*, and Col-0 were retrieved from previous studies^29,23^. Illumina reads were trimmed using fastp (v0.21.0; parameters: -l 36 -r)^88^.

### Processing DRS sequencing data

For ONT-DRS data analysis of the Col-0 and epigenetics mutants, we first converted raw RNA sequence data into cDNA sequences with seqkit (v0.12.1; option: seq --rna2dna)^89^. ONT-DRS reads from replicates were concatenated and then error-corrected using LorDEC (v0.9; parameters: -k 21, -s 3)^90^ with the trimmed Illumina paired-end data of the corresponding genotype. Minimap2^91^ was used to align corrected ONT-DRS reads to the *Arabidopsis* genome (TAIR10) retrieved from The Arabidopsis Information Resource (TAIR) (https://www.arabidopsis.org/) with parameters: -ax splice -G 10k -uf -k14, as previously described^39^. The percentage of Araport11 genes covered by at least 10 reads of ONT-DRS or Illumina RNA-seq reads shown in Supplementary Figure 4a, was obtained using featureCount (v2.0.2; options: -L for ONT-DRS and -p for Illumina reads).

### ONT-DRS-based *de novo* transcriptome analysis

A *de novo* transcriptome of Col-0 was built using Stringtie2 (v2.1.4)^48^, as previously demonstrated^42,92^. We compared Stringtie2 “-R” or “-L” modes using either Araport11^47^ or AtRTD3^43^ annotations as references (option -G). The “-R” mode promoted a higher number of isoforms and was chosen to create the Col-0 DRS transcriptome with either Araport11 or AtRTD3 as references (Supplementary Fig. 1b). In order to improve annotation of *A. thaliana* to facilitate identification of altTE-Gis, we merged the two DRS transcriptomes (using Araport11 or AtRTD3 as a reference) with the corresponding annotations of Araport11 or AtRTD3 dataset using Stringtie2 (parameters: --merge -F 0 -T 0 -f 0 -g 1 -i -G) to generate DRS-Araport11 and DRS-AtRTD3 transcriptome annotations (Supplementary Fig. 1b). The *de novo* transcriptomes of the epigenetics mutants were built using Stringtie2 (v2.1.4) with “-R” mode and the AtRTD3^43^ annotation as the references (option -G).

### ParasiTE: a new tool for prediction of TE-Gts

To detect TE-Gt and altTE-Gi productions, we developed a new tool named ParasiTE. A detailed description of the pipeline can be found in Supplementary Note 1. ParasiTE can be downloaded at (https://github.com/JBerthelier/ParasiTE).

### DNA extraction and Nanopore DNA sequencing

Ten-day-old whole seedlings of wild-type Col-0, *met1*, and *ddm1* were pooled for DNA extraction. Extraction of high-molecular-weight DNAs was performed using NucleoBond HMW DNA (TAKARA). Library preparation was performed using the Ligation Sequencing Kit (Oxford Nanopore Technologies) and sequencing was performed using MinION (device: MIN-101B, flow cell: R9.4.1) at OIST SQC.

### DNA methylation calling of Nanopore sequence reads

Bases-calling of Col-0, *met1*, and *ddm1* was accomplished using Guppy (v4.4.2; https://nanoporetech.com/). DNA methylation at CG, CHG, or CHH was called with DeepSignal-Plant^53^. The unmethylated mitochondrial genomes were used as negative controls for the DNA methylation calling and to calculate the error rates. Estimated error rates were incorporated into a binomial test to assess evidence of methylation at each site, with q-value of <0.01^93^. Cytosine sites with non-significant calls were treated as unmethylated. Methylation levels were calculated using the ratio of mC/(mC + unmC), as previously indicated^94^, and were converted to bedGraph format for visualization. We also calculated the percentage of DNA methylation of TEs as previously described^27^. The difference in the percentage of TE DNA methylation in *met1* or *ddm1* compared with Col-0 was also calculated (Supplementary Data 3).

### GO term/TE enrichment and TE-RI splicing site motifs search

AGI codes of gene models in the DRS-AtRTD3 transcriptome were retrieved for DRS-AtRTD3 from the original AtRTD3 data using gffcompare^95^. AGI codes of genes involved in TE-Gt and altTE-Gi events were retrieved and submitted independently to ShinyGO (v0.75)^96^. For each TE superfamily, the expected number of TEs for each type of altTE-Gi event was calculated with the following equation: Expected number of TEs belonging to a superfamily = ([Number of TEs in the superfamily]*[Number of TEs involved in the altTE-Gi events])/Total number of TEs.

Fold-change = [Expected number of TEs belonging to a superfamily]/[Number of TEs belonging to a superfamily involved in the altTE-Gi events]. *P*-values for under- or over-enrichment were calculated based on the cumulative distribution function of hypergeometric distribution. Splicing sites of TE-IR for Mutator families VANDAL17 and ATMU6N1 were manually extracted (four nucleotides up/downstream of the acceptor and donor splicing site for every transcript), and figures were built with WebLogo^97^.

### ChIP-seq analysis

ChIP-seq data of RNA Pol II in wild-type Col-0 and the epigenetic mutants were obtained as previously described^23^ and using anti-RNA polymerase II CTD repeat YSPTSPS (phospho S2, ab5095; Abcam, Cambridge, UK) and anti-RNA polymerase II CTD repeat YSPTSPS (phospho S5, ab5408; Abcam) antibodies. ChIP-seq for FLAG-HA-IBM2 and EDM2-MYC-HA was performed using an anti-HA antibody (ab9110; abcam). ChIP-seq peak of H3K9me2 in Col-0 and *ibm1* were retrieved from published data^98^. Raw sequence reads were trimmed with fastp (v0.21.0; parameters: -l 36 -r)^88^ and aligned to the *A. thaliana* genome (TAIR10) with Bowtie2 (v2.4.2). ChIP peak calling was performed with MACS2 (v2.2.7.1; parameters: --broad --cutoff-analysis). Peaks overlapping with genes or TEs were extracted using the intersect function of bedtools (v2.29.2)^99^. For figures, normalized (log2 [ChIP/input]) bigwig files were generated with the bamCoverage function of deepTools (v3.4.3; parameters: --scaleFactorsMethod None -bs 10 -- normalizeUsing BPM -e 150 --operation log2)^100^ and visualized using Integrated Genome Viewer (IGV)^101^.

### Isoform switching and differential expression analysis

Illumina short reads from RNA-seq were mapped on transcriptome assemblies with Hisat2^102^ (v2.2.0; parameters: -p 100 --no-mixed --max-intronlen 10000). The tool Salmon^103^ (v1.3.0) was used to perform the read count. Isoform switching under various stress conditions or in mutants was predicted with the pipeline IsoformSwitchAnalyzeR (v1.6.0)^104^ employing DEXSeq^105^ with “isoform switch q-value” less than 0.05. The functions switchPlotIsoUsage and switchPlotIsoExp were used to investigate the *AT2G40960* locus (Fig. 3c). Differential expression of isoforms was detected with DESeq2^106^ by importing the Salmon read counts with tximport^107^.

### Identification of representative epigenetically regulated Epi-altTE-Gis

To identify Epi-altTE-Gis, we applied the following filtering steps to sort candidate altTE-Gis detected in DRS-AtRTD3 or in mutant transcriptomes by ParasiTE (Supplementary Data 5, 6). For Epi-altTE-Gis in *met1* or *ddm1*, we sorted Epi-altTE-Gi candidates that were associated with reduced DNA methylation in the mutants. For Epi-altTE-Gis in *suvh456*, we searched for overlaps of altTE-Gi loci (TE and transcript) with ChIP-seq signals of Col-0 H3K9me2. For Epi-altTE-Gis in *ibm1*, we searched for overlaps of altTE-Gi loci (TE and transcript) with ChIP-seq signals of Col-0 H3K9me2 and *ibm1* H3K9me2^98^. For Epi-altTE-Gis in *ibm2* and *edm2*, we searched for overlaps of altTE-Gi loci (TE and transcript) with ChIP-seq signals of IBM2, EDM2, as well as Col-0 H3K9me2^98^. Finally, candidate loci were manually filtered by visualization of mapped DRS reads, published CAGE-seq datasets of Col-0, *met1, ddm1*, and *suvh456^23^*, and ChIP-seq signals and DNA methylation levels using IGV.

### Investigation of mutants and stress conditions on Epi-altTE-Gi expression and isoform switching events

A consensus transcriptome was built by combining DRS-AtRTD3 and DRS-based mutant transcriptomes using GffCompare^95^ (v0.12.6; -r with the AtRTD3 annotation as a reference). Next, Epi-altTE-Gi candidates that were retrieved, as described above, were manually selected. A list and information about the RNA-seq data (short paired-end reads) used for the stress condition analyses are provided in Supplementary Table 4. For the heatmap presentation in Figure 7, a maximum of five Epi-altTE-Gis are shown for each gene, and only altTE-Gis with at least one significant isoform switching event in an epigenetic mutant are presented. Because of the complexity of *RPP4* altTE-Gis, only the first five altTE-Gis in Figure 6a are presented in the heatmap. Significant isoform switching was analyzed using IsoformSwitchAnalyzeR (v1.6.0)^104^. The function switchPlotIsoUsage was used to investigate the *GER5* and *F-box* loci (Supplementary Fig. 20). The transcripts in the heatmaps presenting isoform switching were also analyzed for differential expression by DESeq2^106^.

### Poly(A) site prediction analyses

Uncorrected ONT-DRS reads of Col-0 (unconverted from cDNA and uncorrected with LorDEC) were mapped to the *Arabidopsis* genome (TAIR10) with minimap2 (-ax splice -uf -k14). Nanopolish^108^ was used to predict ONT-DRS reads with poly(A) tails. Bam files were converted into bed files, and for reads predicted to bear poly(A) tail, the poly(A) sites were assumed to be located at the 3’ end of the read and at ±10 bp from the soft-clipped nucleotide (soft-clipping done by minimap2). The locations of TEs involved in TE-ATTS events predicted by ParasiTE were compared to the APA predictions obtained with ONT-DRS and APA sites retrieved from the PlantAPAdb^55^ (High confidence annotation; http://www.bmibig.cn/plantAPAdb/Bulkdownload.php, using bedtool’s intersect function^99^ (v2.29.2).

### Reverse transcription PCR (RT-PCR) and qPCR

For each experiment, at least three plants were pooled for a replicate, and four independent biological replicates were prepared for each experiment. Total RNA was extracted from 14-day-old seedlings (12-days old for the cordycepin experiments) using the Maxwell 16 LEV Plant RNA Kit and Maxwell 16 Instrument (Promega, Madison, WI). For RT-PCR and qRT-PCR, cDNA was synthesized using Prime Script II (TAKARA) following the supplier’s protocol. RT-PCR was performed using GoTaq DNA Polymerase (Promega) using a T100 Thermal cycler (Bio-Rad, USA). RT-qPCR was performed using TB Green Premix ExTaq II (Tli RNAseH Plus) (TAKARA) with Thermal Cycler Dice ® Real Time System III (one cycle of 95°C for 30 s followed by 40 cycles of 95 °C for 5 s and 60 °C for 30 s). Primer specificity was validated by a dissociation curve. *ACTIN2 (AT3G18780*) and *GAPDH* (AT1G13440) were used as housekeeping genes, as previously recommended^109^. The 2^-ΔΔCT^ method was used to determine mRNA expression levels^110^. All primers used in this study are listed in Supplementary Table 5 and 6.

### *RPP4* ORF prediction

*RPP4* transcripts (annotated from a to g in Supplementary Fig. 15, 18) were predicted from the minimap2 alignment of corrected ONT-DRS reads of Col-0 to the *A. thaliana* genome (TAIR10). ORFs of transcripts were predicted using ORF Finder (https://www.ncbi.nlm.nih.gov/orffinder/), and CDS prediction was performed (annotated A to G in Supplementary Fig. 15). Multialignments were generated with ClustalΩ (https://www.ebi.ac.uk/Tools/msa/clustalo/) and visualized with Geneious (https://www.geneious.com). Domain and homologous superfamily predictions of corresponding amino acid sequences were obtained with InterProScan (https://www.ebi.ac.uk/interpro/search/sequence/).

### ABA treatment

Fourteen-day-old Col-0 and mutants were grown in a pot and 50 μM of ABA + 0.01% SILWET L-77 was sprayed onto the plants. Only water and 0.01% SILWET L-77 was sprayed onto the leaves of the mock plants. Biological replicates were harvested after 2 hours of treatment and frozen in liquid nitrogen.

### Heat shock treatment

The plants were subjected to heat shock stress as described by Ito et al., 2011. Twelve-day-old Col-0 were grown in pots at 21°C before being cooled at 6°C for 24 h. Next, the control plants were moved back to the 21°C cabinet while the others were subjected to heat shock treatment in a growth chamber (Biotron LPH-410SP; NK system, Japan) at 37°C for 24 hours. Biological replicates were harvested after treatment and frozen in liquid nitrogen.

### Cordycepin treatment of the *Arabidopsis* strains

Twelve-day-old Col-0, Ler-0, Sha, and epigenetic mutants were grown on 1/2 MS plates, and experiments were independently performed for each genotype. For each genotype, around 80 plants were transferred into a glass plate containing 80 mL buffer (1 mM PIPES, 1 mM trisodium citrate, 1 mM KCl, and 15 mM sucrose [final pH of 6.25]), as previously described^37^. The plants were incubated for 30 min in the buffer and covered with a transparent tissue layer that was engulfed by the solution. The plants were kept under light and were slowly shaken. Portions of the plants were collected after 30 min incubation; this corresponded to the time 0. Next, 20 mL of 3 mM of 3-deoxyadenosine (cordycepin; Thermo Fisher Scientific) was added to the buffer to obtain a final concentration of 0.6 mM (150 mg/L), and vacuum infiltration was performed for 30 s at 0.04 MPa. The plants were collected at time points t = 30, 60, 90, and 120 min after the start of the treatment and frozen in liquid nitrogen. RT-qPCR was performed as described above. The expression levels of genes at each time point were first normalized using the expression levels of *ACTIN2* and *GAPDH* and then normalized to the expression levels at time 0. Eukaryotic initiation factor-4A (EIF4A1, *AT3G13920*) and EXPANSIN-LIKE A1 (EXLA1, *AT3G45970*) were used as control genes for high and low mRNA stability, respectively, as previously described^37,111^. Primers and the efficiency of primers used for this experiment are reported in Supplementary Table 5.

### *Hpa* resistance assays

Inoculation with *Hpa* Emoy2 was accomplished as described previously^112^. Briefly, the *Arabidopsis* plants were spray-inoculated to saturation with a spore suspension of 1 × 10^5^ conidiospores/mL of Emoy2. Plants were covered with a transparent lid to maintain high humidity (90–100%) conditions in a growth cabinet at 16°C under a 10-h photoperiod until the day of sampling. To evaluate hyphae growth and dead cells, leaves inoculated with Emoy2 were stained with trypan blue. Infected leaves were transferred to trypan blue solution (10 mL of lactic acid, 10 mL of glycerol, 10 g of phenol, 10 mL of H_2_O, and 10 mg of trypan blue) diluted in ethanol (1:1 v/v) and boiled for 1 min. Leaves were destained overnight in chloral hydrate solution (60% w/v) and then stored in 60% (v/v) glycerol.

Trypan blue-stained leaves were analyzed with a stereomicroscope (KL200 LED; Leica, Schott, Germany), and assigned to different classes based on Emoy2 interaction: class I (white), hypersensitive response surrounding conidia penetration sites; class II (light green), presence of trailing necrosis in ≤50% leaf area; class III (dark green), presence of trailing necrosis in ≤75% leaf area; class IV (black), compromised ETI immunity, presence of pathogen hyphae not targeted by hypersensitive response (HR) and conidiophores. Statistically significant differences in frequency distribution of the *Hpa* colonization classes between lines and Col-0 were determined by Pearson’s chi-squared test. Between 70–130 leaves were analyzed per line across three separate experimental replicates.

## Data availability

Sequencing data obtained in this study have been deposited. Illumina RNA-seq data: XXXXXXXXXX; ChIP-seq data of epigenetics mutants: XXXXXXXXXX; ChIP-seq data of RNA Pol II in Col-0, *ibm2*, and *edm2:* XXXXXXXXXX; ONT-DRS data of epigenetics mutants: XXXXXXXXXXX; ONT DNA of Col-0, *met1*, and *ddm1*: XXXXXXXXXX. Processed ONT-DRS transcriptomes, ONT methylation data, and ChIP-seq data are also accessible via the web interface: https://plantepigenetics.oist.jp/.

## Code availability

ParasiTE is available in Github https://github.com/JBerthelier/ParasiTE.

## Acknowledgements

This work was supported by MEXT Grant-in-Aid for Transformative Research Areas (A) JP20H05913 to H.S., JP20H05909 and JP22H00364 to K.S., JP20H02995 to S.A., and by OIST. We thank the Arabidopsis Biological Resource Center and the Salk Institute Genomic Analysis Laboratory for providing *Arabidopsis* T-DNA insertion mutants, OIST SQC for Nanopore DRS-seq and DNA-seq, Illumina RNA-seq, ChIP-seq, and BS-seq sequencing services, Dr. Tetsuji Kakutani for providing mutant seeds, OIST IT Section for technical supports in building the web interface to access data, as well as Dr. Tu Le and OIST English editing service for proofreading of the manuscript.

## Author contributions

DRS experiments were designed by J.B. and H.S. and performed by T.S., J.B., and H.S. mRNA expression experiments were designed by J.B., H.S., and L.F. and performed by J.B. with the help of L.F. Pathogen experiments were performed by S.A., K.S., and L.F. ParasiTE was developed by J.B. Sequence data analysis was performed by J.B. and with the help of M.S for ONT methylation calling. The manuscript was prepared by J.B. and H.S.

## References

1. Bennetzen, J. L. & Wang, H. The contributions of transposable elements to the structure, function, and evolution of plant genomes. Annual review of plant biology 65, 505–530 (2014).

2. Bourque, G. et al. Ten things you should know about transposable elements. Genome biology 19, 1–12 (2018).

3. Furci, L., Berthelier, J., Juez, O., Miryeganeh, M. & Saze, H. Plant Epigenomics. in Handbook of Epigenetics 263–286 (Elsevier, 2023).

4. Casacuberta, E. & González, J. The impact of transposable elements in environmental adaptation. Molecular Ecology 22, 1503–1517 (2013).

5. Galindo-González, L., Mhiri, C., Deyholos, M. K. & Grandbastien, M.-A. LTR-retrotransposons in plants: Engines of evolution. Gene 626, 14–25 (2017).

6. Hirsch, C. D. & Springer, N. M. Transposable element influences on gene expression in plants. Biochimica et Biophysica Acta (BBA) - Gene Regulatory Mechanisms 1860, 157–165 (2017).

7. Cosby, R. L. et al. Recurrent evolution of vertebrate transcription factors by transposase capture. Science 371, eabc6405 (2021).

8. Nekrutenko, A. & Li, W.-H. Transposable elements are found in a large number of human protein-coding genes. TRENDS in Genetics 17, 619–621 (2001).

9. Lev-Maor, G., Sorek, R., Shomron, N. & Ast, G. The birth of an alternatively spliced exon: 3’splice-site selection in Alu exons. Science 300, 1288–1291 (2003).

10. Sela, N. et al. Comparative analysis of transposed element insertion within human and mouse genomes reveals Alu’s unique role in shaping the human transcriptome. Genome biology 8, 1–19 (2007).

11. Lockton, S. & Gaut, B. S. The contribution of transposable elements to expressed coding sequence in Arabidopsis thaliana. Journal of molecular evolution 68, 80–89 (2009).

12. Lanciano, S. & Cristofari, G. Measuring and interpreting transposable element expression. Nature Reviews Genetics 21, 721–736 (2020).

13. Reddy, A. S. N., Marquez, Y., Kalyna, M. & Barta, A. Complexity of the Alternative Splicing Landscape in Plants. The Plant Cell 25, 3657–3683 (2013).

14. Lee, J. Y., Ji, Z. & Tian, B. Phylogenetic analysis of mRNA polyadenylation sites reveals a role of transposable elements in evolution of the 3’-end of genes. Nucleic acids research 36, 5581–5590 (2008).

15. Filichkin, S. A. et al. Genome-wide mapping of alternative splicing in Arabidopsis thaliana. Genome research 20, 45–58 (2010).

16. Martín, G., Márquez, Y., Mantica, F., Duque, P. & Irimia, M. Alternative splicing landscapes in Arabidopsis thaliana across tissues and stress conditions highlight major functional differences with animals. Genome biology 22, 1–26 (2021).

17. Law, J. A. & Jacobsen, S. E. Establishing, maintaining and modifying DNA methylation patterns in plants and animals. Nature Reviews Genetics 11, 204–220 (2010).

18. Lippman, Z. et al. Role of transposable elements in heterochromatin and epigenetic control. Nature 430, 471–476 (2004).

19. Saze, H., Shiraishi, A., Miura, A. & Kakutani, T. Control of genic DNA methylation by a jmjC domain-containing protein in Arabidopsis thaliana. science 319, 462–465 (2008).

20. Brocks, D. et al. DNMT and HDAC inhibitors induce cryptic transcription start sites encoded in long terminal repeats. Nature genetics 49, 1052–1060 (2017).

21. Neri, F. et al. Intragenic DNA methylation prevents spurious transcription initiation. Nature 543, 72–77 (2017).

22. Clayton, E. A. et al. An atlas of transposable element-derived alternative splicing in cancer. Phil. Trans. R. Soc. B 375, 20190342 (2020).

23. Le, N. T. et al. Epigenetic regulation of spurious transcription initiation in Arabidopsis. Nat Commun 11, 3224 (2020).

24. Zabala, G. & Vodkin, L. O. Methylation affects transposition and splicing of a large CACTA transposon from a MYB transcription factor regulating anthocyanin synthase genes in soybean seed coats. PLoS One 9, e111959 (2014).

25. Ong-Abdullah, M. et al. Loss of Karma transposon methylation underlies the mantled somaclonal variant of oil palm. Nature 525, 533–537 (2015).

26. Le, T. N., Miyazaki, Y., Takuno, S. & Saze, H. Epigenetic regulation of intragenic transposable elements impacts gene transcription in Arabidopsis thaliana. Nucleic Acids Research 43, 3911–3921 (2015).

27. Espinas, N. A. et al. Transcriptional regulation of genes bearing intronic heterochromatin in the rice genome. PLoS genetics 16, e1008637 (2020).

28. Eulgem, T. et al. EDM2 is required for RPP7-dependent disease resistance in Arabidopsis and affects RPP7 transcript levels: EDM2-mediated disease resistance. The Plant Journal 49, 829–839 (2007).

29. Saze, H. et al. Mechanism for full-length RNA processing of Arabidopsis genes containing intragenic heterochromatin. Nat Commun 4, 2301 (2013).

30. Coustham, V. et al. SHOOT GROWTH1 maintains Arabidopsis epigenomes by regulating IBM1. PLoS One 9, e84687 (2014).

31. Duan, C.-G. et al. A protein complex regulates RNA processing of intronic heterochromatin-containing genes in *Arabidopsis*. Proc Natl Acad Sci USA 114, E7377–E7384 (2017).

32. Lai, Y. et al. The Arabidopsis PHD-finger protein EDM2 has multiple roles in balancing NLR immune receptor gene expression. PLoS Genet 16, e1008993 (2020).

33. Zhang, Y. et al. Genome-wide distribution and functions of the AAE complex in epigenetic regulation in *Arabidopsis*. J. Integr. Plant Biol. 63, 707–722 (2021).

34. Guo, C., Spinelli, M., Liu, M., Li, Q. Q. & Liang, C. A genome-wide study of “non-3UTR” polyadenylation sites in Arabidopsis thaliana. Scientific reports 6, 1–10 (2016).

35. Mayr, C. Regulation by 3’-untranslated regions. Annu Rev Genet 51, 171–194 (2017).

36. Shen, J. et al. Translational repression by a miniature inverted-repeat transposable element in the 3′ untranslated region. Nature Communications 8, 1–10 (2017).

37. Niu, X.-M. et al. Transposable elements drive rapid phenotypic variation in <em>Capsella rubella</em>. Proc Natl Acad Sci USA 116, 6908 (2019).

38. Zhang, S. et al. New insights into Arabidopsis transcriptome complexity revealed by direct sequencing of native RNAs. Nucleic acids research 48, 7700–7711 (2020).

39. Parker, M. T. et al. Nanopore direct RNA sequencing maps the complexity of Arabidopsis mRNA processing and m6A modification. eLife 9, e49658 (2020).

40. Jia, J. et al. Post-transcriptional splicing of nascent RNA contributes to widespread intron retention in plants. Nature Plants 6, 780–788 (2020).

41. Mo, W. et al. Landscape of transcription termination in Arabidopsis revealed by singlemolecule nascent RNA sequencing. Genome biology 22, 1–21 (2021).

42. Parker, M. T. et al. Widespread premature transcription termination of Arabidopsis thaliana NLR genes by the spen protein FPA. eLife 10, e65537 (2021).

43. Zhang, R. et al. A high-resolution single-molecule sequencing-based Arabidopsis transcriptome using novel methods of Iso-seq analysis. Genome Biology 23, 149 (2022).

44. Panda, K. & Slotkin, R. K. Long-read cDNA sequencing enables a “gene-like” transcript annotation of transposable elements. Plant Cell 32, 2687–2698 (2020).

45. Garalde, D. R. et al. Highly parallel direct RNA sequencing on an array of nanopores. Nature methods 15, 201–206 (2018).

46. van der Biezen, E. A., Freddie, C. T., Kahn, K., ParkerP, J. E. & Jones, J. D. G. Arabidopsis RPP4 is a member of the RPP5 multigene family of TIR-NB-LRR genes and confers downy mildew resistance through multiple signalling components. Plant J 29, 439–451 (2002).

47. Cheng, C. et al. Araport11: a complete reannotation of the Arabidopsis thaliana reference genome. The Plant Journal 89, 789–804 (2017).

48. Kovaka, S. et al. Transcriptome assembly from long-read RNA-seq alignments with StringTie2. Genome Biol 20, 278 (2019).

49. Pinson, M.-E., Pogorelcnik, R., Court, F., Arnaud, P. & Vaurs-Barrière, C. CLIFinder: identification of LINE-1 chimeric transcripts in RNA-seq data. Bioinformatics 34, 688–690 (2018).

50. Babaian, A. et al. LIONS: analysis suite for detecting and quantifying transposable element initiated transcription from RNA-seq. Bioinformatics 35, 3839–3841 (2019).

51. Shiau, C.-K., Huang, J.-H. & Tsai, H.-K. CATANA: a tool for generating comprehensive annotations of alternative transcript events. Bioinformatics 35, 1414–1415 (2019).

52. Wicker, T. et al. A unified classification system for eukaryotic transposable elements. Nature Reviews Genetics 8, 973–982 (2007).

53. Ni, P. et al. Genome-wide detection of cytosine methylations in plant from Nanopore data using deep learning. Nature communications 12, 1–11 (2021).

54. Yan, X. et al. DNA methylation signature of intergenic region involves in nucleosome remodeler DDM1-mediated repression of aberrant gene transcriptional read-through. Journal of Genetics and Genomics 43, 513–523 (2016).

55. Zhu, S. et al. PlantAPAdb: a comprehensive database for alternative polyadenylation sites in plants. Plant physiology 182, 228–242 (2020).

56. Tsuchiya, T. & Eulgem, T. Mutations in EDM2 selectively affect silencing states of transposons and induce plant developmental plasticity. Scientific reports 3, 1–9 (2013).

57. Baron, K. N., Schroeder, D. F. & Stasolla, C. GEm-Related 5 (GER5), an ABA and stress-responsive GRAM domain protein regulating seed development and inflorescence architecture. Plant Science 223, 153–166 (2014).

58. Chen, C.-Y. et al. AU binding proteins recruit the exosome to degrade ARE-containing mRNAs. Cell 107, 451–464 (2001).

59. Asai, S. et al. A downy mildew effector evades recognition by polymorphism of expression and subcellular localization. Nature Communications 9, 1–11 (2018).

60. Ma, S. et al. Direct pathogen-induced assembly of an NLR immune receptor complex to form a holoenzyme. Science 370, eabe3069 (2020).

61. Deremetz, A. et al. Antagonistic Actions of FPA and IBM2 Regulate Transcript Processing from Genes Containing Heterochromatin. Plant Physiol. 180, 392–403 (2019).

62. Asai, S. et al. Expression profiling during Arabidopsis/downy mildew interaction reveals a highly-expressed effector that attenuates responses to salicylic acid. PLoS Pathogens 10, e1004443 (2014).

63. Krasileva, K. V. et al. Global analysis of Arabidopsis/downy mildew interactions reveals prevalence of incomplete resistance and rapid evolution of pathogen recognition. PLoS One 6, e28765 (2011).

64. Hu, L. & Yang, L. Time to fight: Molecular mechanisms of age-related resistance. Phytopathology 109, 1500–1508 (2019).

65. Wang, Y.-H. & Warren Jr, J. T. Mutations in retrotransposon AtCOPIA4 compromises resistance to Hyaloperonospora parasitica in Arabidopsis thaliana. Genetics and molecular biology 33, 135–140 (2010).

66. Sorek, R., Ast, G. & Graur, D. Alu-Containing Exons are Alternatively Spliced. Genome Res. 12, 1060–1067 (2002).

67. Oki, N. et al. A genome-wide view of miniature inverted-repeat transposable elements (MITEs) in rice, Oryza sativa ssp. japonica. Genes & genetic systems 83, 321–329 (2008).

68. Zhang, X., Zhao, M., McCarty, D. R. & Lisch, D. Transposable elements employ distinct integration strategies with respect to transcriptional landscapes in eukaryotic genomes. Nucleic acids research 48, 6685–6698 (2020).

69. Saze, H. & Kakutani, T. Heritable epigenetic mutation of a transposon-flanked Arabidopsis gene due to lack of the chromatin-remodeling factor DDM1. EMBO J 26, 3641–3652 (2007).

70. Ito, H. et al. An siRNA pathway prevents transgenerational retrotransposition in plants subjected to stress. Nature 472, 115–119 (2011).

71. Leandro, Q. et al. The Arabidopsis thaliana mobilome and its impact at the species level. eLife 5, (2016).

72. Stuart, T. et al. Population scale mapping of transposable element diversity reveals links to gene regulation and epigenomic variation. elife 5, e20777 (2016).

73. Baduel, P. et al. Genetic and environmental modulation of transposition shapes the evolutionary potential of Arabidopsis thaliana. Genome biology 22, 1–26 (2021).

74. Mohn, F., Sienski, G., Handler, D. & Brennecke, J. The rhino-deadlock-cutoff complex licenses noncanonical transcription of dual-strand piRNA clusters in Drosophila. Cell 157, 1364–1379 (2014).

75. Chen, Y.-C. A. et al. Cutoff suppresses RNA polymerase II termination to ensure expression of piRNA precursors. Molecular cell 63, 97–109 (2016).

76. Cavrak, V. V. et al. How a retrotransposon exploits the plant’s heat stress response for its activation. PLoS genetics 10, e1004115 (2014).

77. Roquis, D. et al. Genomic impact of stress-induced transposable element mobility in Arabidopsis. Nucleic Acids Research 49, 10431–10447 (2021).

78. Annacondia, M. L. et al. Aphid feeding induces the relaxation of epigenetic control and the associated regulation of the defense response in Arabidopsis. New Phytologist 230, 1185–1200 (2021).

79. Meyers, B. C., Kozik, A., Griego, A., Kuang, H. & Michelmore, R. W. Genome-wide analysis of NBS-LRR–encoding genes in Arabidopsis. The Plant Cell 15, 809–834 (2003).

80. Kawakatsu, T. et al. Epigenomic diversity in a global collection of Arabidopsis thaliana accessions. Cell 166, 492–505 (2016).

81. Yi, H. & Richards, E. J. A cluster of disease resistance genes in Arabidopsis is coordinately regulated by transcriptional activation and RNA silencing. The Plant Cell 19, 2929–2939 (2007).

82. Gloggnitzer, J. et al. Nonsense-mediated mRNA decay modulates immune receptor levels to regulate plant antibacterial defense. Cell host & microbe 16, 376–390 (2014).

83. Raxwal, V. K. et al. Nonsense-mediated RNA decay factor UPF1 is critical for posttranscriptional and translational gene regulation in Arabidopsis. Plant Cell 32, 2725–2741 (2020).

84. Yu, A. et al. Dynamics and biological relevance of DNA demethylation in Arabidopsis antibacterial defense. Proceedings of the National Academy of Sciences 110, 2389–2394 (2013).

85. Vongs, A., Kakutani, T., Martienssen, R. A. & Richards, E. J. Arabidopsis thaliana DNA methylation mutants. Science 260, 1926–1928 (1993).

86. Saze, H., Scheid, O. M. & Paszkowski, J. Maintenance of CpG methylation is essential for epigenetic inheritance during plant gametogenesis. Nature genetics 34, 65–69 (2003).

87. Osabe, K., Harukawa, Y., Miura, S. & Saze, H. Epigenetic regulation of intronic transgenes in Arabidopsis. Scientific reports 7, 1–13 (2017).

88. Chen, S., Zhou, Y., Chen, Y. & Gu, J. fastp: an ultra-fast all-in-one FASTQ preprocessor. Bioinformatics 34, i884–i890 (2018).

89. Shen, W., Le, S., Li, Y. & Hu, F. SeqKit: a cross-platform and ultrafast toolkit for FASTA/Q file manipulation. PloS one 11, e0163962 (2016).

90. Salmela, L. & Rivals, E. LoRDEC: accurate and efficient long read error correction. Bioinformatics 30, 3506–3514 (2014).

91. Li, H. Minimap2: pairwise alignment for nucleotide sequences. Bioinformatics 34, 3094–3100 (2018).

92. Kovaka, S. Transcriptome assembly from long-read RNA-seq alignments with StringTie2. 13 (2019).

93. Lister, R. et al. Highly integrated single-base resolution maps of the epigenome in Arabidopsis. Cell 133, 523–536 (2008).

94. Stroud, H., Greenberg, M. V., Feng, S., Bernatavichute, Y. V. & Jacobsen, S. E. Comprehensive analysis of silencing mutants reveals complex regulation of the Arabidopsis methylome. Cell 152, 352–364 (2013).

95. Pertea, G. & Pertea, M. GFF utilities: GffRead and GffCompare. F1000Research 9, (2020).

96. Ge, S. X., Jung, D. & Yao, R. ShinyGO: a graphical gene-set enrichment tool for animals and plants. Bioinformatics 36, 2628–2629 (2020).

97. Crooks, G. E., Hon, G., Chandonia, J.-M. & Brenner, S. E. WebLogo: a sequence logo generator. Genome research 14, 1188–1190 (2004).

98. Inagaki, S. et al. Gene-body chromatin modification dynamics mediate epigenome differentiation in Arabidopsis. The EMBO journal 36, 970–980 (2017).

99. Quinlan, A. R. & Hall, I. M. BEDTools: a flexible suite of utilities for comparing genomic features. Bioinformatics 26, 841–842 (2010).

100. Ramírez, F. et al. deepTools2: a next generation web server for deep-sequencing data analysis. Nucleic acids research 44, W160–W165 (2016).

101. Thorvaldsdóttir, H., Robinson, J. T. & Mesirov, J. P. Integrative Genomics Viewer (IGV): high-performance genomics data visualization and exploration. Briefings in bioinformatics 14, 178–192 (2013).

102. Kim, D., Paggi, J. M., Park, C., Bennett, C. & Salzberg, S. L. Graph-based genome alignment and genotyping with HISAT2 and HISAT-genotype. Nature biotechnology 37, 907–915 (2019).

103. Patro, R., Duggal, G., Love, M. I., Irizarry, R. A. & Kingsford, C. Salmon provides fast and bias-aware quantification of transcript expression. Nature methods 14, 417–419 (2017).

104. Anders, S. & Huber, W. Differential expression analysis for sequence count data. Nature Precedings 1–1 (2010).

105. Li, Y., Rao, X., Mattox, W. W., Amos, C. I. & Liu, B. RNA-seq analysis of differential splice junction usage and intron retentions by DEXSeq. PloS one 10, e0136653 (2015).

106. Love, M. I., Huber, W. & Anders, S. Moderated estimation of fold change and dispersion for RNA-seq data with DESeq2. Genome biology 15, 1–21 (2014).

107. Soneson, C., Love, M. I. & Robinson, M. D. Differential analyses for RNA-seq: transcript-level estimates improve gene-level inferences. F1000Research 4, (2015).

108. Loman, N. J., Quick, J. & Simpson, J. T. A complete bacterial genome assembled de novo using only nanopore sequencing data. Nature methods 12, 733–735 (2015).

109. Czechowski, T., Stitt, M., Altmann, T., Udvardi, M. K. & Scheible, W.-R. Genome-Wide Identification and Testing of Superior Reference Genes for Transcript Normalization in Arabidopsis. Plant Physiology 139, 5–17 (2005).

110. Livak, K. J. & Schmittgen, T. D. Analysis of relative gene expression data using real-time quantitative PCR and the 2-ΔΔCT method. methods 25, 402–408 (2001).

111. Golisz, A., Sikorski, P. J., Kruszka, K. & Kufel, J. Arabidopsis thaliana LSM proteins function in mRNA splicing and degradation. Nucleic Acids Research 41, 6232–6249 (2013).

112. Asai, S., Shirasu, K. & Jones, J. D. G. Hyaloperonospora arabidopsidis (Downy Mildew) Infection Assay in Arabidopsis. Bio-protocol 5, e1627 (2015).

