## Supplementary Note 1 for "Long-read direct RNA sequencing reveals epigenetic regulation of chimeric gene-transposon transcripts in *Arabidopsis thaliana*"

Long-read direct RNA sequencing reveals epigenetic regulation of chimeric gene-transposon transcripts in *Arabidopsis thaliana*

Berthelier *et al.*

**Detection of TE-gene transcripts by the ParasiTE pipeline.**

Input data for the ParasiTE pipeline:

The following four input data were used for ParasiTE analysis in this study:

1) Transcriptome annotation of *A. thaliana* Araport11 (https://phytozome-next.jgi.doe.gov/info/Athaliana_Araport11, "Athaliana_447_Araport11.gene_exons.gff3")[1] or AtRTD3[2] or DRS-Araport11 or DRS-AtRTD3 or mutant-DRS transcriptomes obtained with Stringtie2[3].

2) Gene model annotation of *A. thaliana* Araport11 (https://phytozome-next.jgi.doe.gov/info/Athaliana_Araport11, "Athaliana_447_Araport11.gene.gff3")[1] or AtRTD3 (https://ics.hutton.ac.uk/atRTD/RTD3/)[2]. Gene and transcript feature of AtRTD3 gtf were retrieved using gffread (v0.12.2; parameters: --keep-genes)[4].

3) TE annotation of *A. thaliana* of TAIR10 (retrieved from the URGI laboratory browser; https://urgi.versailles.inra.fr/gb2/gbrowse/tairv10_pub_TEs/)[5]. Only TEs with a length ≥200 bp were analyzed in this study.

4) A gene-like TE annotation retrieved from TAIR (ftp://ftp.arabidopsis.org/home/tair/Genes/TAIR10_genome_release/TAIR10_gff3/TAIR10_GFF3_genes_transposons.gff) and published gene-like TE annotation data by Panda et al.[6] were merged with bedtools' merge function[7].

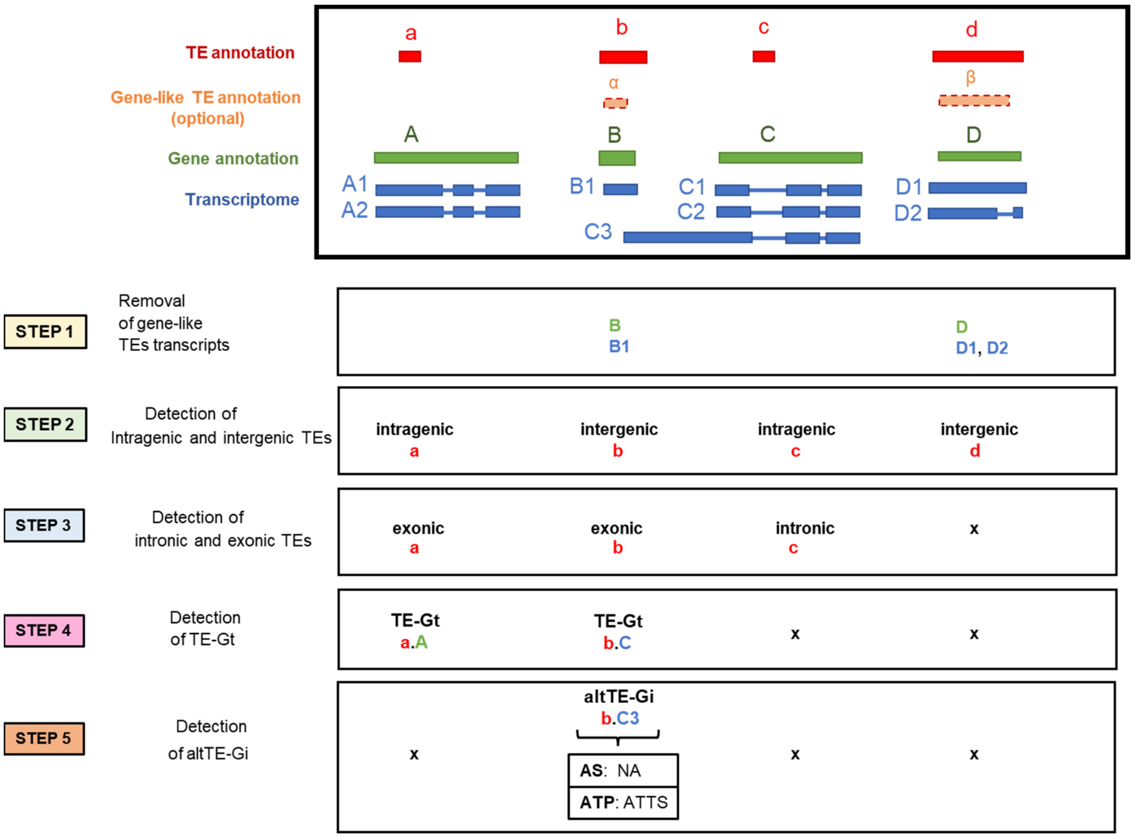


**Supplementary Note Figure 1.** Main steps of the ParasiTE pipeline.

<u>Step1: Removal of gene-like TEs</u>

ParasiTE first detects gene models of active TEs called "gene-like TEs"[6]. Next, it removes the annotation of gene-like TEs from the gene model annotation and the associated transcripts in the transcriptome annotation (Supplementary Note Figure 1). Indeed, gene-like TEs are the major cause of false positive events in the detection of TE-gene transcripts (TE-Gts) or alternative TE-gene transcript isoforms (altTE-Gis) in downstream analysis.

For this task, ParasiTE uses bedtools' intersect function[7] to compare the gene model annotation with the provided TE annotation. ParasiTE detects gene annotations overlapping TEs by at least 80% in length, which are considered to be gene-like TEs. ParasiTE generates a list of these gene models and removes them from the original gene model annotation. Next, ParasiTE uses the created list of gene models, defined as gene-like TEs, to remove transcripts (and associated exons) in the transcriptome dataset that overlap their annotation by least 80% in length.

Optionally, a gene-like TE annotation can be provided. In this case, ParasiTE applies the second round of filtering and identifies leftover transcripts that overlap the provided gene-like TE annotation by at least 80% in length and removes the transcript (and associated exon) annotations from the transcriptome dataset. This step may help to detect gene-like TEs that have been imprecisely identified in the TE annotation (for example, because of fragmented TE annotation). This step improves the mis-annotation of gene-like TEs as TE-Gts (Supplementary Note Figure 14).

Step 2: Detection of transcripts containing intragenic and intergenic TEs

In the next step, ParasiTE classifies TEs according to their location based on the gene annotation (Supplementary Note Figure 1). This step employs the bedtools intersect function[7]. TEs that overlap a gene by at least 80% in length are regarded as “intragenic TE”, and the remaining TEs as “intergenic TEs”. Then, ParasiTE looks for intergenic TEs close to genes potentially involved in TE-Gt formation. Intergenic TEs within 2 kb of an annotated gene are extracted by bedtools’ closest function[7] for further processing. Three rounds of extraction are applied at each side of the TE.

Step 3 and 4: Annotation of transcripts containing exonic and intronic TEs

Next, ParasiTE compares the intragenic and selected intergenic TEs to the transcriptome dataset with the bedtools intersect function[7] to annotate TEs that are fully exonic, partially exonic, or intronic. Two methods are applied to identify “exonic TEs”, with method 1 identifying TEs overlapping exons by at most 80% in length, while method 2 detects TEs that overlap more than 80% the length of an exon. Next, method 3 detects leftover TEs that overlap exons by at least 1% in length and are classified as "partially exonic TEs” (Supplementary Note Figure 2). Intragenic TEs not classified by either of these methods and which do not overlap by 1% or more an exon are considered as “intronic TEs”.

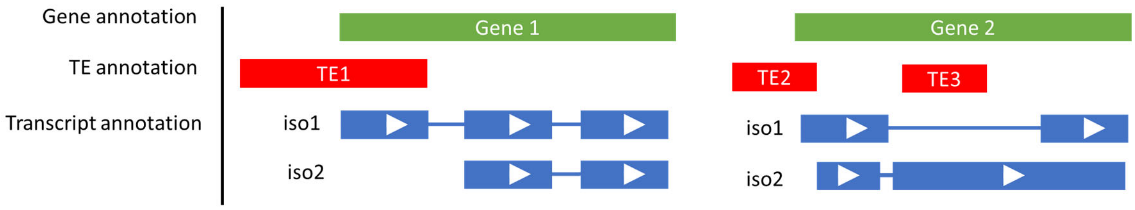


**Supplementary Note Figure 2:** Schematic illustration of different types of exonic TEs and methods used to find them. TE3 is detected by method 1, TE1 by method 2, and TE2 by method 3.

Step 4 and 5: Identification of TE-Gts and altTE-Gis

At this step, intragenic and intergenic TEs defined as "exonic TEs" or "partially exonic TEs" are used as TE-Gt candidates. Intergenic TEs may be identified as TE-Gt candidates if they were not properly annotated as part of the gene model annotation.
ParasiTE generates a list of candidate TEs and the associated transcripts and exons with which they form TE-Gt events. More precisely, ParasiTE lists every pair of TE and overlapping exon as TE-Gt candidates.

For example: "AT1G03410.1_845951_847683" (Exon id) & AT1TE02770 (TE id).

Next, ParasiTE takes advantage of the tool CATANA[8] to identify altTE-Gis among TE-Gt candidates in the transcriptome dataset. Using the transcriptome annotation (.gtf or .gff) as an input, CATANA detects and classifies the occurrence of alternative splicing (AS) and alternative transcription product (ATP) events for each exon, transcript, and gene. CATANA finds more AS and ATP events than the tools MISO[9] or ASprofile[10] and has the advantage of detecting both AS and ATP events[8].

ParasiTE uses this information to retrieve exon-TE pairs involved in altTE-Gi events. ParasiTE retrieves AS and ATP predictions from CATANA for exon-TE pair events (Supplementary Note Figure 1, step 5). This information is first given as raw output data and combined by ParasiTE to summarize the events appearing for every TE-gene pair. Although CATANA[8] can identify single and multiple skipping exons, this information is combined as "exon skipping" (ES) events in ParasiTE. Moreover, this first version of ParasiTE does not consider mutually exclusive exon (MXE) events found by CATANA[8].

ParasiTE eventually identifies alternative TE-related splicing sites (TE-AS) composed of alternative TE-related 5'/3' splice sites (TE-A5SS/TE-A3SS), TE-related exon skipping (TE-ES), and TE-related intron retention (TE-IR). Moreover, it can identify TE-related ATPs composed of TE-related alternative transcriptional start/termination sites (TE-ATSS/TE-ATTS), and among those, it predicts TE-related alternative first/last exons (TE-AFE/TE-ALE; Supplementary Note Figure 3).

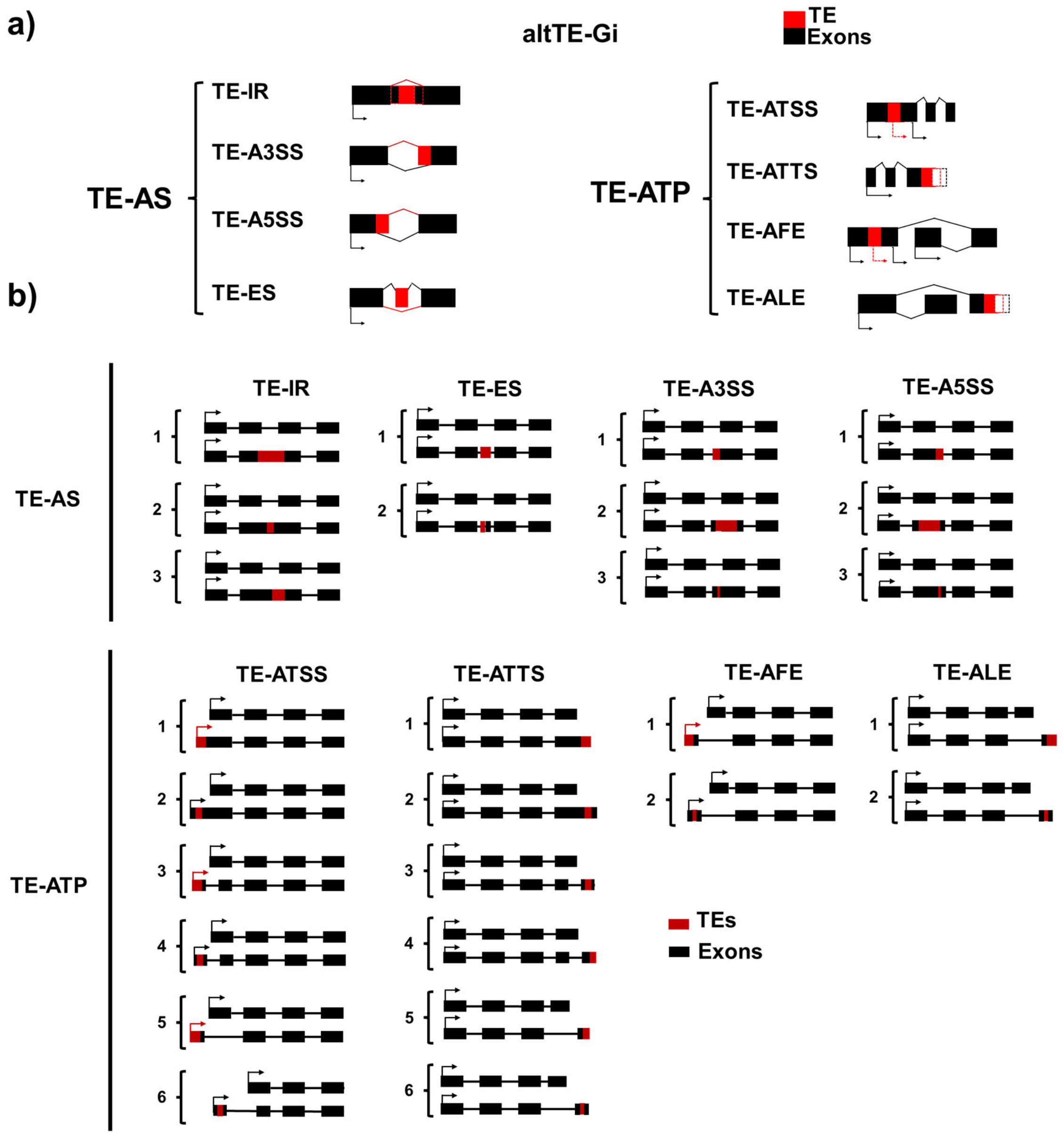


**Supplementary Note Figure 3:** a) Schematic illustration of TE-AS and TE-ATP events identified by ParasiTE. b) Illustration of altTE-Gi events considered as positive predictions by ParasiTE using the CATANA classification[8]. Several cases of positive predictions are displayed for each type of TE-AS and TE-ATP event.

Filtering used during step 5

Step 5 introduces potential false positive events because CATANA[8] may give unprecise predictions of AS or ATP events at the exonic level. Indeed, we noticed that CATANA may predict false positive AS or ATP events for an exon if a neighboring exon features these events. Moreover, an exon overlapping with a TE may be regarded as an AS or ATP event even if it is not directly related to the TE (Supplementary Note Figure 4). This may introduce false positives during the altTE-Gi prediction process by ParasiTE.

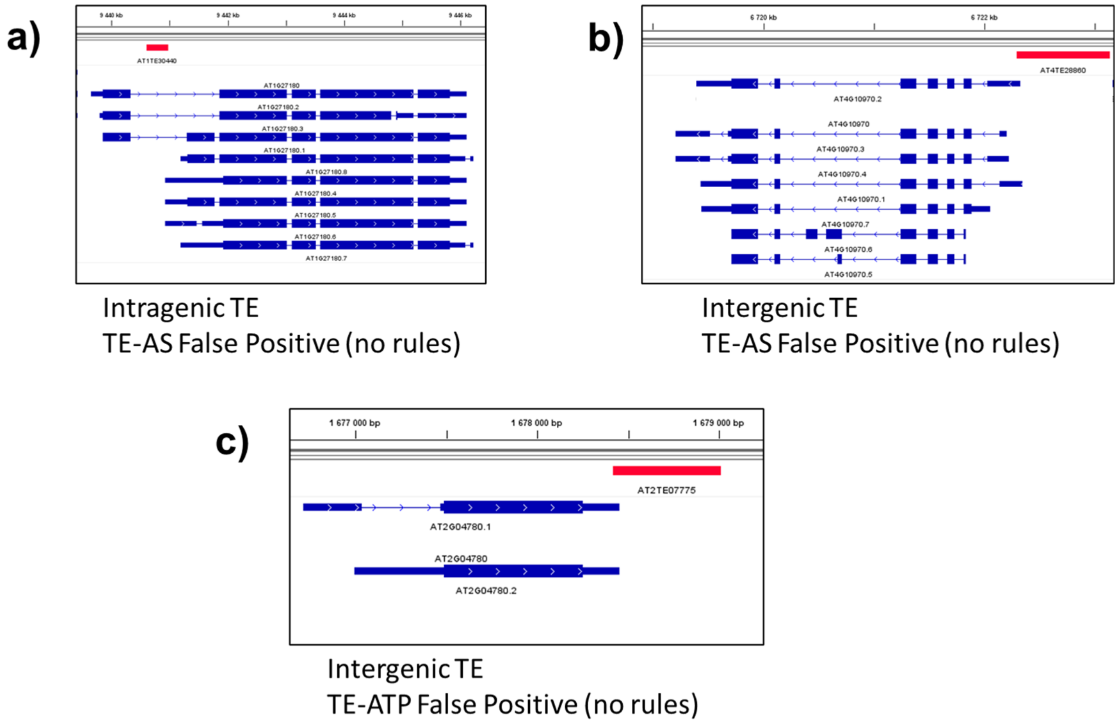


**Supplementary Note Figure 4**: Examples of false positive predictions of TE-AS or TE-ATP events without the application of filtering rules. a) and b) False positive TE-AS prediction by ParasiTE (without filtering rules). CATANA found an IR event for the nearby exon that was not related to the TE sequence. c) False positive TE-ATP prediction (TE-ATSS) by ParasiTE (without filtering rules). CATANA found an ATSS event for the overlapping exons, because of the neighboring exon features this event, but this ATSS event was not related to the TE sequence.

Therefore, several filtering rules were further implemented in ParasiTE to minimize false positive predictions of TE-AS or TE-ATP events. These rules use splicing site information of transcripts retrieved using the "extract_splice_sites.py" script from Hisat2[11]. ParasiTE also uses information of the location of the TE (intragenic or intergenic) to check for overlapping between the TE and edge of the associated exon and the frequency with which the TE overlaps transcripts of the same associated gene (noted “F”). Rules are illustrated below (Supplementary Note Figure 5-11).

**Rule 1:** A TE-AS event(s) is/are not predicted for a TE that does not overlap any splicing site(s) and that overlaps all isoforms of the gene (i.e., frequency of overlapping with isoforms of the associated gene is 100%, F = 1; Supplementary Note Figure 5).

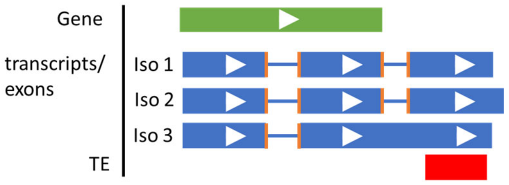


**Supplementary Note Figure 5:** Example of rule 1. Orange lines represent splicing sites.

**Rule 2:** A TE-ATP event(s) is/are not predicted for an intragenic TE that overlaps with all isoforms of the gene (F = 1), is located at the first or last exon of the transcript, and does not overlap with the edge of the associated exon (Supplementary Note Figure 6).

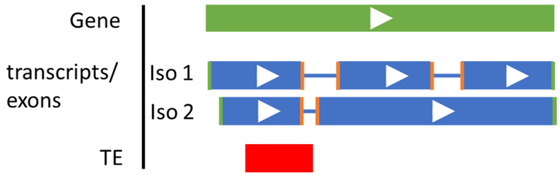


**Supplementary Note Figure 6:** Example of rule 2. Orange lines represent spicing sites and green lines represent the edge of the exon.

**Rule 3:** A TE-ATP event(s) is/are not predicted for an intragenic TE that has a TE-AS prediction, does not overlap with a splicing site, and does not overlap with the edge of the overlapping exon(s) (Supplementary Note Figure 7).

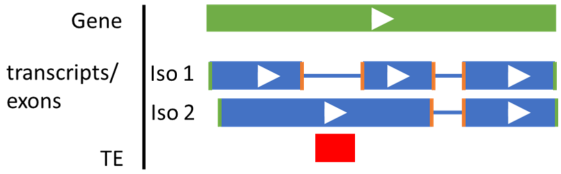


**Supplementary Note Figure 7:** Example of rule 3. Orange lines represent spicing sites and green lines represent the edge of the exon.

**Rule 4:** A TE-AS event(s) is/are not predicted for an intragenic TE that has a TE-ATP prediction, does not overlap with any splicing site, and overlaps with a single-exon transcript or the first or last exon of the associated transcript (Supplementary Note Figure 8)

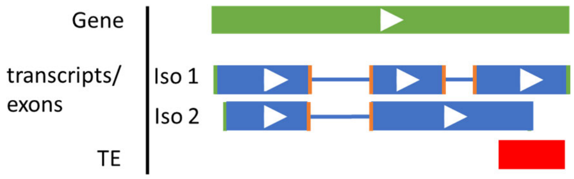


**Supplementary Note Figure 8:** Example of rule 4. Orange lines represent spicing sites and green lines represent the edge of the exon.

**Rule 5:** A TE-AS event(s) is/are not predicted for an intergenic TE which does not overlap with any splicing site and overlaps with the first or last exon of the associated transcript (Supplementary Note Figure 9).

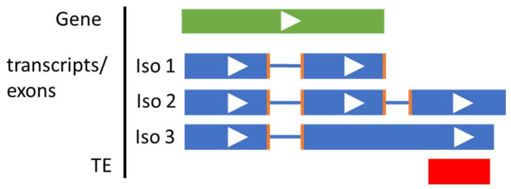


**Supplementary Note Figure 9:** Example of rule 5. Orange lines represent spicing sites and green lines represent the edge of the exon.

**Rule c1**: TE-A5SS and/or TE-A3SS is/are not predicted for a TE that is not overlapping with the right side of the associated exon (Supplementary Note Figure 10).

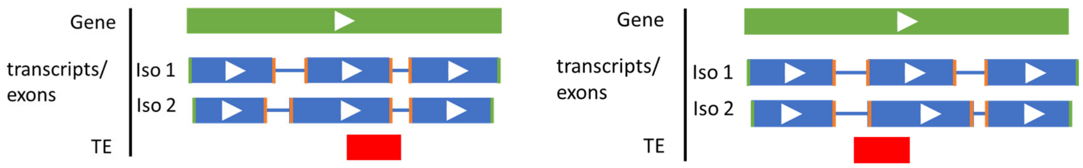


**Supplementary Note Figure 10:** Example of rule c1. Orange lines represent spicing sites and green lines represent the edge of the exon. Left: a false positive TE-A3SS event; right:

a false positive TE-A5SS event.

**Rule c2:** A TE-ATP event(s) is/are not predicted for a TE composed of one, two, or three exons and is not overlapping with the edge of the associated transcript (Supplementary Note Figure 11). This rule removes false positives displayed in Supplementary Note Figure 4c.

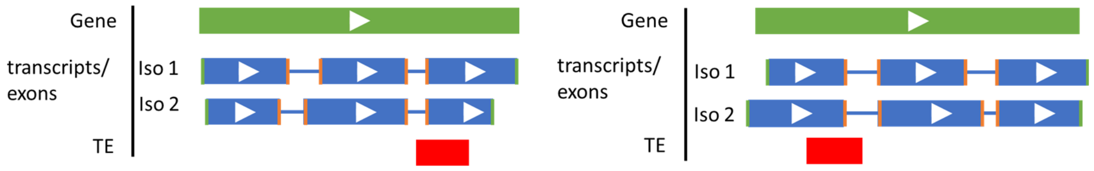


**Supplementary Note Figure 11:** Example of rule c2. Orange lines represent spicing sites and green lines represent the edge of the exon.

<u>Output of ParasiTE</u>

ParasiTE generates output files which list TE-Gt and altTE-Gi candidates at the exonic level (with transcript level information) and gene level.

Evaluation of the ParasiTE pipeline using *A. thaliana* gene annotation

We evaluated the accuracy of the ParasiTE pipeline with the Araport11 transcriptome dataset, using gene model and TE annotations, transcriptomic data, and gene-like TE annotation as described above. We first ran ParasiTE without applying any filtering rules (no rules) and manually checked the prediction of TE-AS and/or TE-ATP events for each TE-gene pair. We labeled as true positive or false positive the TE-AS or TE-ATP predictions of each TE-Gene pair. Moreover, because TE-Gene pair associated to altTE-Gi may be involved only in TE-AS but not in TE-ATP (or vice-versa), we also checked the "NA" predictions for TE-AS and TE-ATP (example of TE-AS: "NA" and TE-ATP: "ATTS" in Supplementary Note Figure 1). A correct "NA" prediction was labeled as true negative, but an incorrect "NA" prediction was labeled as false negative. In this analysis, we only checked if TEs were directly involved in TE-AS or TE-ATP events without checking the accuracy of the type of events (such as TE-IR or TE-A3SS). For altTE-Gi associated to intragenic TE we checked events found in chromosomes 1 to 5 (300 altTE-Gi events with no rules) of the *Arabidopsis* genome. For altTE-Gi associated to intergenic TEs we checked events found in chromosomes 1 and 2 (259 altTE-Gi events with no rules), but five altTE-Gi events were manually found associated to potential gene-like TEs and were removed, resulting in 254 altTE-Gi events (with no rules). Next, we applied the five filtering rules (see above) to the ParasiTE pipeline to minimize false positive predictions (Supplementary Note Figure 12). We obtained reasonably good positive predictive values (PPV) for TE-AS and TE-ATP by applying rules 1-5, which achieved a total (intergenic and intragenic TEs) PPV of ~78% for TE-AS and ~96% for TE-ATP events (Supplementary Note Figure 12a). Applying rules 1-5 did not affected the NPV of ~100% for TE-AS or TE-ATP events (Supplementary Note Figure 12b).

a)

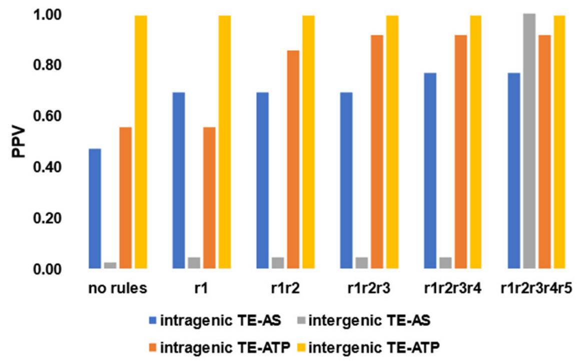


b)

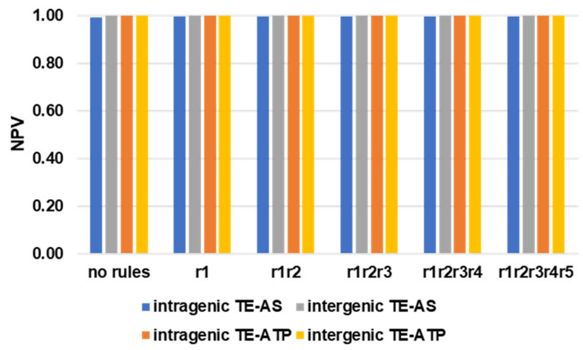


**Supplementary Note Figure 12:** a) Positive predictive values (PPV) and b) Negative predicted values obtained by ParasiTE with the Araport11 annotation. Filtering rules (from no rules to rules 1, 12, 123, 1234, and 12345) improved the detection accuracy of altTE-Gi events.

Next, we further evaluated the tool's accuracy for predicting TE-gene isoform formation events by distinct molecular mechanisms (Supplementary Note Figure 3a). TE-AS events can be further classified into splicing events, including TE-IR, TE-ES, TE-A5SS, and TE-TE-A3SS, while TE-ATP events can be further classified into TE-ATSS, TE-ATTS, TE-AFE, and TE-ALE. For this analysis, predictions for potential gene-like TEs were not removed in order to measure their impact on the tool's accuracy.

After manual inspection of predicted TE-AS events (n = 144 after c1c2) in chromosomes 1 and 2 of the *Arabidopsis* genome, we found that 60% of TE-ES (n = 10 after c1c2), 77% of TE-IR (n = 88 after c1c2), 70% of A5SS (n = 20 after c1c2), and 46% of A3SS (n = 26 after c1c2) events were correctly predicted (Supplementary Note Figure 13). In addition, we found that 96% of ATSS (n = 150 after c1c2), 94% of ATTS (n = 148 after c1c2), 93% of AFE (n = 43 after c1c2), and 87% of ALE (n = 15 after c1c2) events were correctly predicted (Supplementary Note Figure 13). Overall, the ParasiTE pipeline can reasonably identify and classify altTE-Gi formation events.

a)

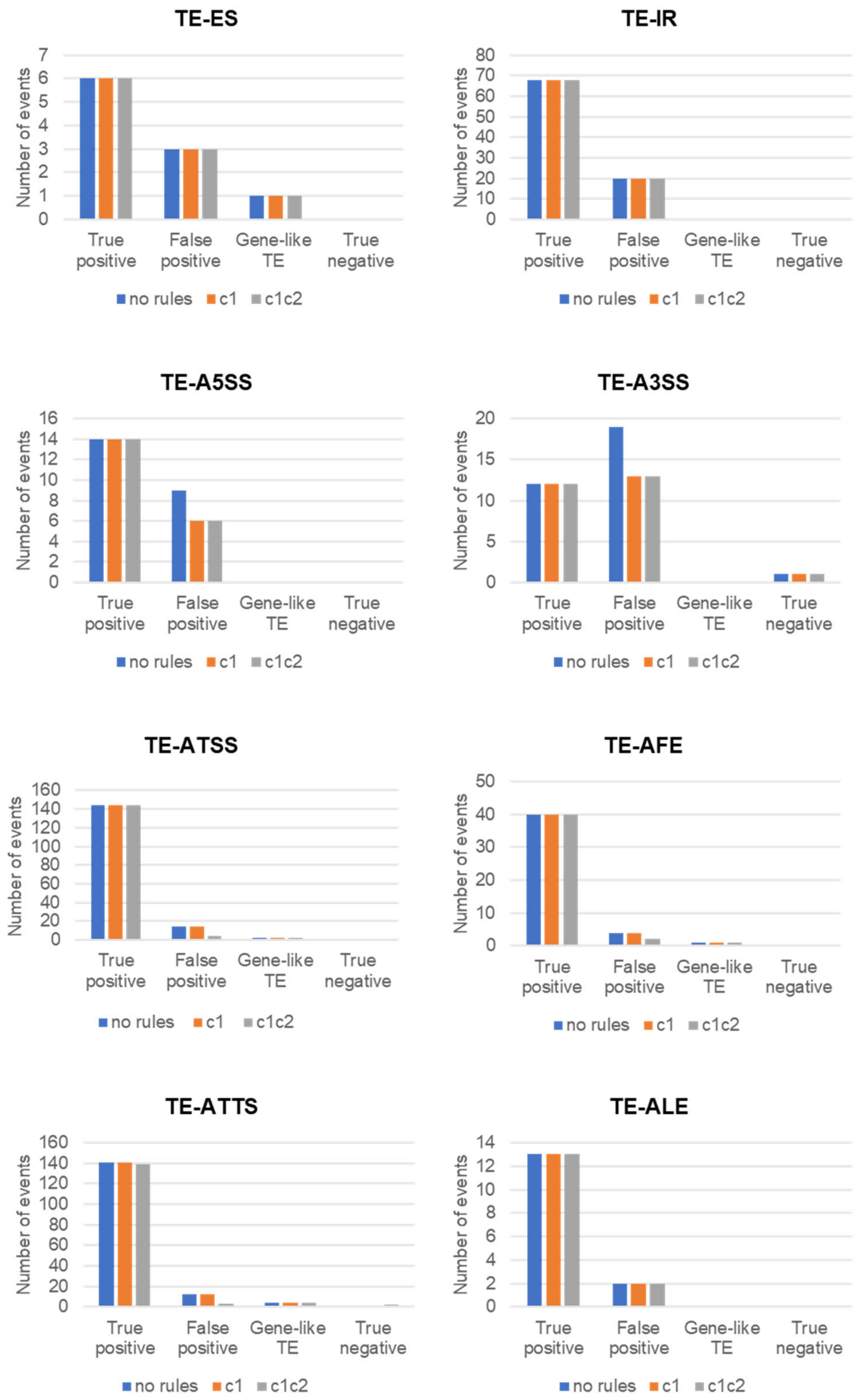
TE-ES
Number of events
7
6
5
4
3
2
1
0
True positive
False positive
Gene-like TE
True negative
no rules
c1
c1c2
TE-IR
Number of events
80
70
60
50
40
30
20
10
0
True positive
False positive
Gene-like TE
True negative
no rules
c1
c1c2
TE-A5SS
Number of events
16
14
12
10
8
6
4
2
0
True positive
False positive
Gene-like TE
True negative
no rules
c1
c1c2
TE-A3SS
Number of events
20
15
10
5
0
True positive
False positive
Gene-like TE
True negative
no rules
c1
c1c2
TE-ATSS
Number of events
160
140
120
100
80
60
40
20
0
True positive
False positive
Gene-like TE
True negative
no rules
c1
c1c2
TE-AFE
Number of events
50
40
30
20
10
0
True positive
False positive
Gene-like TE
True negative
no rules
c1
c1c2
TE-ATTS
Number of events
160
140
120
100
80
60
40
20
0
True positive
False positive
Gene-like TE
True negative
no rules
c1
c1c2
TE-ALE
Number of events
14
12
10
8
6
4
2
0
True positive
False positive
Gene-like TE
True negative
no rules
c1
c1c2

b)

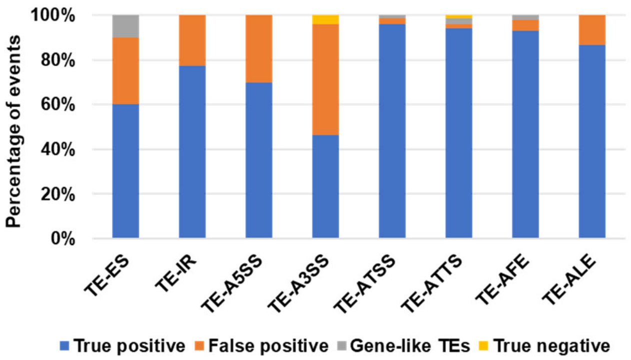


**Supplementary Note Figure 13:** a) Evaluation of prediction efficiency of distinct TE-AS and TE-ATP events obtained by ParasiTE with the Araport11 annotation and either with (rules c1 and c2) or without (no rules) correction rules. b) Evaluation of the efficiency of ParasiTE to predict TEs involved in altTE-Gi (TE-AS or TE-ATP) events.

We also evaluated the efficiency of the removal of gene-like TEs in step 1 of the ParasiTE pipeline (see Supplementary Note Figure 14) using an optional "gene-like TE annotation". A comparison of ParasiTE outputs obtained from the Col-0 DRS and *ddm1*-DRS data (which contains many activated "free" gene-like TE transcripts) showed that the step greatly reduced false positive predictions of TE-Gt events and improved the prediction of true events (Supplementary Note Figure 14). In the *ddm1*-DRS dataset, numerous TE transcripts are included, which increased the number of false positive detections of TE-AS and TE-ATP events. The optional filtering step using a "gene-like TEs annotation" helped to reduce this number (orange bars).

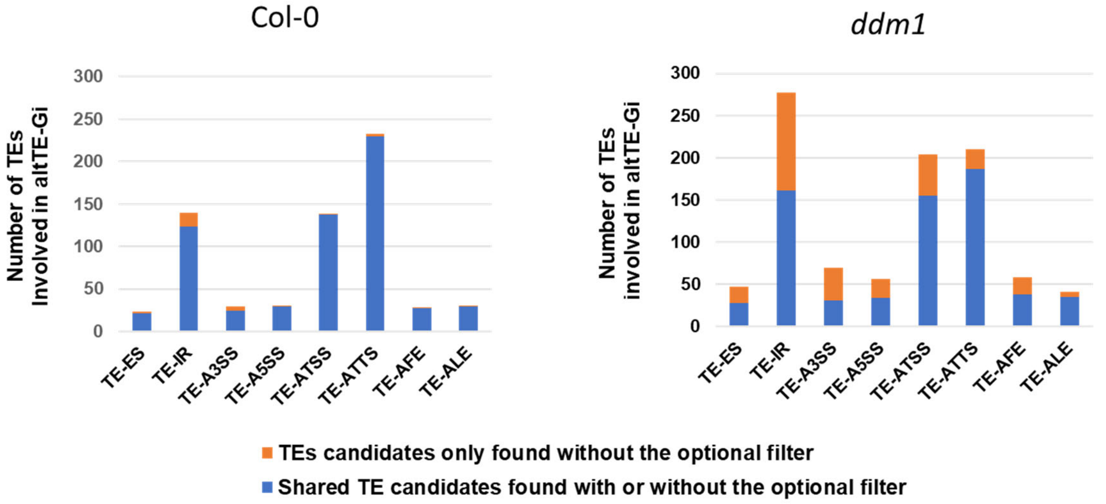


**Supplementary Note Figure 14:** Number of TEs involved in altTE-Gi events found by ParasiTE using the *de novo* DRS transcriptome of Col-0 and *ddm1* (Stringtie2 "-L mode" and Araport11 as the reference). Blue: number of TEs predicted using the optional filter with a TE-gene like annotation. Orange: additional TEs predicted as being involved in altTE-Gi events without the filter; these correspond to gene-like TEs and are false positive predictions. Many gene-like TEs were retrieved in *ddm1* because of the lack of DNA methylation, causing TE re-activation.

Finally, ParasiTE can estimate the contribution of TE-Gts and altTE-Gis to CDS or 5′/3′-UTRs. For each TE-Gt event, ParasiTE uses bedtools' intersect function to overlap each exonic region of TEs involved in TE-Gts to the CDS and 5'/3'-UTRs of genes. Features per gene can be counted, as demonstrated in Supplementary Note Figure 15**.**

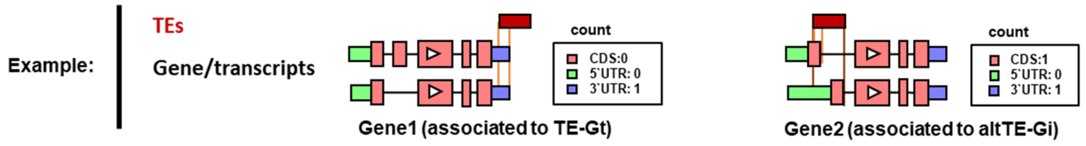


**Supplementary Note Figure 15:** Example of ParasiTE's counting feature.
