## Supplementary Information for "Long-read direct RNA sequencing reveals epigenetic regulation of chimeric gene-transposon transcripts in *Arabidopsis thaliana*"

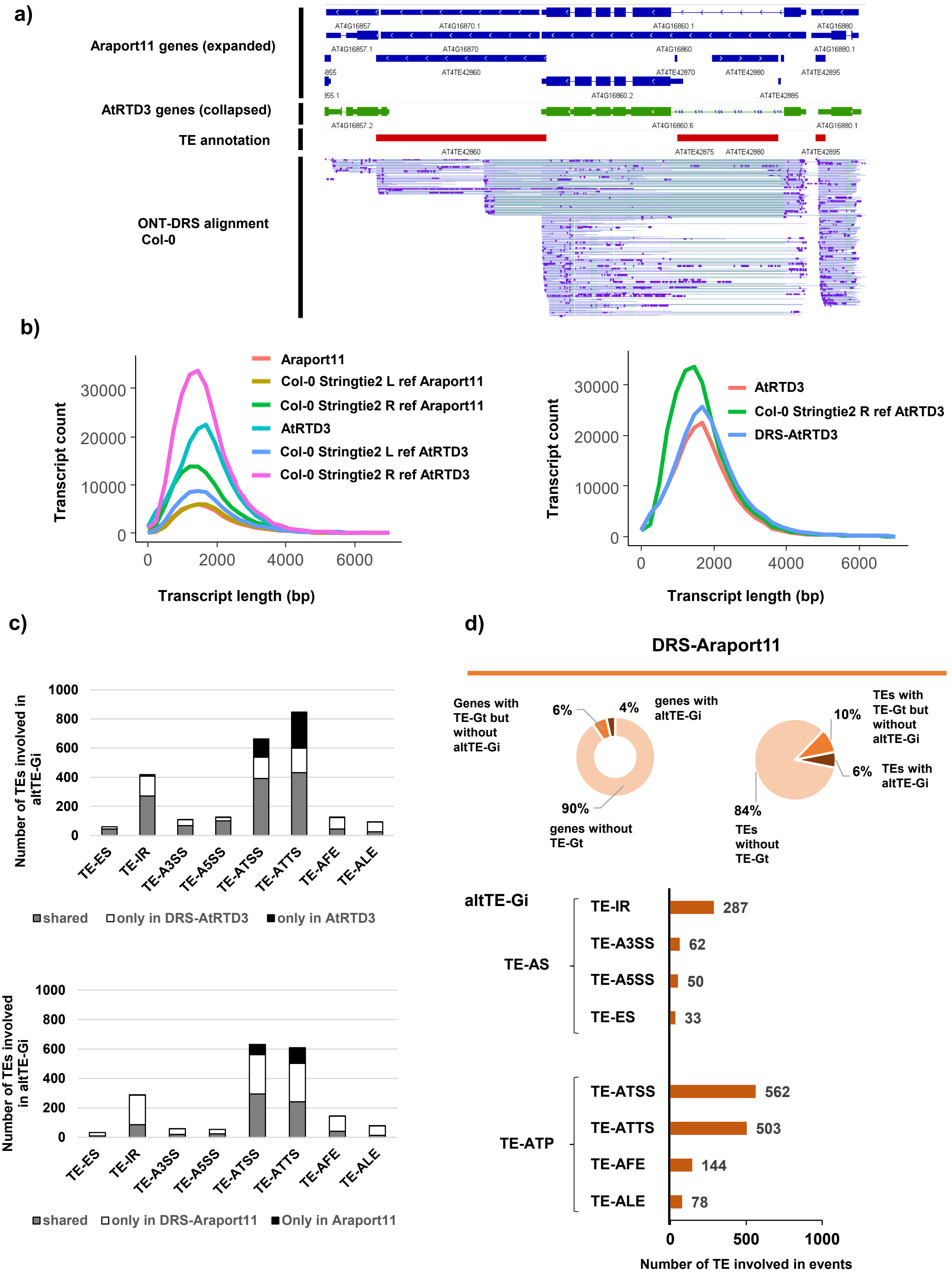


**Supplementary Fig. 1.** Detection of TE-Gts and altTE-Gis in the *Arabidopsis* transcriptome. **a)** A representative TE-Gt associated with the *RPP4-ATCOPIA4* locus. ONT-DRS reads were aligned to the TAIR10 genome assembly of *A. thaliana*. **b)** DRS transcriptome assembly. Left: the number and length of transcripts obtained from *de novo* assembly of Col-0 ONT-DRS data with “L” or “R” mode in Stringtie2 using Araport11 or AtRTD3 as references. Right: DRS-AtRTD3 transcriptome contains longer and a higher number of transcripts than the original AtRTD3 data. The "Col-0 Stringtie2 R ref AtRTD3" transcriptome contains more transcripts and are, on average, shorter than those from DRS-AtRTD3, reflecting fragmentation of the transcriptome. **c)** Number of TEs involved in altTE-Gi according to the RNA processing events, detected by ParasiTE using AtRTD3, Araport11, DRS-AtRTD3, or DRS-Araport11 as a reference. **d)** Top: Number of genes (left) and TEs (right) associated with TE-Gts and altTE-Gis identified in DRS-Araport11. Bottom: RNA processing events associated with altTE-Gis (TE-AS and TE-ATP), and the number of events identified by ParasiTE with DRS-Araport11.

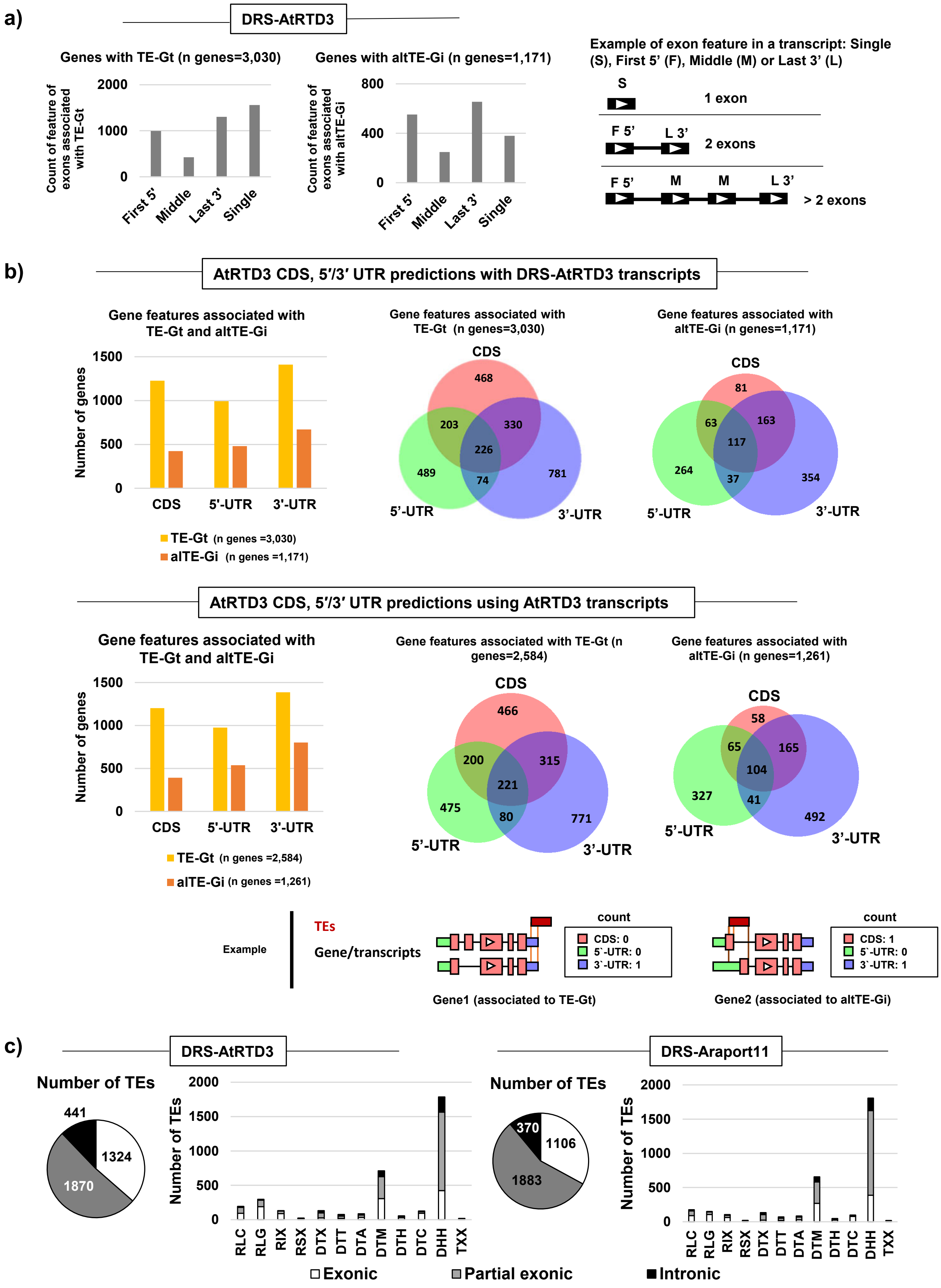


**Supplementary Fig. 2**. Profile of TEs involved in TE-Gts and altTE-Gis. **a)** Exonic positions of TEs identified in TE-Gts or altTE-Gis. Definitions for the features are displayed in the right panel. **b)** Contribution of TEs in TE-Gts or altTE-Gis to CDS, 5′/3′-UTRs. CDS, 5′/3′-UTRs in DRS-AtRTD3 and AtRTD3 were predicted using annotations from the AtRTD3 dataset. Examples of the counting of features are displayed in the bottom panels. **c)** Number of exonic, partial exonic, and intronic TEs found by ParasiTE in DRS-AtRTD3 and DRS-Araport11. See the Supplementary Notes for details on the criteria of the classes.

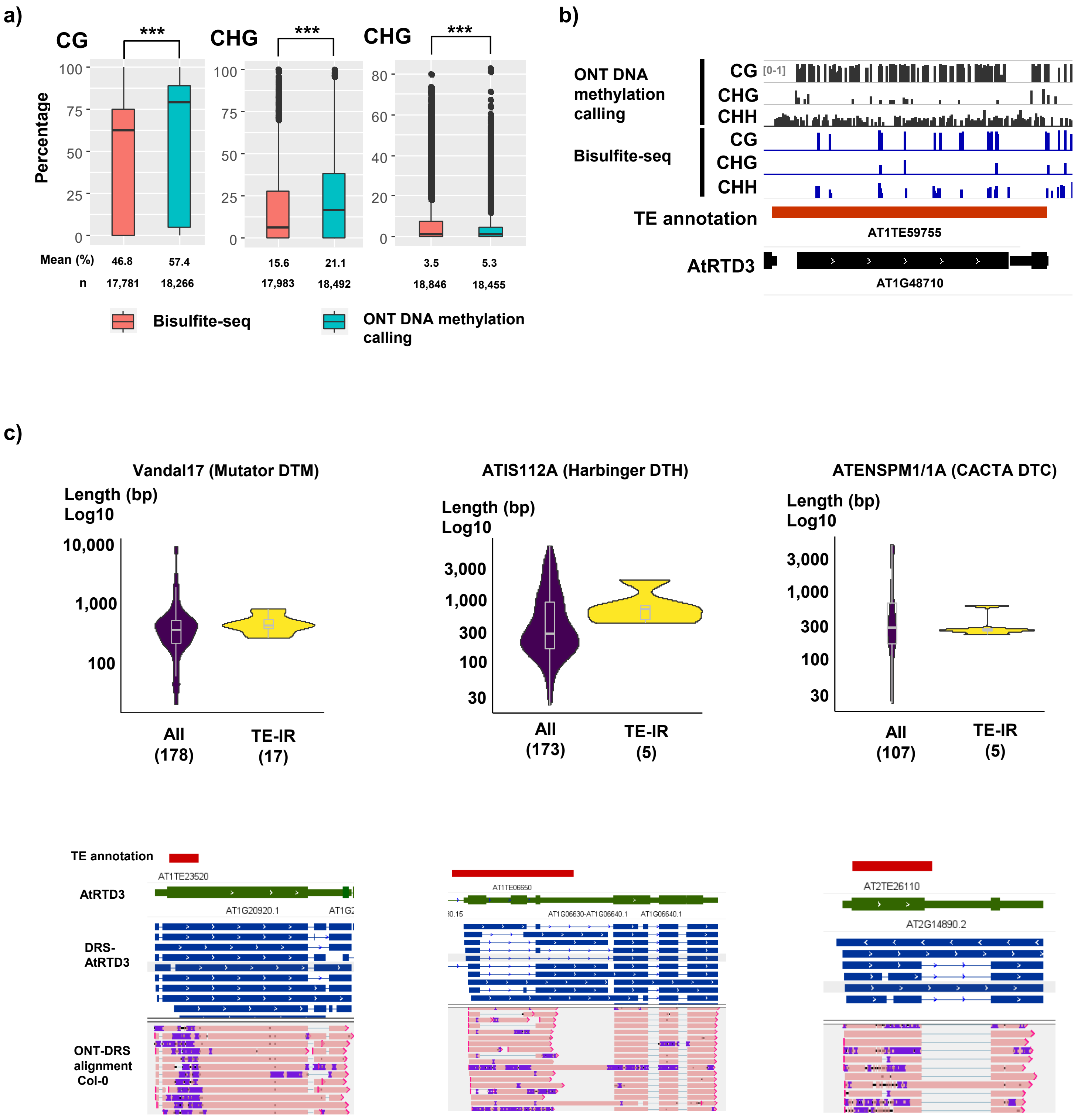


**Supplementary Fig. 3**: ONT methylation calling of TEs and the profiles of TE superfamilies enriched in TE-IR events. **a)** Comparison of methylated CG, CHG, and CHH in TEs obtained by Bisulfite-Seq[90] and ONT DNA methylation calling. ***, $p$<0.001 by the Mann–Whitney $U$ test. **b)** Genome browser tracks showing a comparison of DNA methylation calling obtained from Bisulfite-Seq and ONT DNA methylation calling at the *LTR/ATCOPIA78* (*ONSEN*) (*AT1TE59755*) locus. **c)** Length of TE superfamilies showing enrichment in TE-IR events. Genome browser tracks show representative loci associated with TE-IR events with the TE superfamilies.

a)

b)

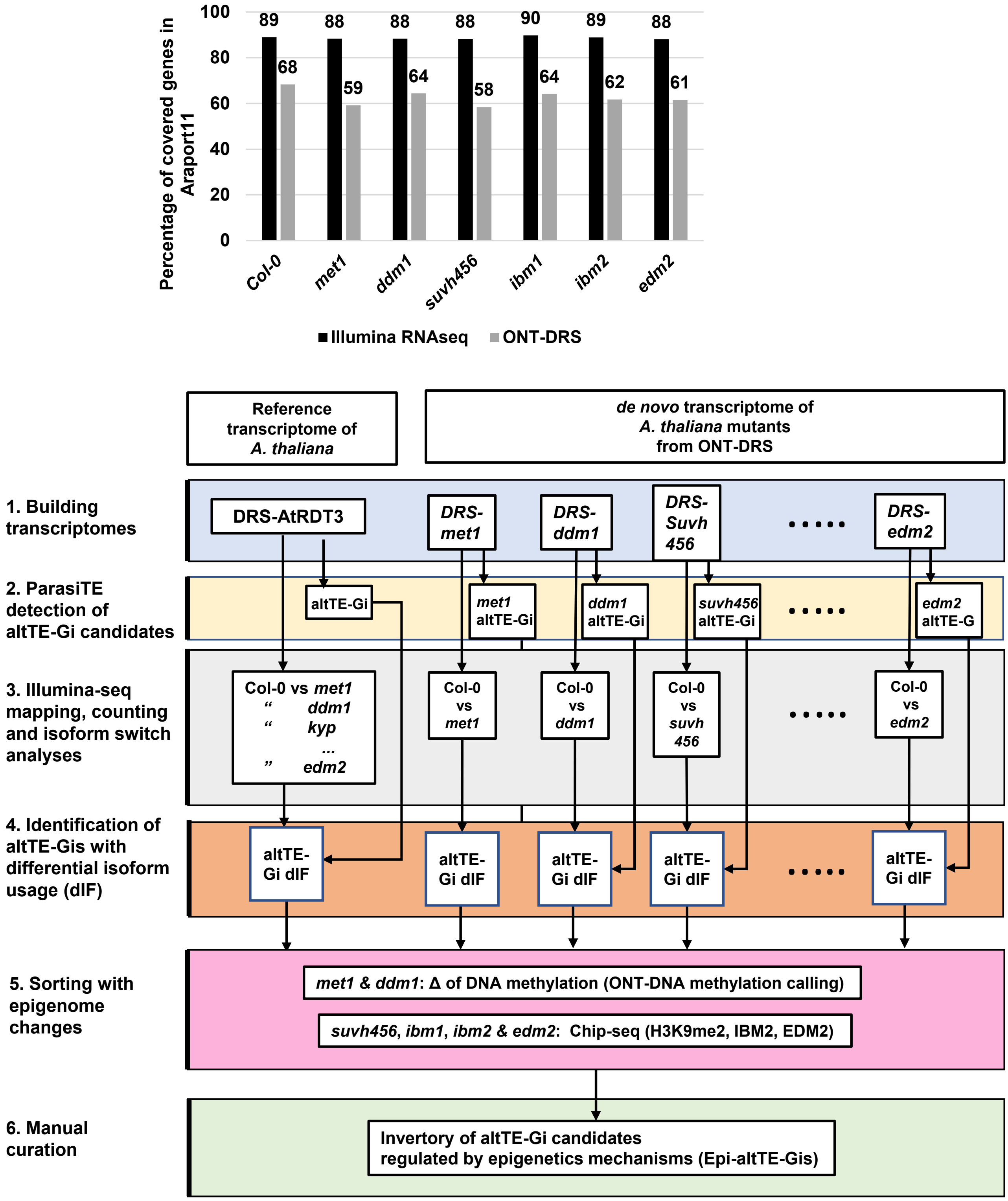


**Supplementary Fig. 4.** ONT-DRS sequencing of epigenetic mutants and identification of Epi-altTE-Gis. **a)** Coverage of Araport11 genes by at least 10 reads with ONT-DRS and Illumina RNA-seq in Col-0 and epigenetic mutants. **b)** Scheme of the methodology used to detect Epi-altTE-Gi: 1) DRS-AtRTD3 was obtained by merging the AtRTD3 transcriptome annotation and the new DRS-Col-0 transcriptome annotation. In parallel, *de novo* transcriptome assembly of epigenetic mutants was built from the ONT-DRS data; 2) ParasiTE detected altTE-Gi candidates in the DRS-AtRTD3 and mutant-DRS datasets; 3) and 4) Illumina RNA-seq data of Col-0 and epigenetic mutants were aligned to the DRS-AtRTD3 transcriptome assembly. Next, altTE-Gis in DRS-AtRTD3 displaying significant isoform switch events were extracted. Following this, Illumina RNA-seq data of Col-0 and mutants were also aligned to the mutant-DRS datasets. Next, altTE-Gis in mutant-DRS associated with significant isoform switch events were retrieved; 5) The direct effects of epigenome changes on Epi-altTE-Gis were verified. Epi-altTE-Gis associated with DNA methylation changes in *met1* and *ddm1*, H3K9me2 ChIP-seq peaks in Col-0 for *suvh456,* H3K9me2 ChIP-seq peaks in Col-0 for *ibm1*, and IBM2/EDM2 ChIP-seq peaks for *ibm2* and *edm2* were extracted.

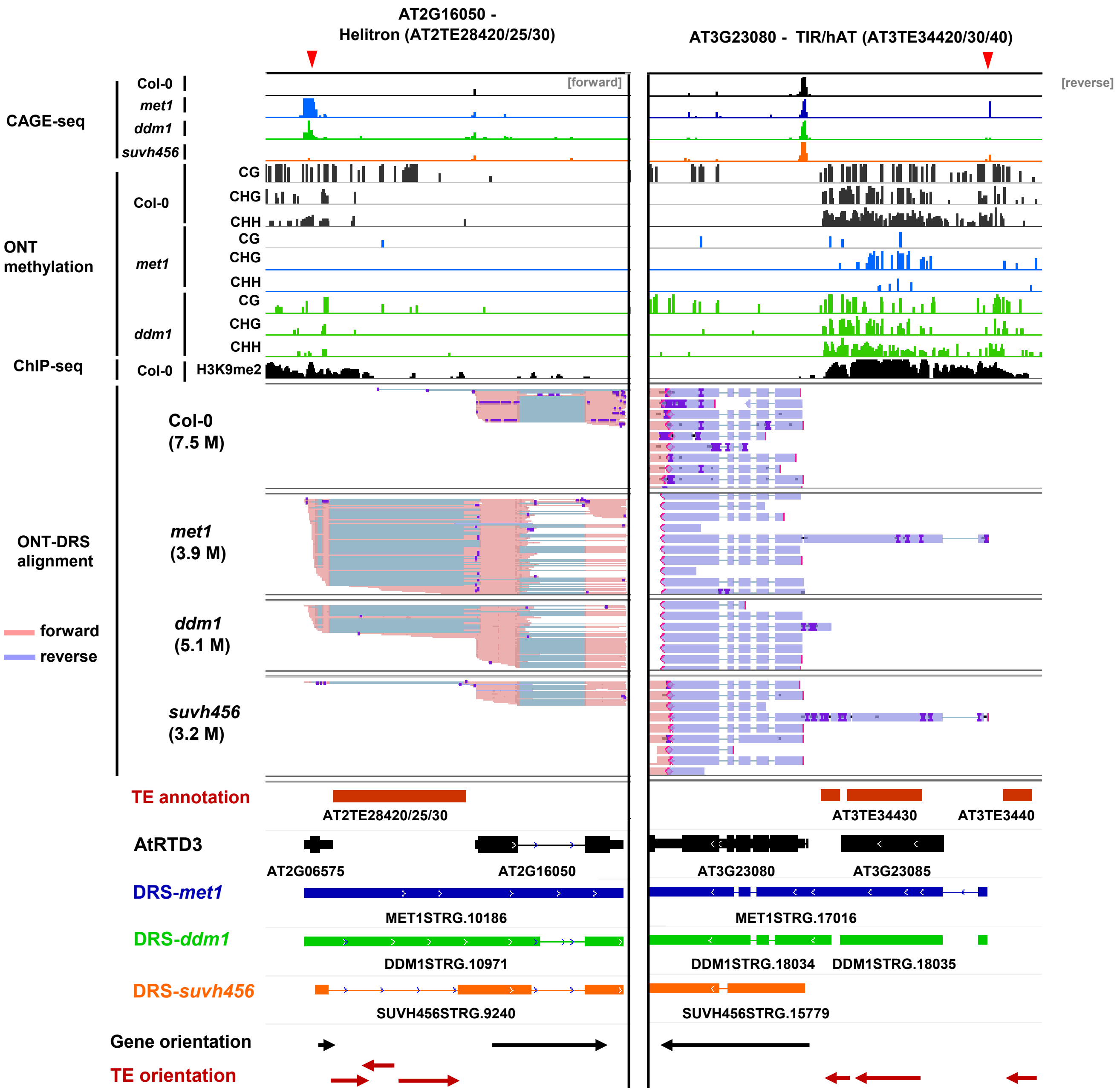


**Supplementary Fig. 5.** Epigenetic regulation of TE-ATSS. Representative genome loci showing Epi-altTE-Gis with TE-ATSS events. Tracks (from top to bottom): CAGE-seq (reads per million; 0–1. Only forward or reverse strands are shown); methylation level of each mutant in CG, CHG, and CHH contexts (0–100%); Col-0 ChIP-seq of H3K9me2 (reads per million; 0–1); DRS read alignments of Col-0 and indicated mutants; TE and AtRTD3 transcript annotations, *de novo* assembly of transcripts in mutants, and the orientation of genes and TEs. Red arrows indicate cryptic TSSs detected in the epigenetic mutants.

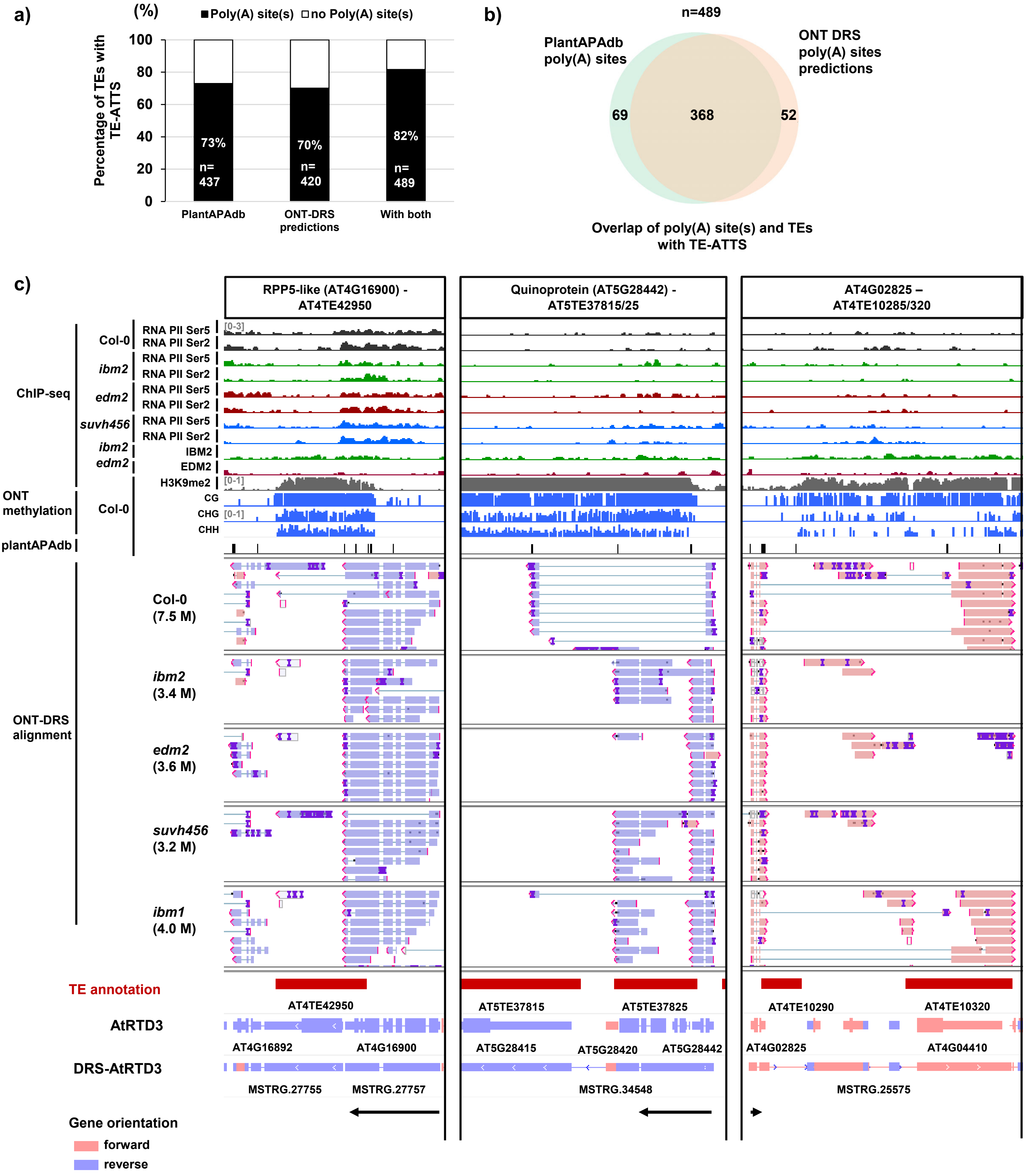


**Supplementary Fig. 6. a)** TEs with TE-ATTS events that overlap with poly(A) sites in the PlantAPA database or detected in the DRS data. **b)** Overlap of poly(A) sites with TEs associated with ATTS. **c)** Epigenetic regulation of TE-ATTS. Representative genome loci showing Epi-altTE-Gi with TE-ATTS events. Tracks (from top to bottom): ChIP-seq data for RNA Pol II phosphorylated at Ser5/Ser2 in CTD repeats (bins per million); ChIP-seq data for IBM2 and EDM2 localization (bin per million); Col-0 ChIP-seq of H3K9me2 (reads per million); methylation levels of Col-0 in CG, CHG, and CHH contexts (0–100%); poly(A) sites obtained from the PlantAPA database; DRS read alignments of Col-0 and indicated mutants; TE and transcript annotations of AtRTD3 and DRS-AtRTD3 in this study and the orientation of genes.

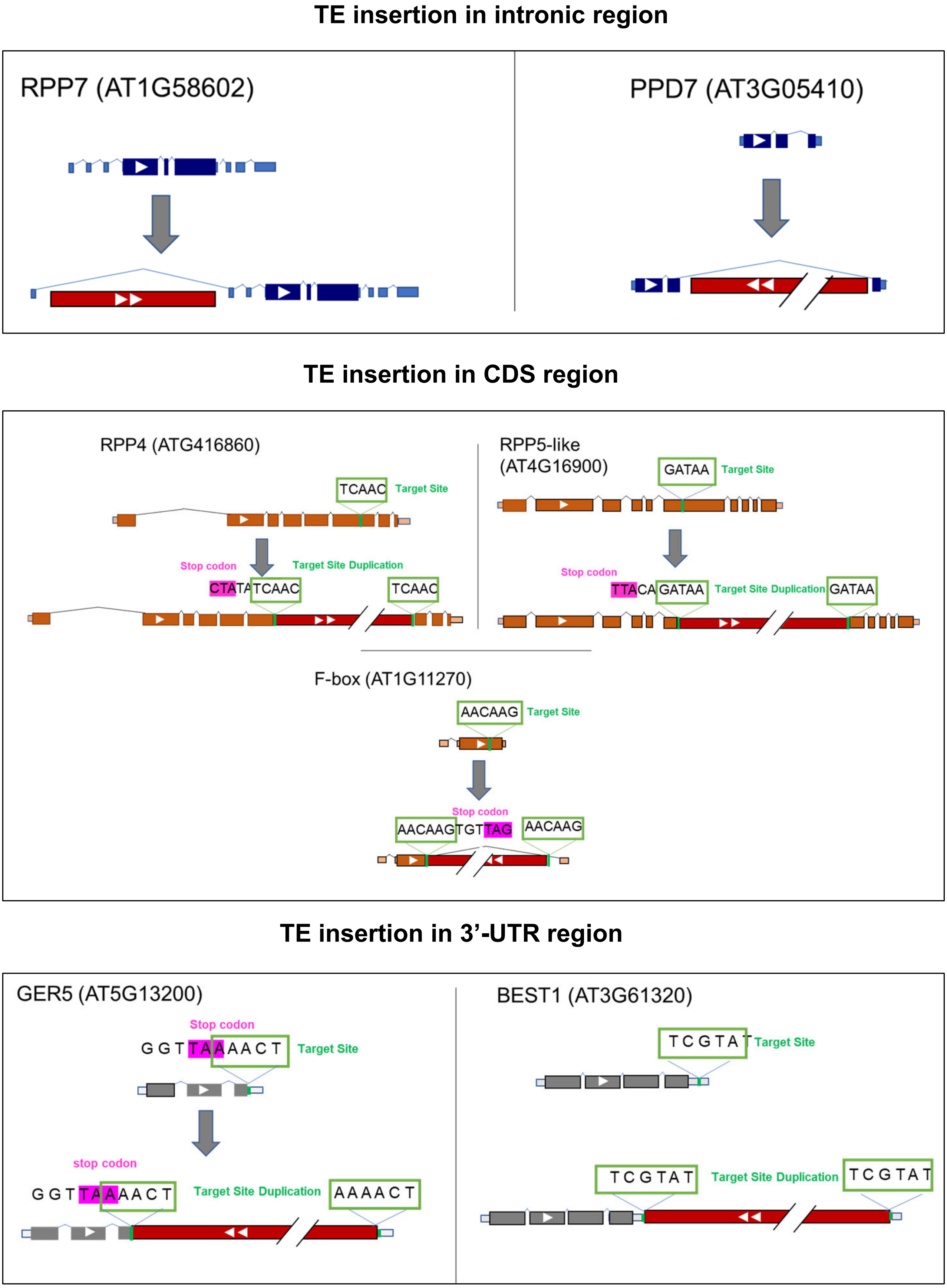


**Supplementary Fig. 7.** Insertion of TEs in intragenic regions (intron, CDS/exon, and 3′-UTR) of genes analyzed in this study. Target site duplications are indicated. For *PPD7*, *GER5*, and *BEST1*, the Ler-0 genome sequence was used as an ancestral reference ecotype before TE insertion.

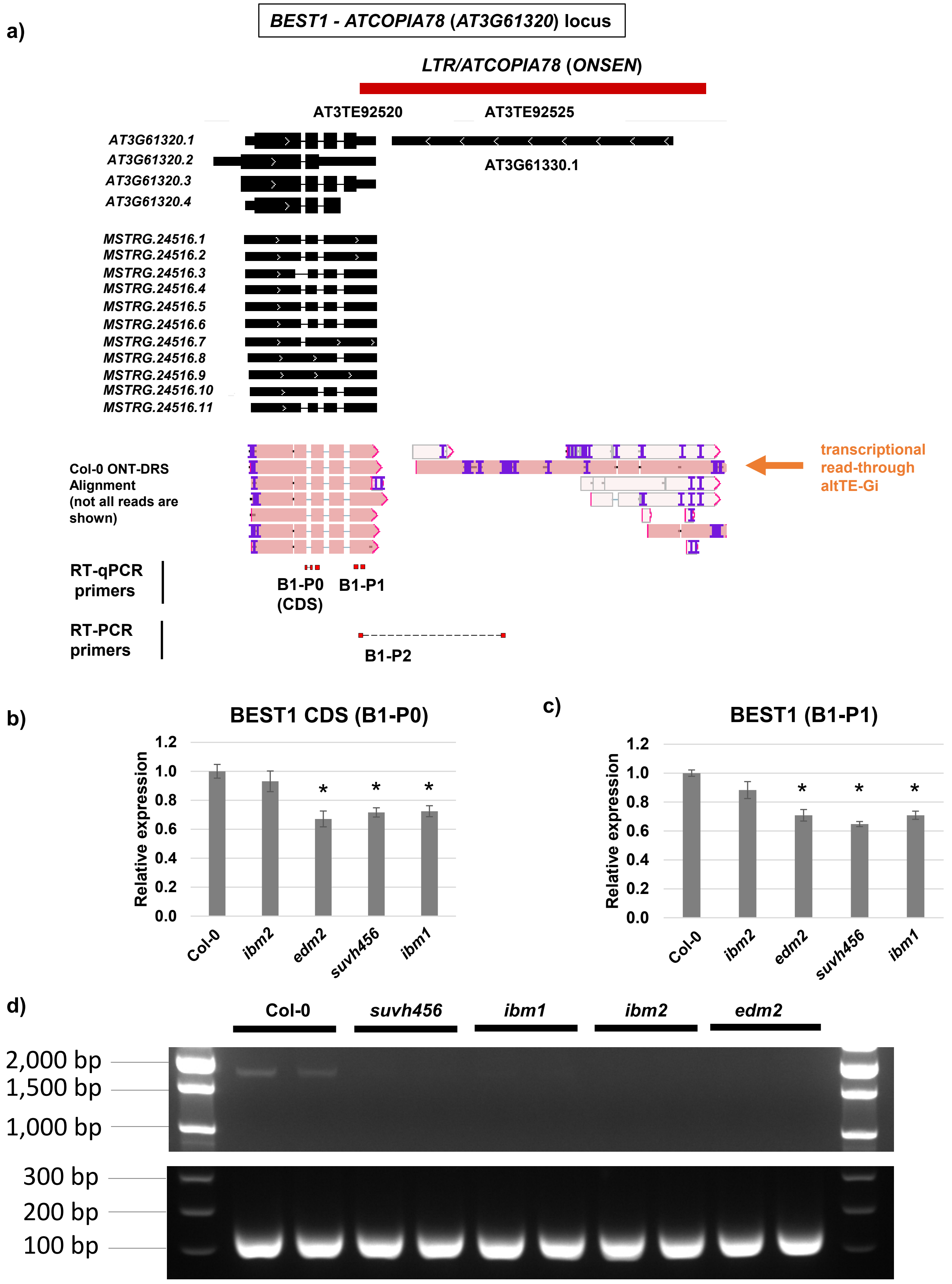


**Supplementary Fig. 8.** *BEST1-ATCOPIA78* (*ONSEN*) (*AT3G61320*) locus. **a)** Genome browser tracks showing the complete DRS-AtRTD3 transcript annotation. The Col-0 ONT-DRS alignment is also shown with a transcriptional read-through altTE-Gi detected in the Col-0-DRS data (orange arrow). Positions of primers used for RT-qPCR in **b** and **c** and RT-PCR in **d** are indicated. **b), c)** Relative expression of *BEST1* CDS and altTE-Gis in Col-0 and the epigenetic mutants. Bars represent the means of four biological replicates ± SEM. *, $p < 0.05$ by $t$-test.
**d)** Agarose gel electrophoresis of RT-PCR amplicons with primers targeting the predicted transcriptional read-through of *BEST1*. *ACT2* (*AT3G18780*) was used as a positive control.

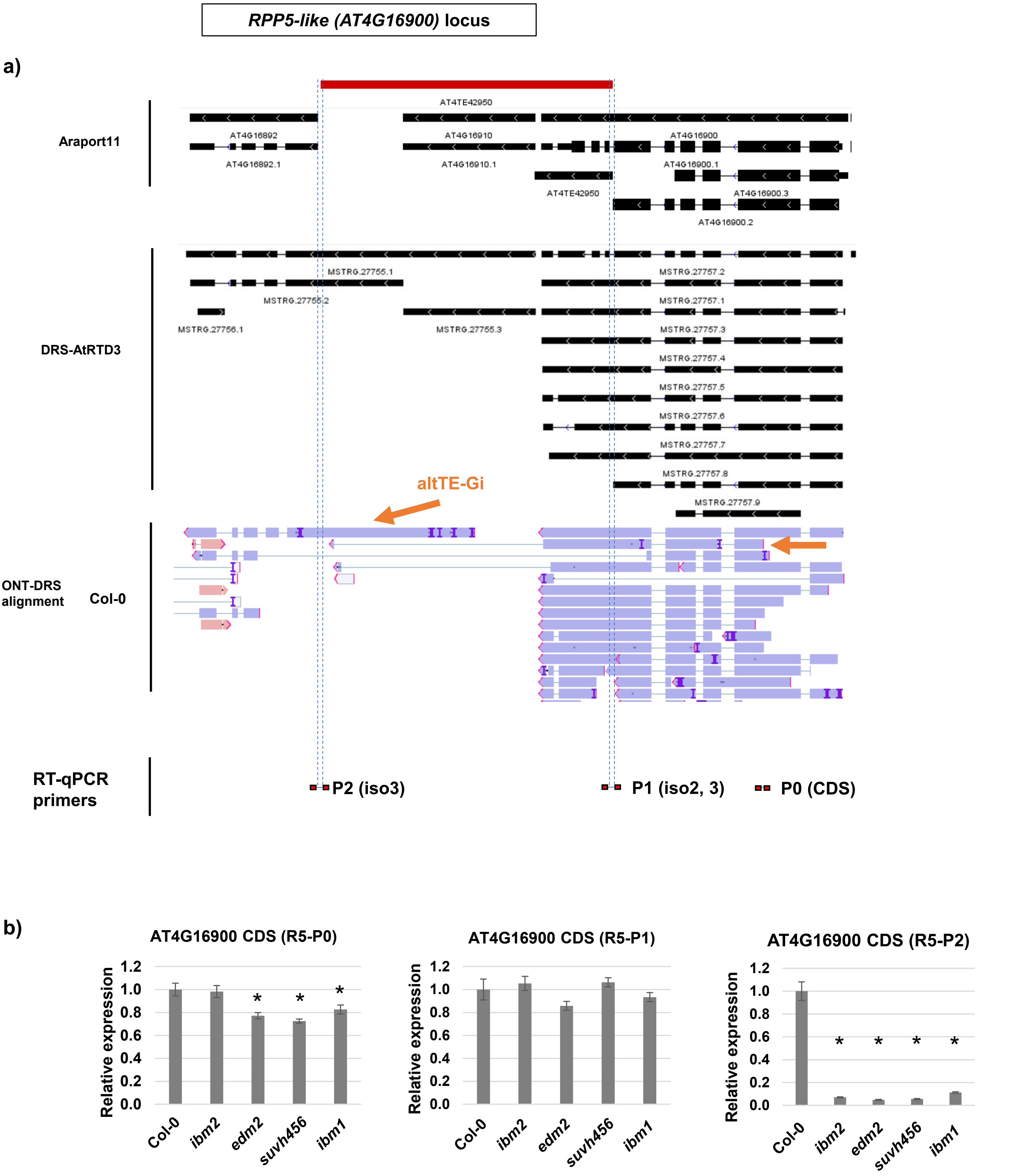


**Supplementary Fig. 9.** The *RPP5-like-AT4TE42950* (*AT4G16900*) locus. **a)** Genome browser tracks showing the complete DRS-AtRTD3 transcript annotation, DRS reads, and primer positions for RT-qPCR in **b**. Arrows indicate altTE-Gis covering the downstream *AT4G16892* locus. **b)** Relative expression of the *RPP5-like* CDS and altTE-Gis in Col-0 and epigenetic mutants. Target-specific primers for RT-qPCR are illustrated above. Bars represent the means of four biological replicates ± SEM. *, $p < 0.05$ by *t*-test.

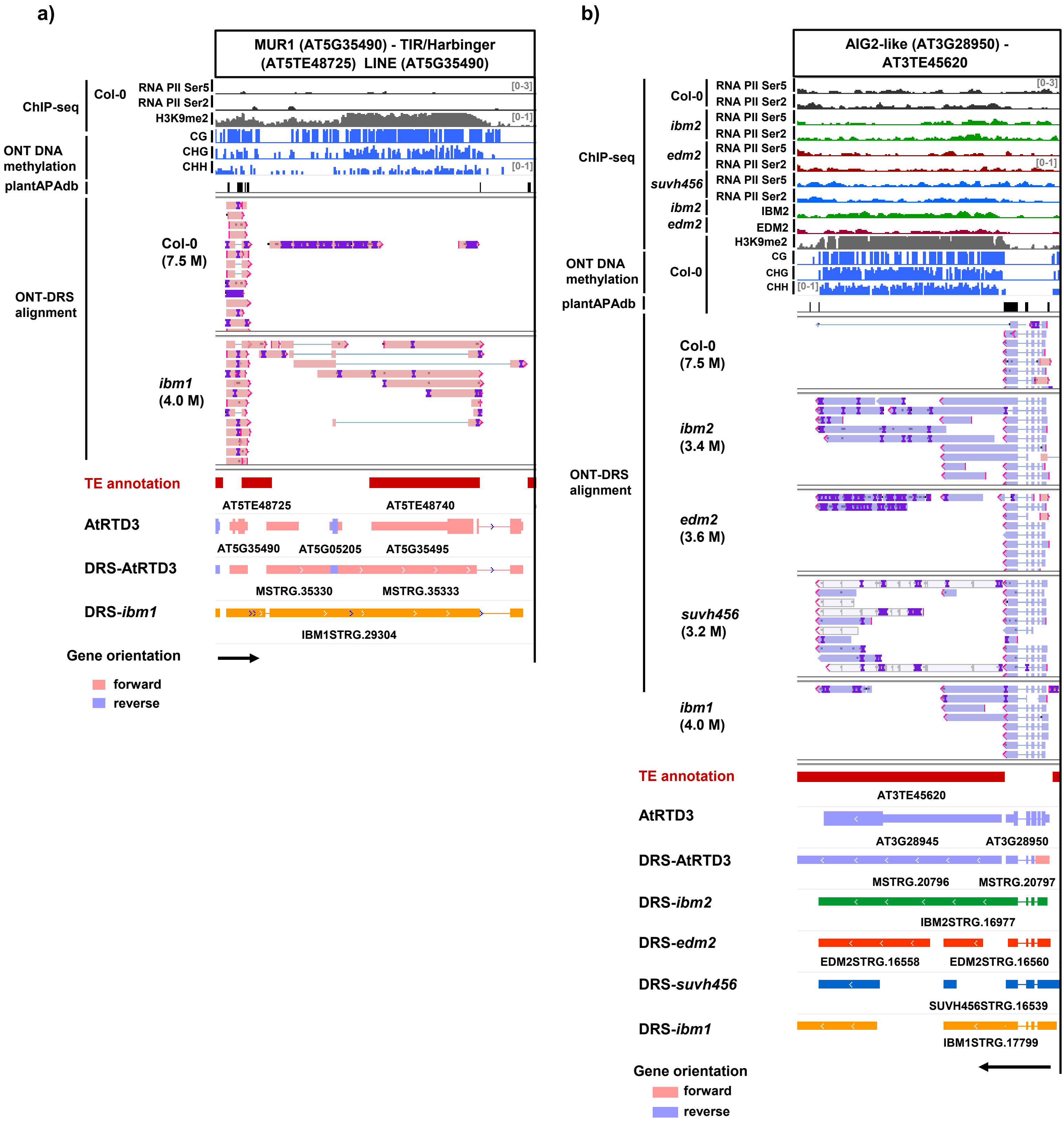


**Supplementary Fig. 10. a)** *MUR1* (*AT5G35490*) and **b)** *AIG2-like* (*AT3G28950*) gene loci showing Epi-altTE-Gi production with TE-ATTS events, which were particularly enhanced in the mutant background. Tracks (from top to bottom): ChIP-seq data for RNA Pol II phosphorylated at Ser5/Ser2 in CTD repeats (bin per million); Col-0 ChIP-seq of H3K9me2 (reads per million); methylation levels of Col-0 in CG, CHG, and CHH contexts (0–100%); poly(A) sites obtained from the PlantAPA database; DRS read alignments of Col-0 and indicated mutants; TE and transcript annotations of AtRTD3, DRS-AtRTD3, and mutant-DRS in this study and the orientation of genes.

a)

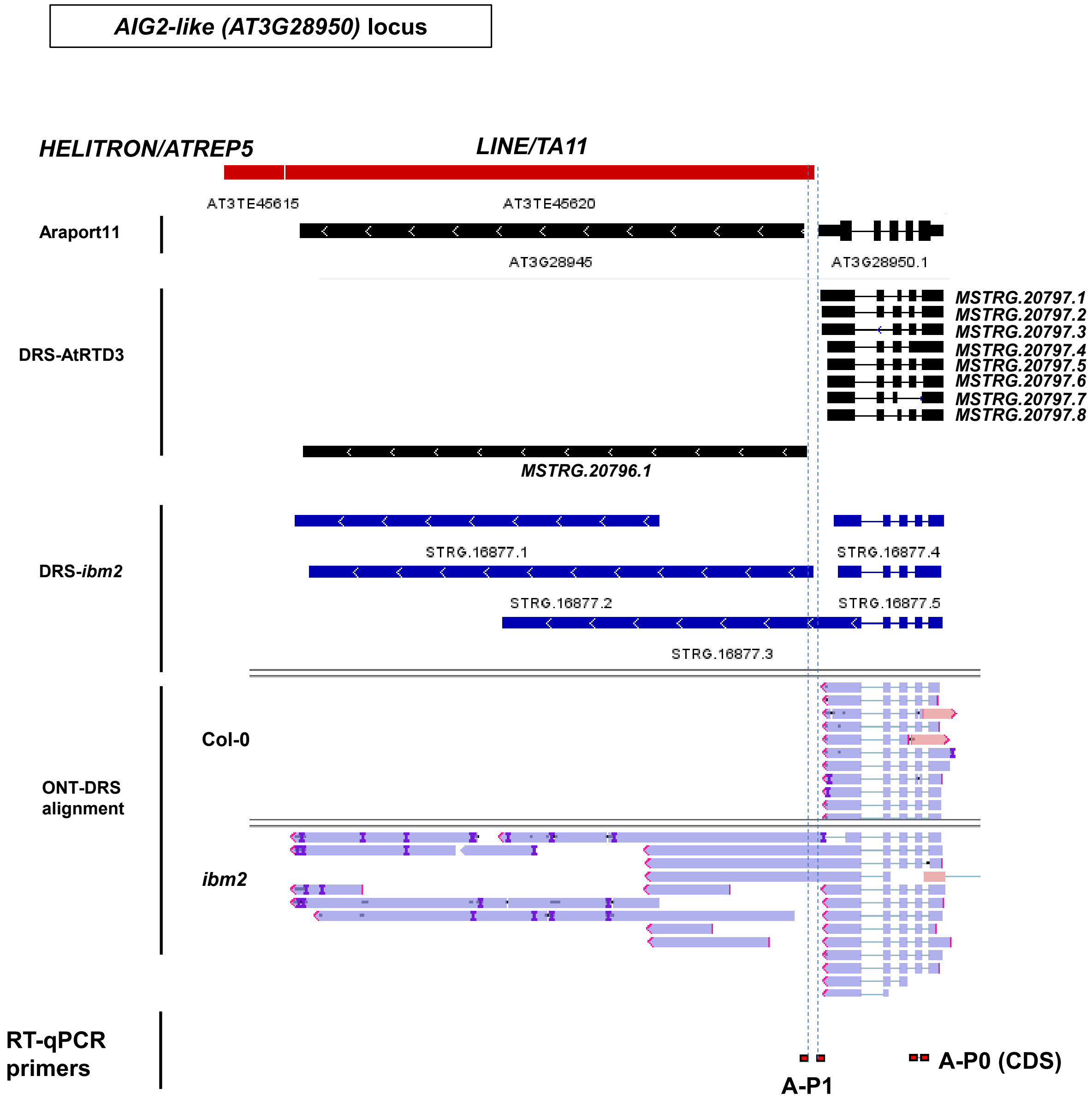


b)

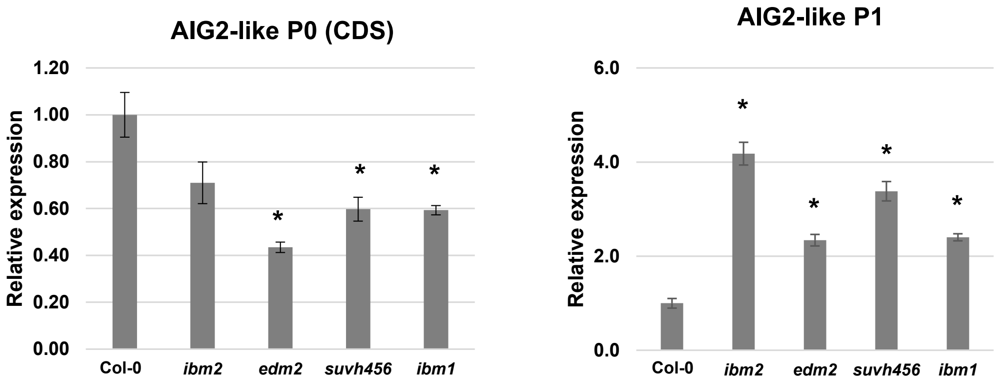


**Supplementary Fig. 11.** *AIG2-like-TA11* (*AT3G28950*) locus. **a)** Genome browser tracks showing the complete DRS-AtRTD3 transcript annotation, DRS reads, and primer positions for RT-qPCR in **b**.
**b)** Relative expression of the *AIG2-like* CDS and altTE-Gis in Col-0 and the epigenetic mutants. Bars represent the means of four biological replicates ± SEM. *, $p < 0.05$ by *t*-test.

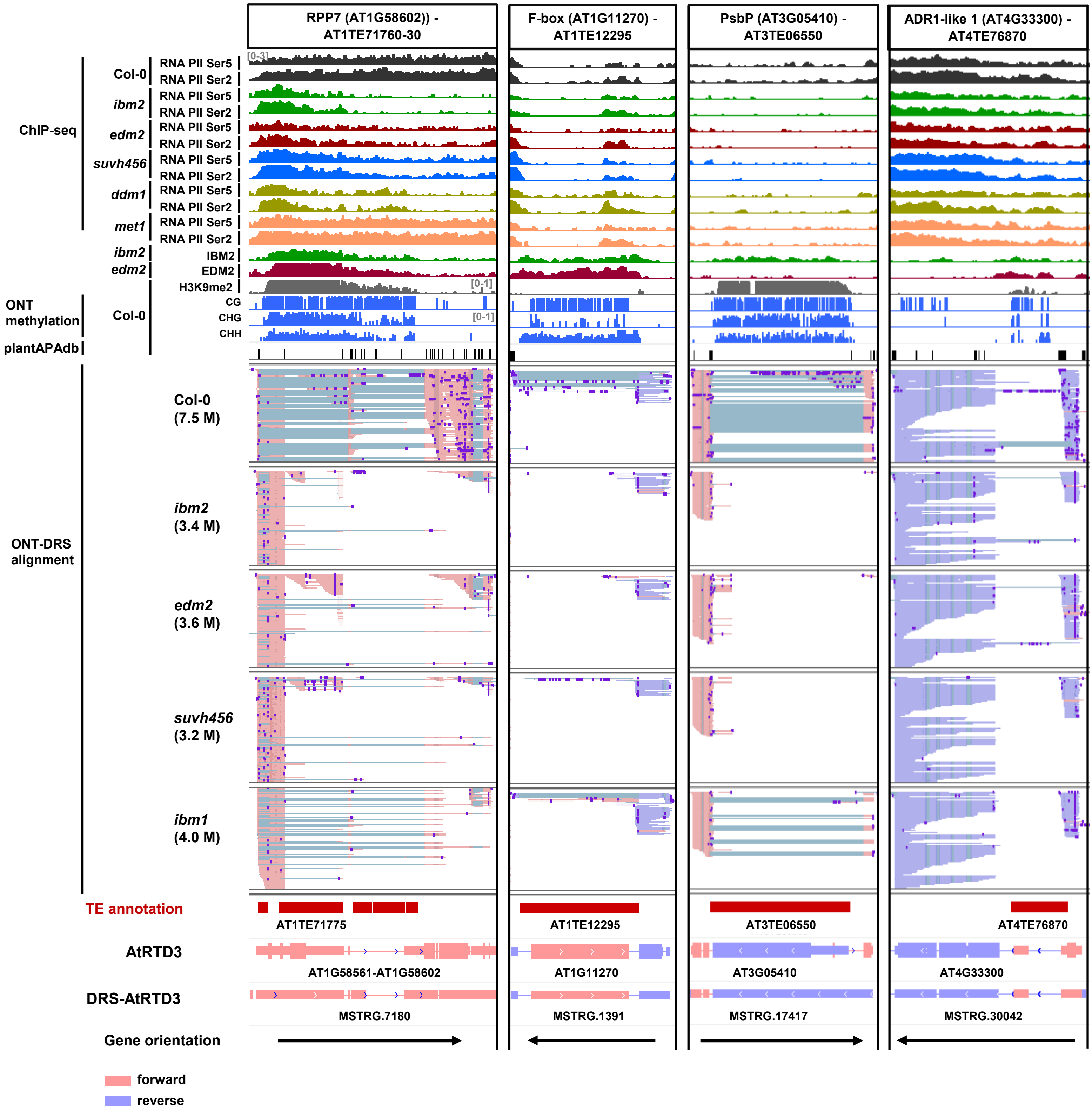


**Supplementary Fig. 12.** Epigenetic regulation of intronic TEs and altTE-Gi production. Representative genome loci showing Epi-altTE-Gi production with TE-ATTS events. Tracks (from top to bottom): ChIP-seq data for RNA Pol II phosphorylated at Ser5/Ser2 in CTD repeats (bins per million); ChIP-seq data for IBM2 and EDM2 localization (bins per million); Col-0 ChIP-seq of H3K9me2 (bins per million); methylation level of Col-0 in CG, CHG, and CHH contexts (0–1); poly(A) sites obtained from the PlantAPA database; DRS read alignments of Col-0 and indicated mutants; TE and transcript annotations of AtRTD3 and DRS-AtRTD3 in this study and the orientation of genes.

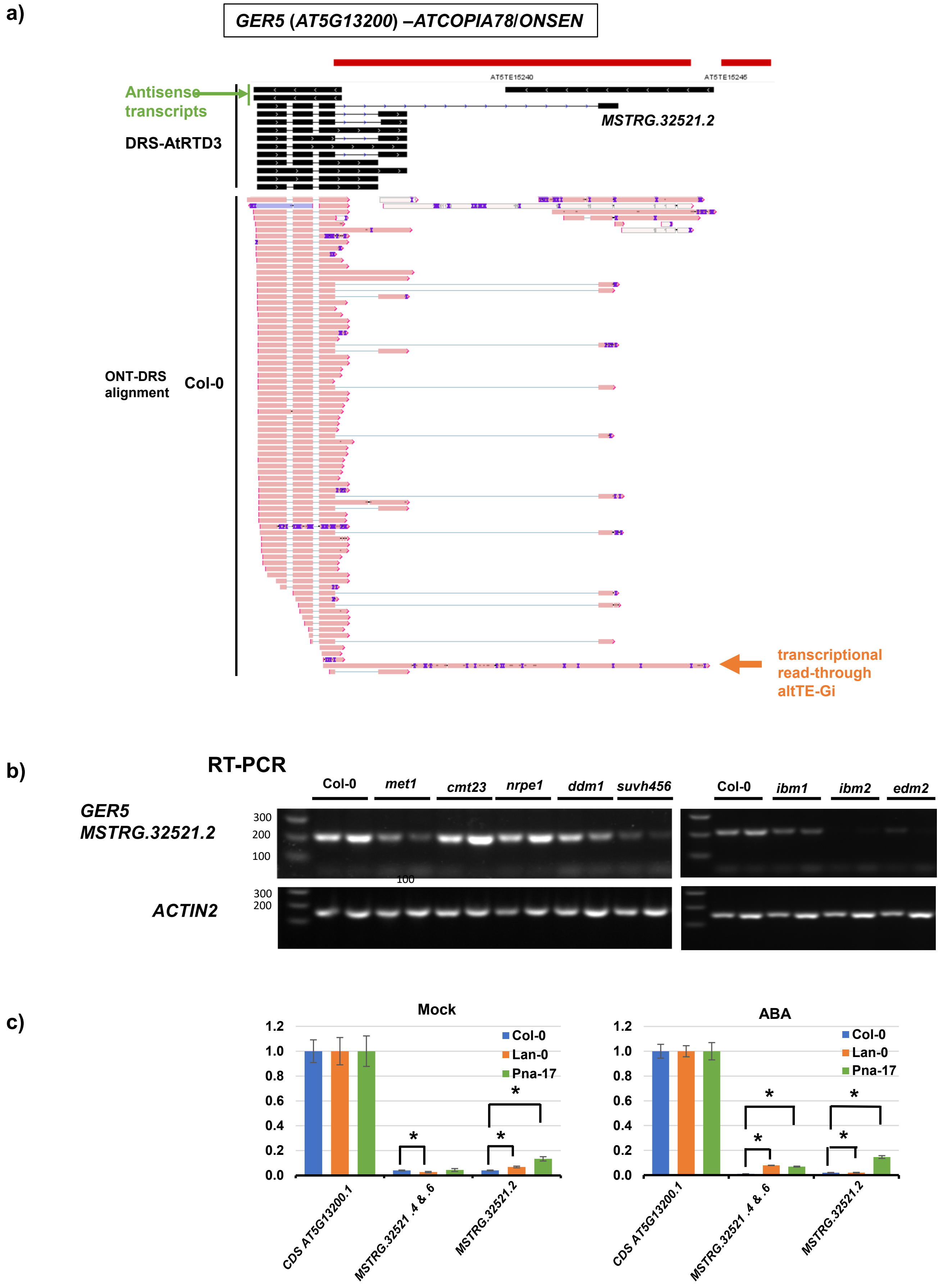


**Supplementary Fig. 13.** The *GER5-ATCOPIA78* (*ONSEN*) (*AT5G13200*) locus. **a)** Genome browser tracks showing the complete DRS-AtRTD3 transcript annotation and Col-0 DRS reads. The green arrow indicates an antisense transcript of *GER5* from the LTR sequence of *ONSEN*. A transcriptional read-through altTE-Gi was also detected in the Col-0-DRS data (orange arrow). **b)** RT-PCR of altTE-Gis (*MSTRG.32521.2*) in Col-0 and the epigenetic mutants. *ACTIN2* (*AT3G18780*) was used as a control. **c)** Relative expression of altTE-Gis (*MSTRG.3521.2*, *4*, *6*) under mock and ABA stress conditions in ecotypes with the *ATCOPIA78/ONSEN* insertion (Col-0, Lan-0, and Pna-17). Relative expression to the transcripts of *GER5* CDS (*AT5G13200.1*) is shown. Bars represent the means of four biological replicates ± SEM. *, $p < 0.05$ by *t*-test.

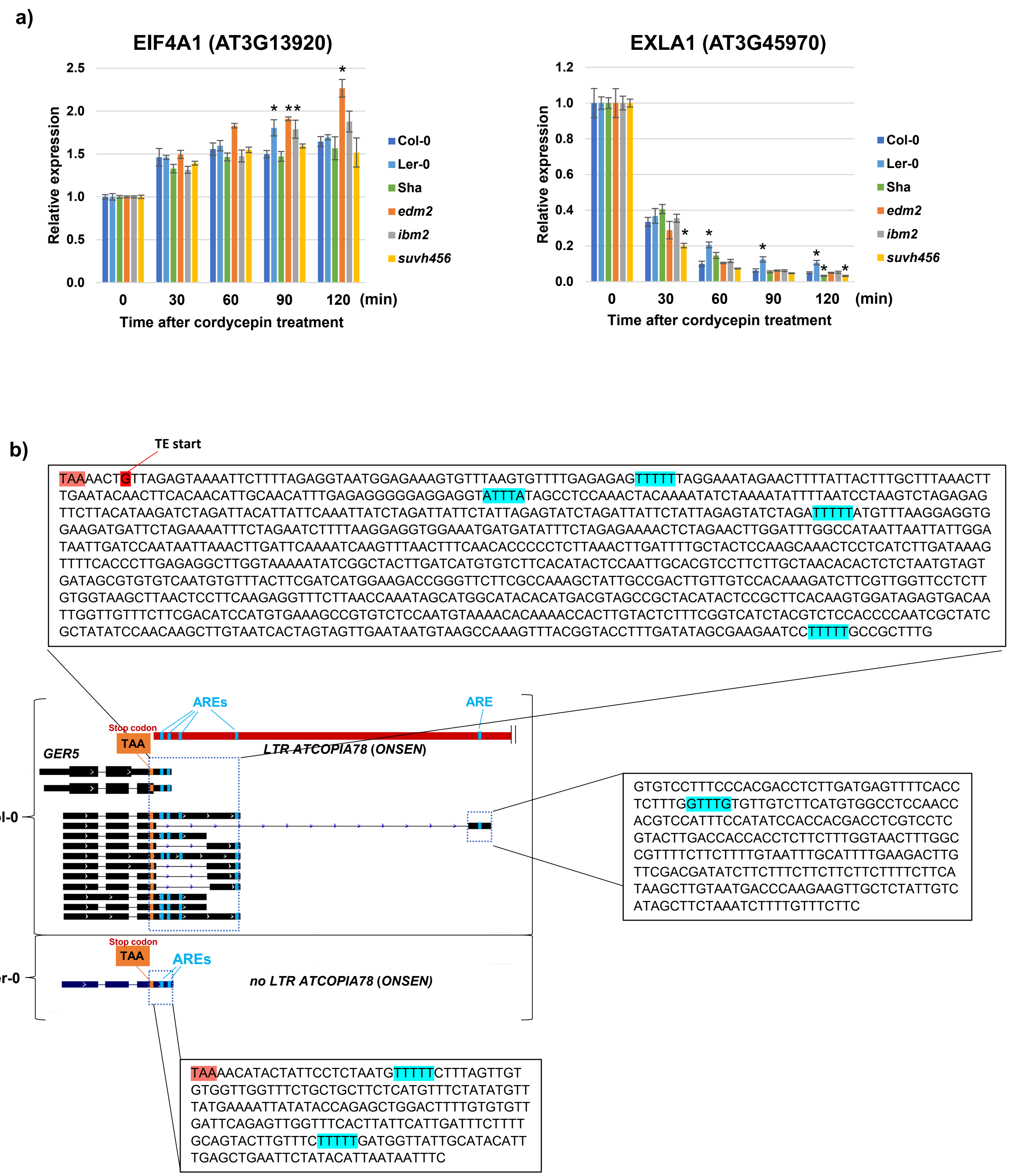


**Supplementary Fig. 14.** RNA stability analysis. **a)** Relative expression of eukaryotic initiation factor-4A (*EIF4A1, AT3G13920*) and EXPANSIN-LIKE A1 (*EXLA1, AT3G45970*) at time 0, 30, 60, 90, and 120 min after cordycepin treatment for Col-0, Ler-0, Sha, *ibm2*, *edm2*, and *suvh456*. Expression levels at 0 min were set as 1. Bars represent the means of four biological replicates ± SEM. *, $p < 0.05$ by *t*-test. *EIF4A1* and *EXLA1* were examined as control transcripts for high and low mRNA stability, respectively. **b)** The *ATCOPIA78/ONSEN* sequence in *GER5* locus provides an AU-rich element (ARE) in altTE-Gi. Ler-0 with no *ATCOPIA78/ONSEN* insertion is also shown.

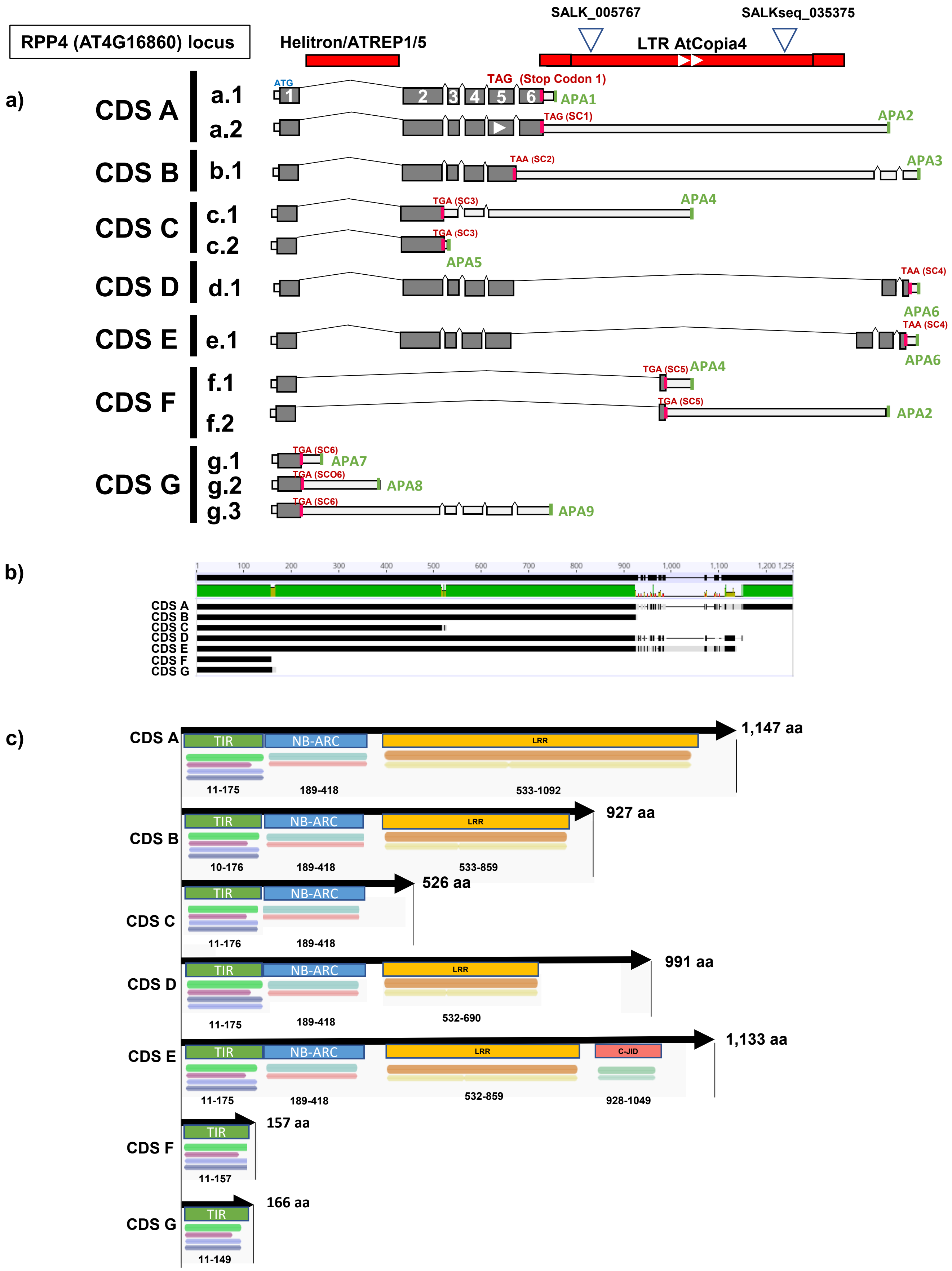


**Supplementary Fig. 15.** Prediction of protein CDS in the *RPP4* locus based on the ONT-DRS transcript data. **a)** Prediction of CDSs encoded by selected *RPP4* transcript isoforms predicted by DRS-AtRTD3 and manually retrieved DRS-ONT read alignments in Supplementary Figure 17. Predicted top codons (SCs) and alternative poly(A) sites (APA) are also represented. **b)** Multiple amino acid alignments of predicted *RPP4* CDS (in the longest isoforms of CDS A–G) show that the *RPP4* locus encodes for at least seven protein isoforms. **c)** Prediction of protein domains in the transcript isoforms (CDS in the longest isoforms of CDS A–G).

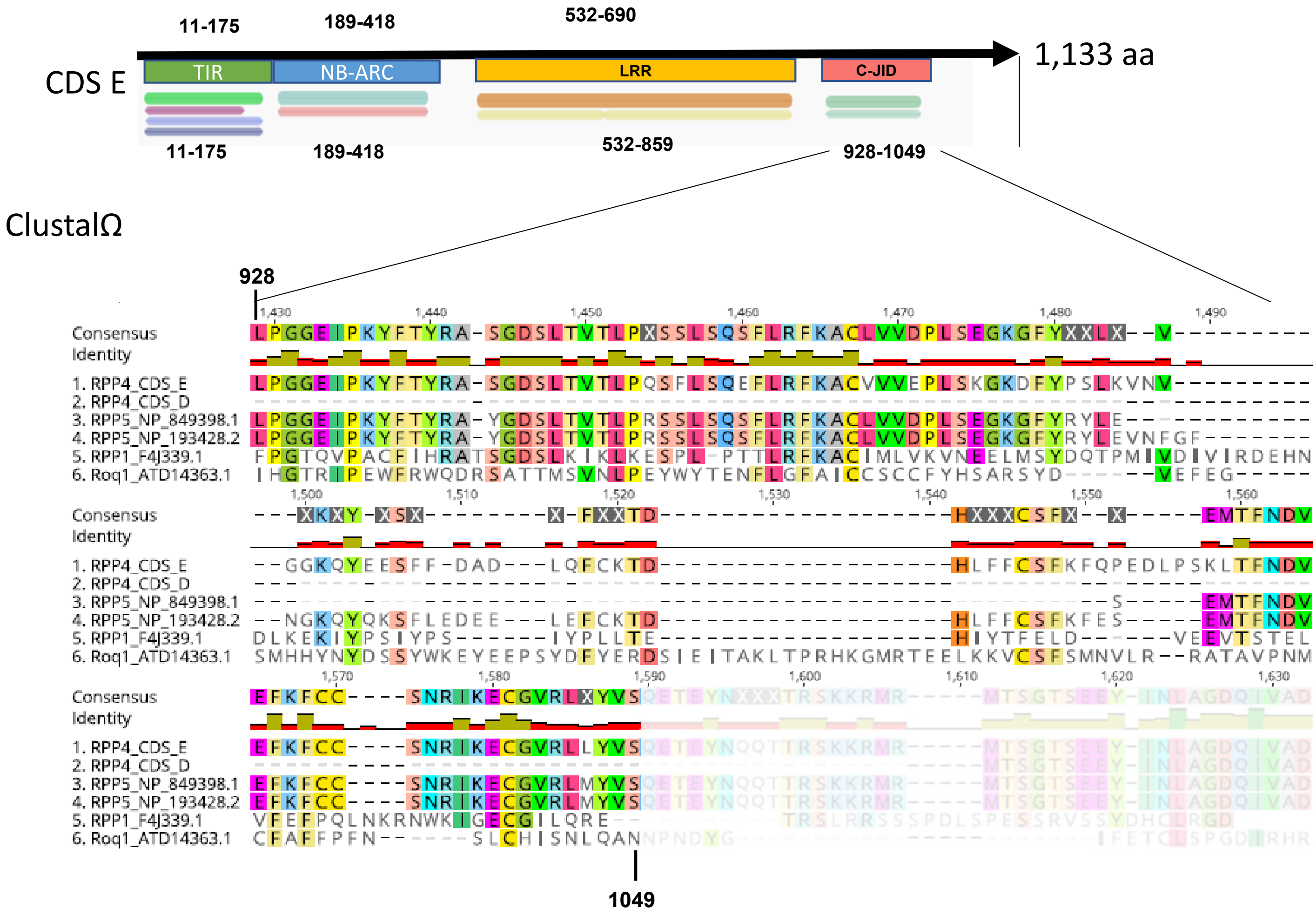


**Supplementary Fig. 16.** *RPP4* CDS "E" in Supplementary Figure 15c is predicted to encode for a C-JID domain in the original protein sequence. Multiple alignments of the amino acids predicted to be encoded by "RPP4 CDS E", "RPP4 CDS D" (no C-JID predicted), RPP5 proteins variants (ID: NP_849398.1 and NP_193428.2), RPP1 (ID: F4J339.1), and Roq1 (ID: ATD14363.1) at the predicted C-JID domain region.

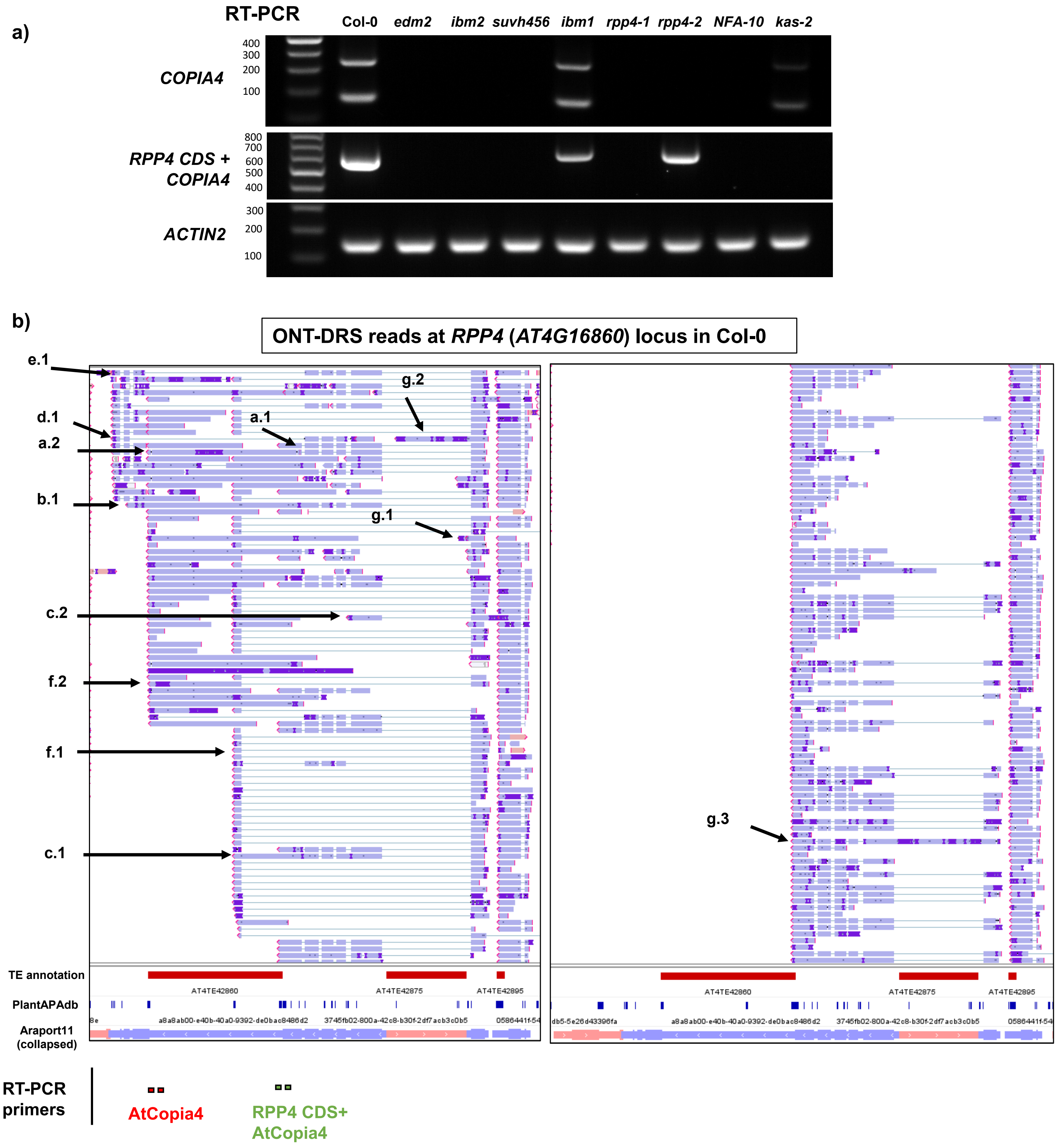


**Supplementary Fig. 17.** Detection of altTE-Gis at the *RPP4* locus. **a)** Agarose gel electrophoresis of RT-PCR amplicons with primers targeting the *ATCOPIA4* or *RPP4-ATCOPIA4* transcribed region. (see Supplementary Fig. 17b for primer positions). *ACT2* (*AT3G18780*) was used as a positive control. **b)** Genome browser tracks showing the alignment of ONT-DRS reads at the *RPP4* locus. Arrows indicate several transcripts classified in the groups A–G shown in Supplementary Fig. 15, which were predicted to encode different protein isoforms. The positions of primers used in Supplementary Figure 17a are illustrated at the bottom.

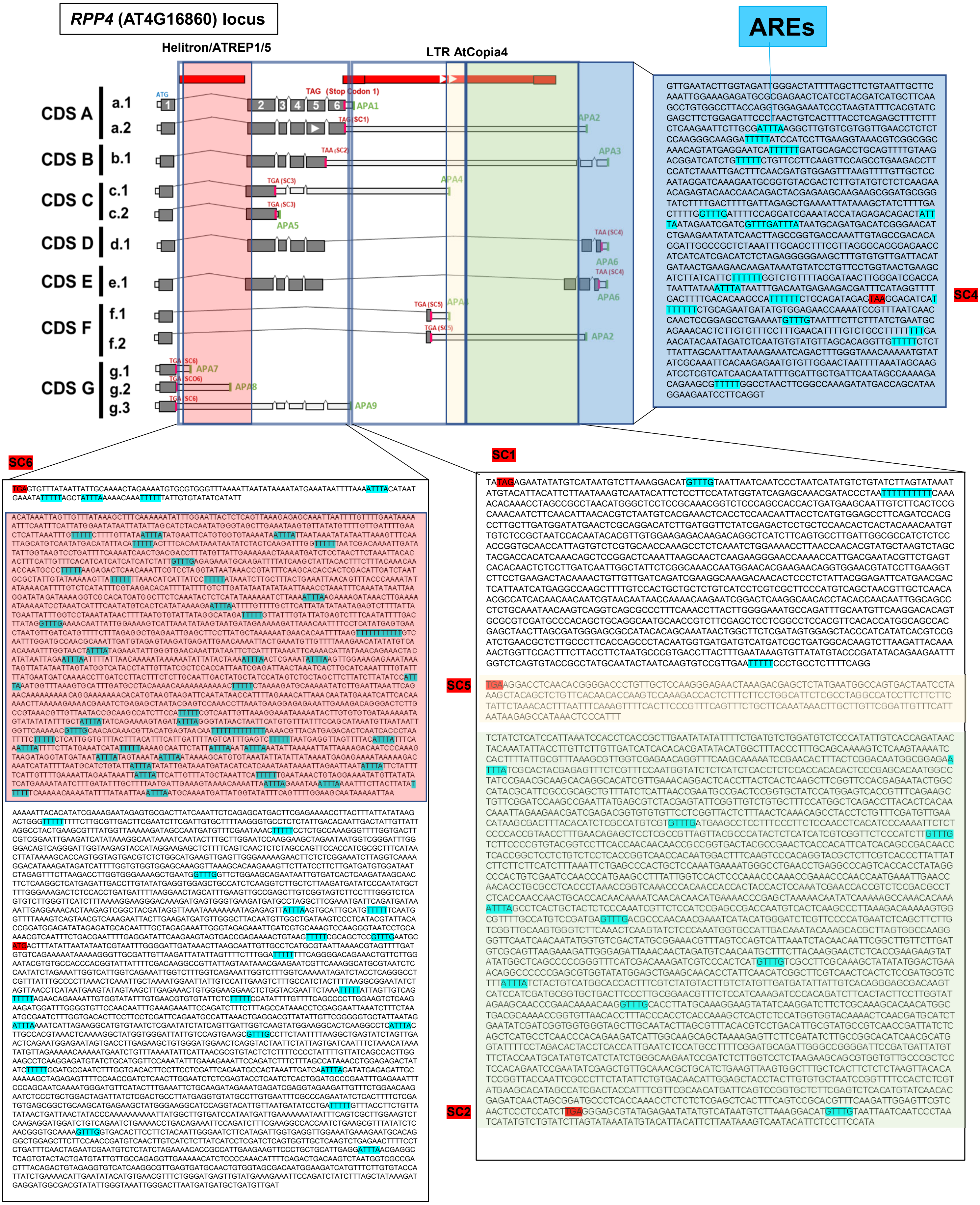


**Supplementary Fig. 18.** The 3′-UTR of *RPP4* transcripts are predicted to contain ARE motifs. Prediction of potential ARE motifs in the *RPP4* transcripts. SC, predicted stop codons (Supplementary Fig. 15).

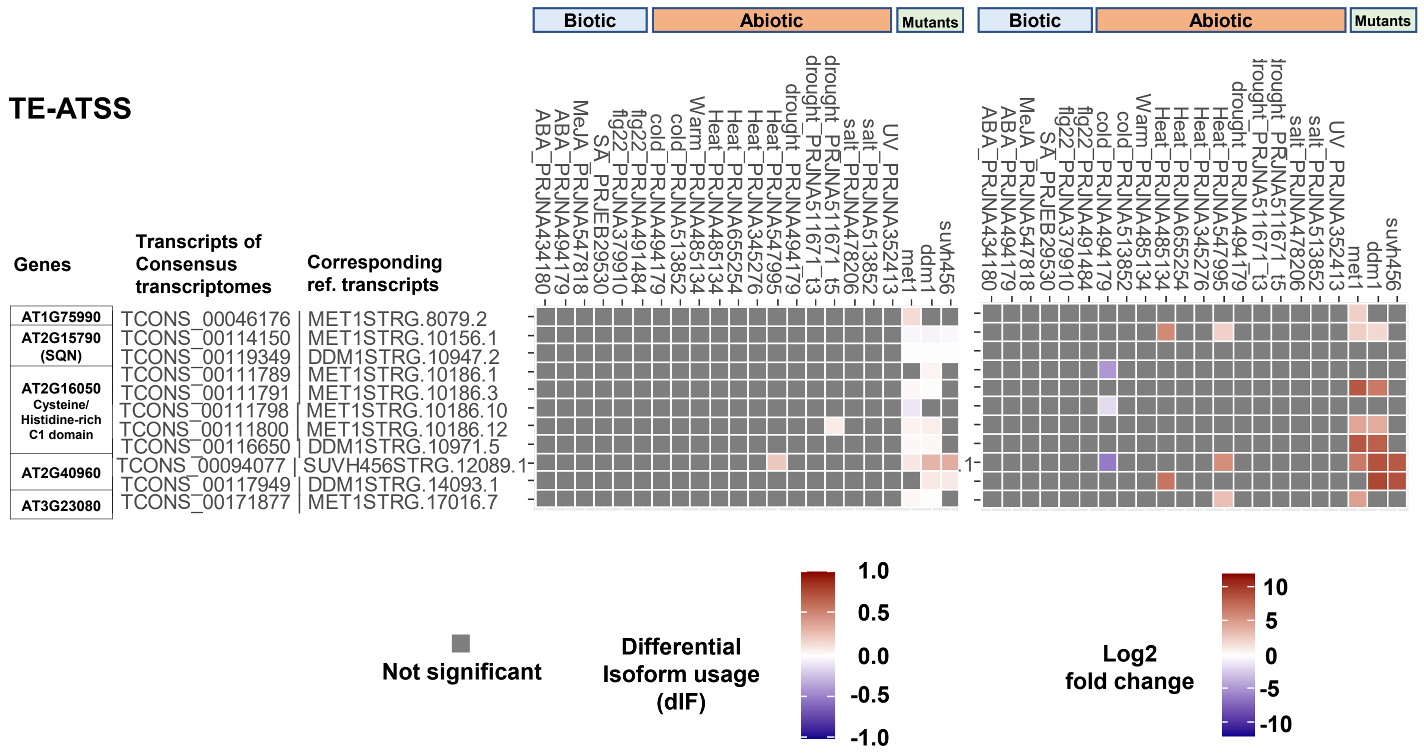


**Supplementary Fig. 19**. Regulation of Epi-altTE-Gi candidates with TE-ATSS under various stress conditions. Heatmaps show the statistically significant dIF (left) and differential expression fold-change (right) under stress conditions or in the epigenetic mutants. Transcriptome data with stress treatment studies are from the public RNA-seq data.

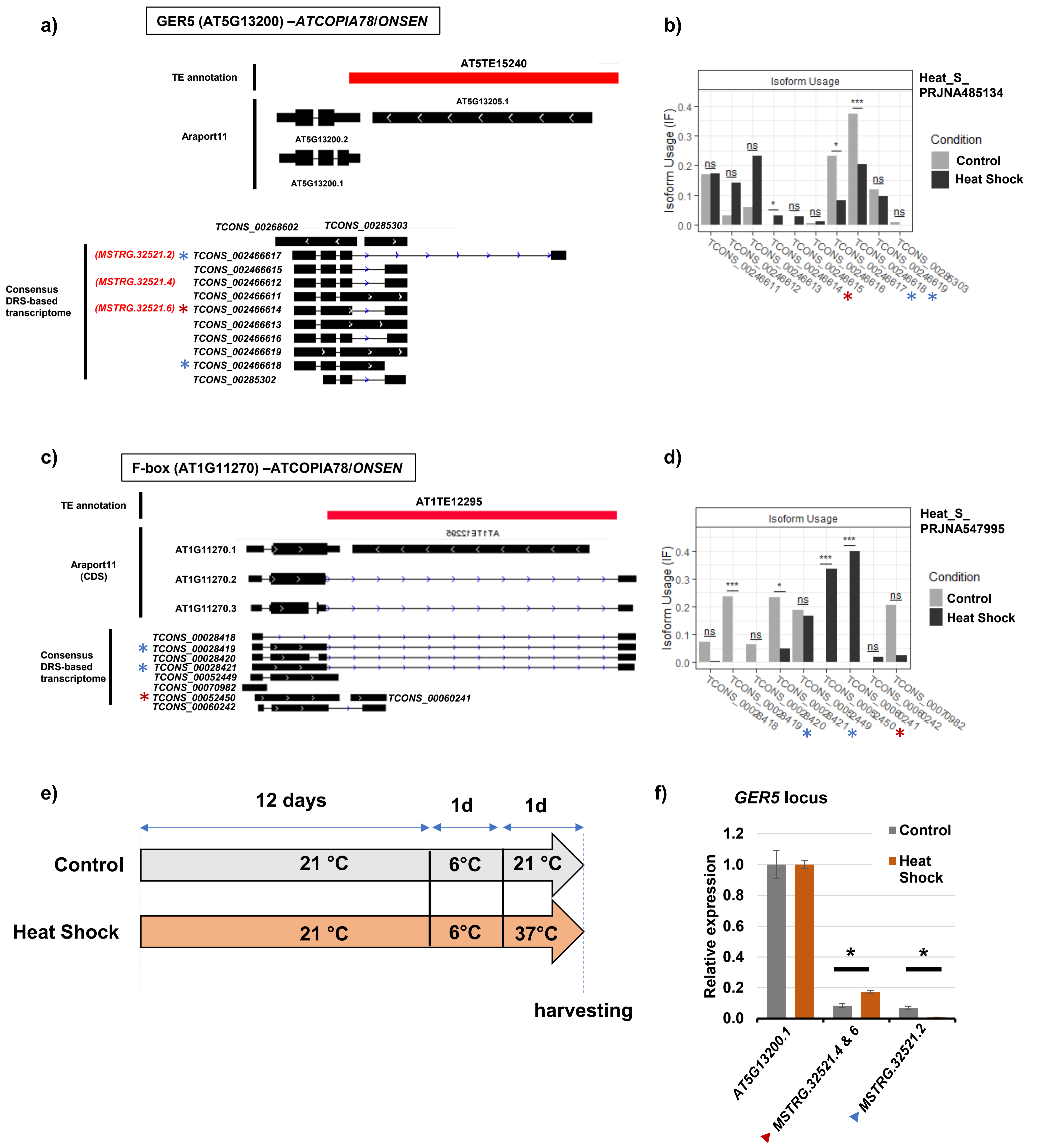


**Supplementary Fig. 20**. Isoform switching events of *ONSEN*-containing genes with heat shock treatment. **a)** *GER5* (*AT5G13200*) transcripts in Araport11 and the consensus DRS transcriptome. **b)** Isoform switching of altTE-Gis associated with the *GER5* locus were detected in the public RNA-seq dataset (PRJNA485134) under heat shock conditions. *, $q < 0.05$; ***, $q < 0.001$ (FDR corrected *p*-values) found by IsoformSwitchAnalyseR. altTE-Gis with red and blue asterisks correspond to transcripts in DRS-AtRTD3 marked in **a**, which showed significantly higher and lower usage, respectively. **c)** F-box gene (*AT1G11270*) transcripts in Araport11 and the consensus DRS transcriptome. **d)** Isoform switching of altTE-Gis associated with the *GER5* locus detected in the public RNA-seq dataset (PRJNA547995) under heat shock conditions found by IsoformSwitchAnalyseR, with *, $q < 0.05$ and ***, $q < 0.001$ (FDR corrected *p*-values). altTE-Gis with red and blue asterisks correspond to transcripts in DRS-AtRTD3 marked in **c**, which showed significantly higher and lower usage, respectively. **e)** Experimental scheme of heat shock treatment. **f)** Relative expression of the *GER5* transcripts (*AT5G13200.1*, *MSTRG.32521.4, 6*, and *MSTRG.32521.2;* see Figure 5a). Bars represent means of four biological replicates ± SEM and *, $p < 0.05$ by *t*-test.

Supplementary Table 1: Number of corrected “passed reads” and aligned reads (Chr1 to Chr5).

| Genotypes | Total corrected reads | Total corrected mapped reads |
|---|---|---|
| col-0 (wt) concatenated with ONT DRS reads from Parker et al 2020 | 8,336,887 | 7,547,469 |
| *met1* | 4,134,823 | 3,974,357 |
| *ddm1* | 5,236,774 | 5,145,090 |
| *suvh456* | 3,378,954 | 3,250,179 |
| *ibm1* | 4,136,991 | 4,030,594 |
| *ibm2* | 3,484,091 | 3,436,936 |
| *edm2* | 3,727,236 | 3,678,813 |

Supplementary Table 2: Number of genes and transcripts in DRS-AtRTD3 and DRS-Araport11 compared to original AtRTD3 and Araport11 (Chr1 to Chr5).

| Annotation | Number of genes | Number of transcripts |
|---|---|---|
| AtRTD3 | 40,570 | 168,949 |
| DRS-AtRTD3 | 39,998 | 199,489 |
| Araport11 | 27,655 | 48,149 |
| DRS-Araport11 | 30,552 | 94,765 |

Supplementary Table 3: Number and percentage of genes and TEs associated to TE-Gt and altTE-Gi found by ParasiTE in AtRTD3, DRS-AtRTD3, Araport11 and DRS-Araport11.

| Gene annotation | AtRTD3 | DRS-AtRTD3 | Araport11 | DRS-Araport11 |
|---|---|---|---|---|
| TE annotation | TAIR10 TE annotation (length ≥200 bp) (18,881) | | | |
| Gene associated to TE-Gt | 2,584 (6.4%) | 3,030 (7.6%) | 2,358 (8.5%) | 2,931 (9.6) |
| Gene associated to altTE-Gi | 1,261 (3.1%) | 1,171 (2.9%) | 705 (2.5%) | 1,082 (3.5%) |
| TEs associated to TE-Gt | 2,734 (14.5%) | 3,194 (16.9%) | 2,511 (13.3%) | 2,989 (15.8%) |
| TEs associated to altTE-Gi | 1,324 (7.0%) | 1,296 (6.9%) | 753 (4.0%) | 1,171 (6.2%) |

Supplementary Table 4: List and detailed of published RNAseq data (paired end) used to investigate the effect of stress conditions on Epi-altTE-Gi expression and isoform switching events.

| Stress conditions | SRA project | Experiments |
|---|---|---|
| ABA (50 μM) 6h | PRJNA434180 | Twelve-day-old seedlings were treated with DMSO control or 50 μM ABA for 6 h. |
| ABA (100 μM) 4h | PRJNA494179 | 2-week-old seedlings were transferred to 1/2 MS liquid medium with 100 μM ABA for 4h. |
| MeJA (200 μM) 4h | PRJNA547818 | Three-week-old seedlings were grown on MS medium and were treated with mock or 200 μM MeJA for 4 h. |
| SA (0.5 mM) 24h | PRJEB29530 | 5-weeks old plants were then sprayed with water or 0.5 mM SA, and harvested after 24 hours. |
| Flg22 (1 μM) 1h | PRJNA491484 | 14 days-old seedlings were grown under short-day conditions (8h light/ 16 h dark) at 22°C and treated with 1μM of flg22 for 1h. |
| Flg22 (1 μM) 0.5h | PRJNA379910 | 2-week-old seedlings were treated with deionized water (mock) or with a final concentration of 1 μM flg22 for 30 min. |
| Salt (NaCl) | PRJNA478206 | 2-week-old seedlings were watered with Hoagland's solution with added 50 mM NaCl. Three days later, they were watered again with Hoagland's with added 100 mM NaCl. The flats were then watered with 150 mM NaCl Hoagland's solution every three days for a total of four additional treatments. Control flats were watered at the same time as salt-treated plants, with solely Hoagland's solution. After treatments, rosette leaves were collected. |
| Salt NaCl (150 mM) 24h | PRJNA513852 | 2-week-old seedlings were transferred to Petri dishes containing half MS medium with 1% (w/v) sucrose supplemented with 150 mM NaCl for 24 h. |
| Warm (28/23 °C) 7d | PRJNA485134 | Plants at 23 days after sowing were subjected to 28/23 °C (day/night) for 7 days as the prolonged warming treatment. Leaves were sampled from plants at 30 days after sowing (at the rosette growth stage). |
| Heat (38 °C) 6h | PRJNA485134 | Plants were exposed to 38 °C for 6 h during the day portion of the photoperiod as the heat shock treatment. Leaves were sampled from plants at 30 days after sowing (at the rosette growth stage). |
| Heat shock (37°C) 1h | PRJNA345276 | 7-d-old seedlings under the normal condition at 22°C (control condition) after an HS treatment at 37°C for 1 h. |
| Heat shock (37°C) 3x12h | PRJNA547995 | 7-day-old seedlings were placed in an incubator set at alternating temperatures of 37 and 22 °C, 12 h each, for 3 days with a 12 h photoperiod. Control were at 22 °C. |
| Heat shock (42 °C) 5h | PRJNA655254 | Plants grown in soil under 16 h light/8 h dark conditions for 3 weeks were used for heat treatment at 42 °C for 5 h. |
| Cold (4 °C) 20 days | PRJNA494179 | 2-week-old seedlings were transferred to 4 °C and cultured under short-days conditions. Harvested after 20 days. |
| Cold (4 °C) 1day | PRJNA513852 | Seedlings grown under long days at 20°C and transferred to 4°C for 24 h |
| Drought 8h | PRJNA494179 | 2-week-old seedlings were removed from the agar and desiccated in dishes. Harvested after 8 hours. |
| Drought ADT3 (30–35% moisture) | PRJNA511671 | Gradient drought treatments were administered 21 days after planting plants. The soil moistures for Arabidopsis ADT1 plants were 50–55%, 40–45% for ADT2, 30–35% for ADT3, 20–25% for ADT4, 10–15% for ADT5. |
| Drought ADT5 (10–15% moisture) | PRJNA511671 | |
| UV-B 24h | PRJNA352413 | 15-day-old seedlings were exposed to UV-B light (0,210 mW/cm2) during 24 h and then recovered for 72 h under controlled conditions (16 h light, 100 μmoles m−2 s−1, 22 ± 2ºC). As control, seedlings covered with a cellulose acetate polyester filter were used. |

Supplementary Table 5: RT-qPCR Primers used in this study.

| Name | Primer sequence 5` → 3` | Primer efficiency |
|---|---|---|
| BEST1_cds_P0_F | CTTTCCCTGTTGCTCTTAAGTGTC | 104.3 |
| BEST1_cds_ P0_R | CAAGTTCCGGAGATCTCTTGCT | |
| BEST1_ P1_F | GGGAGCTAAGAACATGTGATGG | |
| BEST1_ P1_R | CACTTAAACACTTTCTCCATTACCTC | |
| AT4G16900_ cds_ P0_F | CGTACCTCACGTCTATAGCATGAAG | |
| AT4G16900_ cds_ P0_R | ACTTCCCTCCTATCTTTATATCCTTACC | |
| AT4G16900_ P1_F | CGTTACAGATAACCTGCCCTACAG | |
| AT4G16900_ P1_R | GAGTATACTCGTCATCGTTTCTGG | |
| AT4G16900_ P2_F | CTGGTAAGGACATAGGCTTGAAC | |
| AT4G16900_ P2_R | CCTTTCATTATCTCTTCCAGAACTG | |
| GER5_cds_AT5G13200.1&2_F | CTCTGGCACAATCTGAAGACAG | |
| GER5_cds_AT5G13200.1&2_R | GCTCTGTTCCGAAAATCTGTCTG | |
| GER5_cds_AT5G13200.1_F | GGAACTGTCTATCTTTCGAATGC | 104.5 |
| GER5_cds_ AT5G13200.1_R | GGTACAACCACCCTGTAGTAGCTC | |
| GER5_ MSTRG.32521.4&.6_F | CATCTGCTGACCAGTGTCTCC | |
| GER5_ MSTRG.32521.4&.6_R | ACATTGACACACGCTATCACTCTA | |
| GER5_ MSTRG.32521.2_F | CATCTGCTGACCAGTGTCTCC | |
| GER5_MSTRG.32521.2_R | CGTGGGAAAGGACACCTCTA | |
| ACT2_F | GATGGAAACCTCAAAGACCAG | 102.0 |
| ACT2_R | CACAAACGAGGGCTGGAAC | |
| GAPDH_(AT1G13440)_F | CGTTGACCTTATCGTTCACATG | 102.0 |
| GAPDH_(AT1G13440)_R | CAGAACAGTACGAACTCAACCAC | |
| RPP4_cds_ P0_F | CCACGGTATTATTTTCGACAAG | 100.4 |
| RPP4_cds_ P0_ R | GGTAGATCTATTTTTGACCAAAGACC | |
| RPP4_ P1_F | CAAACATGTCCTTTAAGACATTATGAC | |
| RPP4_ P1_R | CGACGTATATTGGGTAAATTGG | |
| RPP4_ P2_F | CTTGAAGCATGATCGTAGGATG | |
| RPP4_ P2_R | CCATAGTTGAATACTTGGTAGATTGG | |
| RPP4_ P3_F | GATGTCATCTGGATGCCTTCC | |
| RPP4_ P3_R | GTGAATTTCGCCCAGAATATCTC | |
| RPP4_ P4_F | GAGGTCCTTCACCCGTTCC | |
| RPP4_ P4_R | CGAGGACAAACCAGAGGATC | |
| RPP4_ P5_F | CACGCACATTTTCTAGTTTTGC | |
| RPP4_ P5_R | GATCTTCGGAACGGGTATG | |
| AIG2-like_(AT3G28950)_cds_P0_F | GGTCTTAAGCCCAAAAGAGGG | |
| AIG2-like_(AT3G28950)_cds_P0_R | GCAATAGTTGCTTCGGAGTGG | |
| AIG2-like_(AT3G28950)_P1_F | GGTCTTAAGCCCAAAAGAGGG | |
| AIG2-like_(AT3G28950)_P1_R | GCAATAGTTGCTTCGGAGTGG | |

Supplementary Table 6: RT-PCR Primers used in this study.

| Name | Primer sequence 5` → 3` |
| --- | --- |
| ACT2_RT-PCR_F | AACTCTCCCGCTATGTATGTCGCCATCCAA |
| ACT2_RT-PCR_R | AGCAAGGTCAAGACGGAGGATGGCATGAGG |
| GER5_RT-PCR_F | CATCTGCTGACCAGTGTCTCC |
| GER5_RT-PCR_R | CCAAAGAAGAGGTGGTGGTC |
| BEST1_RT-PCR_P2_F | GGGAGCTAA GAACATGTGATGG |
| BEST1_RT-PCR_P2_R | GTTAGACTAATCATCTCACTAGC |
| ATCOPIA4_RT-PCR_F | TTGACGCCCAACAACGAAAT |
| ATCOPIA4_RT-PCR_R | GTTTCCGCATAGTCGACACC |
| RPP4_CDS_ATCOPIA4_F | CGATAGAACCATCAAGATGTCCT |
| RPP4_CDS_ATCOPIA4_R | GAGTGATGCAACTGTGGTAGC |
